# Perturbation-induced gene programs map the hidden regulatory architecture of complex diseases

**DOI:** 10.64898/2026.11.03.762080

**Authors:** Berk Turhan, Hanbyul Lee, Dingyu Liu, Ann Le, Arpan Ghosh, Dhruba Nandi, Rahul Mazumder, Jesse M. Engreitz, Xin Jin, Danwei Huangfu, Kushal K. Dey

## Abstract

Steady-state molecular profiles incompletely capture the regulatory programs through which genetic variation influences complex disease, particularly programs that emerge only when cells are perturbed. Here, we introduce Perturbscape, a computational framework that learns multiscale gene programs for individual perturbations in a CRISPR Perturb-seq screen using linear and nonlinear contrastive approaches alongside complementary program-learning methods, and integrates them with human genetics to prioritize disease-relevant genes, perturbations and downstream regulatory pathways. Across 15,859 perturbation-by-context profiles from 7 Perturb-seq systems and 53 complex diseases and traits (mean GWAS N=291K), perturbation-induced programs showed up to 4.3-fold higher heritability informativeness than analogous steady-state multi-tissue gene prioritization, and retained significant conditional heritability informativeness across all 34 matched cell system–trait comparisons after accounting for steady-state prioritization. Preserving perturbation-specific programs further yielded 2.5–5.3-fold higher trait relevance scores than programs derived from control cells, pooled perturbations or differential-expression summaries. In K562, Perturbscape-prioritized genes showed stronger enrichment for rare-variant burden genes than steady-state prioritization and were supported by orthogonal enhancer CRISPRi screens. In HepG2, LDL cholesterol-associated perturbation– induced programs specifically revealed statin-responsive pathways not significantly enriched by steady-state PoPS prioritization, supported by pharmacological transcriptional responses and population plasma proteomics. Perturbscape also recovered blood-related trait-conditioned trans-regulatory relationships that were specifically supported by population trans-eQTLs in IFN-γ-stimulated monocytes (OR = 1.30, P=1.6e-05). Perturbscape further revealed developmental-stage-specific convergence between monogenic diabetes regulators and polygenic T2D risk during pancreatic islet differentiation, and neuronal-subclass-specific autism-associated regulatory programs *in vivo*. Together, these findings establish perturbation-induced gene programs as a complementary layer of disease-relevant regulatory information, revealing genetically informative pathways and context-specific regulatory relationships incompletely represented by steady-state molecular atlases.

## Introduction

Genome-wide association studies (GWAS) have linked tens of thousands of loci to complex human diseases and traits, but identifying the causal genes and regulatory programs that mediate this genetic risk remains a central bottleneck in human genetics^1–4^. Steady-state molecular annotations do not effectively capture regulatory programs that emerge only following a gene perturbation or within specific cellular, developmental or activation contexts^5,6^. CRISPR Perturb-seq screens now measure, directly and at scale, the downstream transcriptional consequences of experimentally perturbing individual genes^7–9^, revealing these context-specific trans-regulatory programs. Recent studies have begun to connect such programs to disease, relating the trans-regulatory targets of GWAS-implicated genes to specific disease pathways^10,11^ and prioritizing perturbed genes through the heritability enrichment of their downstream differentially expressed genes^12–14^. Recent work has further integrated Perturb-seq regulatory effects with gene-level loss-of-function burden associations to construct regulator–program– trait models for blood traits, illustrating the potential of perturbation-derived regulatory structure for interpreting human genetic effects^15^. However, existing approaches have generally examined a limited range of transcriptional programs and cellular systems or relied on well-powered quantitative rare-variant associations for regulatory modeling.

For a disease-relevant perturbation, we expect its *trans*-regulatory gene programs to preferentially contain genes enriched for *cis*-genetic disease signal at their linked loci^13,15^. Building on this principle, we present Perturbscape, an end-to-end computational framework that operationalizes this cis–trans convergence to connect a perturbation from a CRISPR Perturb-seq screen to the genetic architecture of a complex disease. From a single screen, Perturbscape constructs a multiscale atlas of gene programs for each perturbation using several linear and nonlinear contrastive learning approaches alongside complementary canonical and local gene-program learners. These programs capture global and locally coordinated transcriptional responses, including regulatory variation selectively enriched following perturbation relative to control cells. Perturbscape integrates these programs into optimized disease-specific weighted representations, termed *meta-programs* (denoted MetaProg ^(*p*)^ for perturbation *p*), and examines the relevance of these meta-programs to disease heritability^16–18^. Complementary to approaches that prioritize genes using precompiled steady-state expression, pathway or protein network annotations^19,20^, Perturbscape derives its features directly from perturbation atlases that sample experimentally induced regulatory states and identifies those most relevant to matched disease genetics.

Perturbscape provides two primary disease-genetic readouts from each perturbation screen: (i) atlas-wide prioritization of trait-relevant genes using an atlas-wide meta-program (denoted MetaProg^(ALL)^) constructed by combining all perturbation-induced programs; and (ii) perturbation-level trait relevance scores (denoted TRS^(*p*)^ for perturbation *p*) that quantify the heritability relevance of individual trans-regulatory responses, together with perturbation–pathway–disease maps. We applied Perturbscape to 15,859 perturbation-by-context pairs across 7 Perturb-seq systems, spanning diverse cell lines, a stem-cell differentiation time course and an *in vivo* brain screen, and evaluated against 53 matched diseases and quantitative traits (mean GWAS *N* = 290,733). Across these systems, atlas-wide perturbation meta-programs prioritized disease genes that were complementary to, and often more genetically informative than, those prioritized from steady-state multi-tissue resources. Perturbscape further revealed convergence between common- and rare-variant disease associations within perturbation-responsive gene programs and identified disease-relevant regulatory relationships supported by orthogonal enhancer perturbation experiments. Perturbation-specific meta-programs further revealed LDL cholesterol-associated statin-responsive pathways not significantly enriched among genes prioritized by PoPS, and recovered trans-regulatory relationships supported by population trans-eQTLs in an activation-state-specific manner. Across developmental and *in vivo* perturbation systems, Perturbscape resolved the emergence of monogenic–polygenic diabetes convergence during pancreatic differentiation and neuronal-subclass-specific autism-associated regulatory programs. Together, these analyses establish perturbation-induced gene programs as a complementary representation of complex disease regulatory architecture beyond steady-state molecular atlases.

## Results

### Overview of the Perturbscape framework

The central premise of Perturbscape is that disease-relevant perturbations preferentially induce trans-regulatory responses in which the downstream genes are enriched for cis-genetic disease signal at their linked loci^12,13^ (**Figure 1a**). Perturbscape first constructs a multi-resolution library of gene programs corresponding to each perturbation in the screen, rather than relying on a single differential-expression signature or latent representation (**Figure 1b**). For each perturbation, we derive candidate programs using complementary differential expression and co-expression methods (DE, DGCA^21^), canonical latent factor methods (PCA, cNMF^22^), contrastive methods (contrastive PCA, kernel contrastive PCA, and ContrastiveVI^23,24^), and a graph-based gene-module detection method (Hotspot^25^) (**Figure 1b**; **Methods**; **Table S1**). A distinctive component of this library is a family of contrastive PC learners that includes the ContraPC and kernel ContraPC formulations introduced here, designed to isolate transcriptional variation selectively enriched by a focal perturbation relative to control cells corresponding to non-targeting and safe-targeting control guides (**Figure 1b**). Let *ϕ*(*x*) denote a representation of a cell *x*, and let Σ*_p_* and Σ_0_ denote the covariance operators of the perturbation and control populations, respectively, in this representation space. We formulate these learners as

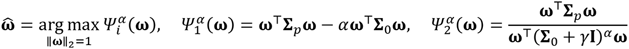

**Figure 1:**
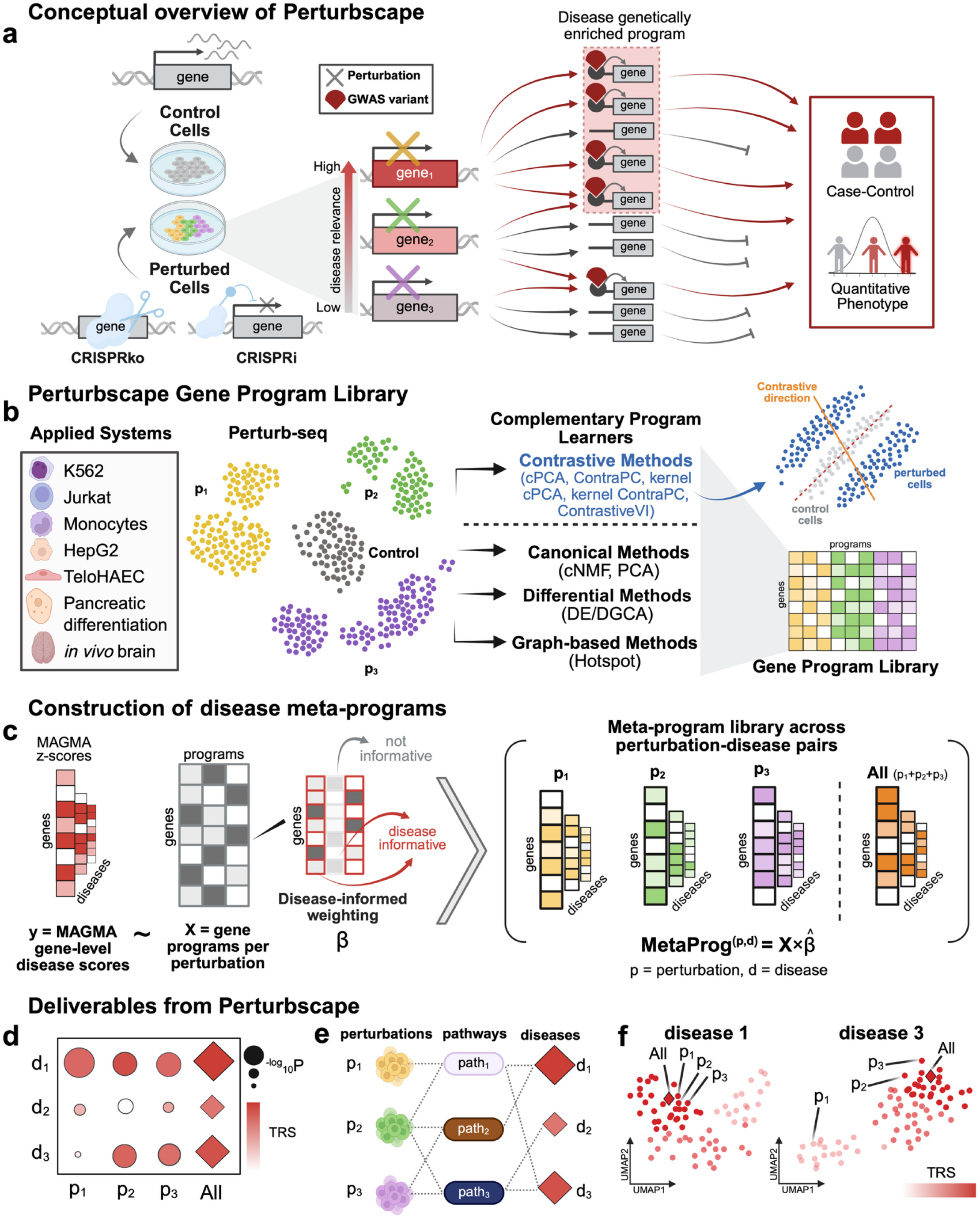
Perturbscape framework for mapping perturbation-induced gene programs to human disease genetics. (a) Perturbscape prioritizes CRISPR perturbations by mapping gene programs underlying their induced transcriptional responses relative to control cells and quantifying their enrichment for inherited disease-related genetic risk. This links experimentally induced cellular responses to the genetic architecture of human complex diseases and traits. (b) Construction of a comprehensive perturbation-response program library across diverse experimental systems. For each perturbation in a Perturb-seq experiment, complementary representation-learning approaches capture distinct aspects of the perturbation response, including differential methods (DE/DGCA), canonical dimensionality-reduction methods (cNMF/PCA), contrastive methods (cPCA, ContraPC, kernel cPCA, kernel ContraPC and ContrastiveVI), and graph-based methods (Hotspot); these representations are pooled into a unified *gene program library*. (c) Gene-level disease-association statistics from MAGMA are modeled using perturbation-derived gene programs as features. Programs carrying disease-relevant information are retained and combined to estimate an optimally weighted *disease meta-program*, whereas non-informative programs receive little or no contribution. This procedure generates MetaProg^(*p*,*d*)^ gene program representations for each perturbation–disease pair, as well as a pooled MetaProg^(ALL,*d*)^ representation integrating programs across all perturbations. (d) Perturbscape prioritizes perturbations for each disease using a trait relevance score (TRS) quantifying the disease heritability informativeness of the corresponding MetaProg^(*p*,*d*)^ meta-programs; MetaProg^(ALL,*d*)^ summarizes disease-relevant signal across perturbations. (e) Enriched pathways within disease-associated meta-programs connect perturbations to downstream disease-relevant biology, enabling construction of perturbation–pathway–disease relationships. (f) Visualization of the perturbation-specific disease meta-program embeddings colored by the TRS highlights perturbation niches in which disease heritability signal is concentrated.

We consider two broad formulations of *Ψ^α^*; a difference-based objective 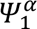 recovers standard cPCA^23^; we term a ratio-based objective 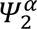 as ContraPC, where *γ* is a small regularization constant (**Methods**). In both formulations, *α* controls the strength of the background contrast, and we evaluate multiple values of *α* to capture different levels of contrastiveness (**Methods**). For the linear formulations, *ϕ*(*x*) = x, whereas for the kernel formulations, *ϕ*(*_x_*) is a nonlinear feature map approximated using scalable random features (**Methods**; **Supplementary Note**). Besides PCA-based approaches, ContrastiveVI provides a complementary nonlinear latent-variable representation.

Together, these approaches capture complementary scales of perturbation response: average transcriptional shifts, dominant variation across cells, variation selectively enriched by perturbation, and locally co-modulated gene modules within cellular state space. Formally, for a perturbation *p*, we obtain a program library 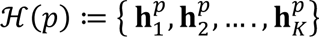, where each 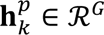 is a continuous or binary gene-weight vector over *G* genes as identified using our suite of methods. The complete program space is ℋ_atlas_ ≔ (∪*_p_*_∈P_ ℋ(*p*)) ∪ ℋ(pooled) where *P* denotes the set of all perturbations and ℋ(pooled) represents programs obtained by jointly modeling all perturbed cells as a single unit against control cells. After removing highly correlated programs, the mean of the screen-level average numbers of retained programs was 113.7 programs per perturbation across 16 analyzed screen–context combinations from 7 Perturb-seq systems (**Methods**; **Table 1**).

**Table 1:** Perturb-seq datasets analyzed with Perturbscape. Summary of the experimental systems, perturbation modalities, cell classes, numbers of expressed genes, cells and perturbations analyzed, and the number of relatively independent gene programs learned per perturbation across the Perturbscape compendium.

| Dataset (technology) | Reference | Cell class | Num. of expressed genes | Number of cells analyzed | Number of perturbations analyzed | Median no. of cells per perturbation | Mean (SD) no. of programs per perturbation |
|---|---|---|---|---|---|---|---|
| K562-GenomeWide (CRISPRi) | Replogle et al. | K562 | 8,175 | 1,801,540 | 7,991 | 202 | 61.1 (9.9) |
| K562-Essential (CRISPRi) | Replogle et al. | K562 | 8,563 | 137,498 | 414 | 268 | 121.9 (33.2) |
| Jurkat-Essential (CRISPRi) | Nadig et al. | Jurkat | 8,882 | 80,082 | 184 | 271 | 132.2 (23.4) |
| HepG2-Essential (CRISPRi) | Nadig et al. | HepG2 | 9,624 | 49,394 | 255 | 136 | 118.4 (47.3) |
| Monocytes KD (CRISPRi) | Yao et al. | THP-1 | 10,000 | 27,601 | 193 | 112 | 109.9 (13.0) |
| Monocytes KO | Yao et al. | THP-1 | 10,000 | 35,090 | 193 | 135 | 115.3 (11.0) |
| T2D CRISPR KO - Day 3 | Liu et al. | hPSC differentiation (DE) | 10,000 | 17,901 | 32 | 450 | 88.3 (21.2) |
| T2D CRISPR KO - Day 7 | Liu et al. | hPSC differentiation (PFG) | 10,000 | 17,544 | 31 | 291 | 80.4 (20.8) |
| T2D CRISPR KO - Day 11 | Liu et al. | hPSC differentiation (PP) | 10,000 | 10,276 | 25 | 182 | 93.2 (30.9) |
| T2D CRISPR KO - Day 18 | Liu et al. | hPSC differentiation (SC-Islet) | 10,000 | 24,843 | 30 | 338 | 174.8 (38.4) |
| In vivo Cortex (CRISPR KO) | Zheng et al. | Developing mouse cortex | 10,000 | 11,688 | 4 | 2,118 | 184.5 (13.8) |
| TeloHAEC (CRISPRi) | Schnitzler et al. | TeloHAEC (human aortic endothelial cells) | 10,000 | 80,329 | 577 | 114 | 64.6 (5.0) |
| Whole Brain in vivo (CRISPR KO) - L4/5 IT CTX Glut | PerturbAI - Shi et al. | Mouse brain (L4/5 IT cortical glutamatergic) | 10,000 | 181,958 | 1,277 | 129 | 130.1 (14.3) |
| Whole Brain in vivo (CRISPR KO) - Pvalb GABA | PerturbAI - Shi et al. | Mouse brain (Pvalb GABAergic) | 10,000 | 158,623 | 1,126 | 126 | 99.0 (9.1) |
| Whole Brain in vivo (CRISPR KO) - TH Prkcd Grin2c Glut | PerturbAI - Shi et al. | Mouse brain (thalamic glutamatergic) | 10,000 | 369,194 | 1,852 | 189 | 121.2 (17.4) |
| Whole Brain in vivo (CRISPR KO) - MB Glut | PerturbAI - Shi et al. | Mouse brain (midbrain glutamatergic) | 10,000 | 299,963 | 1,675 | 166 | 124.5 (16.7) |

For each perturbation *p* and disease or trait *d*, we regress MAGMA gene-level association *z*-scores^26^ on its program library ℋ(*p*) using generalized least squares (GLS) to account for LD-induced correlation (**Figure 1c**; **Methods**). We use leave-one-chromosome-out (LOCO) cross-fitting: for each held-out chromosome, program coefficients are estimated from genes on the remaining chromosomes and then the fitted model is applied to the genes on the held-out chromosome. Concatenating these predictions across chromosomes yields an optimally weighted trait-conditioned combination which we term a *meta-program*. Formally, for gene *g*, the meta-program score is given by MetaProg^(*p*,*d*)^ 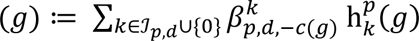 where *c*(*g*) denotes the chromosome containing gene *g*, 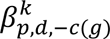 is the GLS coefficient for program *_k_* estimated while excluding genes on chromosome *c*(*g*), 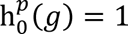, and ℐ*_p_*_,*d*_ ⊆ {1, …, *K*} denotes the indices of informative features selected a priori using a marginal model (**Methods**). This strategy follows the general LOCO strategy used by PoPS and related integrative approaches^19,27,28^ and limits the use of local GWAS signal during weight estimation (see **Supplementary Note** for more details). We further validated this cross-fitting strategy using a split-sample GWAS analysis (**Methods**). We analogously construct an atlas-wide meta-program, MetaProg^(ALL,^*^d^*^)^, by applying the same modeling procedure to the complete perturbation-program library ℋ*_atlas_*. For notational simplicity, we suppress the disease or trait index hereafter and refer to these as MetaProg^(^*^p^*^)^ and MetaProg^(ALL)^.

Perturbscape prioritizes a perturbation based on the heritability informativeness of its meta-program. For each perturbation, we link the top 500 MetaProg^(^*^p^*^)^ genes to variants using tissue-matched ENCODE-rE2G enhancer–gene predictions^w^, and quantify the heritability contribution of the resulting annotation using S-LDSC conditional on 53 baseline epigenomic and coding annotations^17,30^ (**Methods**). We define the standardized S-LDSC effect size, *τ*^∗^, of the resulting annotation as the Trait

Relevance Score, TRS^(^*^p^*^)^, with significance determined by a block jackknife approach (**Methods**, **Figure 1d**). As a secondary metric, we report the heritability enrichment (ENR) and its associated P value. We use *τ*^∗^ as the primary metric because (i) unlike enrichment, it is less sensitive to differences in annotation size, (ii) it quantifies the unique heritability informativeness conditional on the baseline model, and (iii) enables joint heritability analysis of multiple meta-programs. MetaProg^(ALL)^ is the gene-level disease prioritization learned from the complete perturbation atlas and is analogous to PoPS^19^, but derives its features from experimentally induced perturbation programs within an assayed system rather than from precompiled multi-tissue and functional annotations. Evaluating both MetaProg^(ALL)^ and PoPS top N genes using consistently constructed variant annotations and the same conditional S-LDSC framework enables a controlled comparison of the heritability informativeness captured by their respective gene-prioritization strategies. The top 500 per-perturbation MetaProg^(^*^p^*^)^ genes are used for pathway enrichment analysis and constructing perturbation-pathway-disease triplets (**Figure 1e**). To visualize the disease-relevant perturbation landscape, we embed perturbation-specific and all-perturbation meta-programs in a shared UMAP and color each point by its TRS (**Figure 1f**). Perturbscape is implemented as a workflow (**Figure S1**, **Code Availability**), with all gene programs and meta-programs, and perturbation TRS scores released as a public resource (**Data Availability**).

### Perturbscape resolves gene- and perturbation-level red blood cell trait genetics in genome-wide K562 Perturb-seq

We applied Perturbscape to a genome-wide CRISPRi Perturb-seq screen in K562, an erythroleukemia cell line that captures aspects of hematopoietic progenitor and erythroid biology^9^ (7,991 perturbations with ≥100 cells each, 1.8 million cells). We focused on two well-powered hematologic biomarker GWAS traits whose genetic architectures have previously shown strong relevance to K56215,31: mean corpuscular hemoglobin (MCH) and red blood cell count (RBC count)^32^ (average GWAS N=444,430). **Figure 2a** presents the UMAP embedding of the per-perturbation meta-programs for these two red blood cell traits, colored by the TRS scores. Higher TRS scores localized to discrete regions of the embedding exhibiting specific pathway enrichments among their corresponding meta-programs (**Figure S2a**). Relative to the individual-perturbation TRS scores, the atlas meta-program MetaProg^(ALL)^ showed at least 2.7x and 2.3x higher TRS for MCH and RBC, respectively (**Figure 2b**, **S2b**), and its annotations explained 2.0x and 1.8x greater per-SNP heritability enrichment than the strongest individual perturbations. This atlas-level signal reflected distributed contributions across perturbations rather than dominance by a small number of highly redundant programs: for MCH and RBC respectively, 68% and 77% of perturbations shared fewer than 10% of their top 500 MetaProg^(^*^p^*^)^ genes with the top 500 MetaProg^(ALL)^ genes (**Figure S2c**). For both traits, MetaProg^(ALL)^ was enriched in canonical erythropoietic pathways such as *RUNX1* transcriptional regulation and Kit receptor signaling (**Figure S2d**). Joint heritability analysis of the meta-programs retained MetaProg^(ALL)^ together with 2 nominally significant (P < 0.05) perturbation-specific annotations for each of MCH and RBC: *SIGLEC14* and *UBE2V1* for MCH, and *RCL1* and *MYL6B* for RBC (**Methods**, **Figure 2b**, **S2b**). *SIGLEC14* and *MYL6B* implicated distinct pathways, including CREB1 phosphorylation (padj = 7.7e-02) and Gβγ signaling (padj = 1e-02), respectively (**Table 2**).

**Figure 2:**
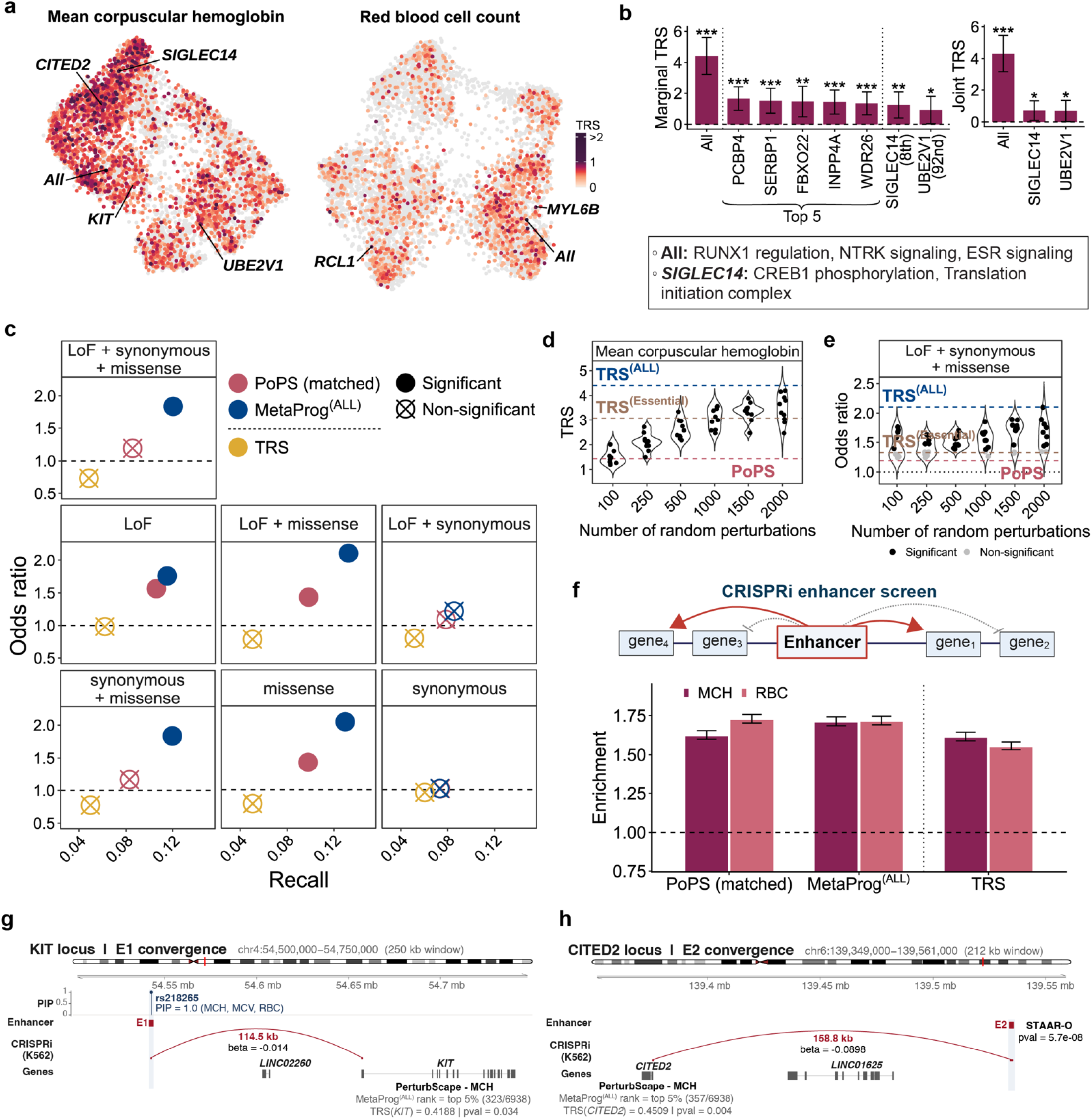
Perturbscape identifies disease-relevant perturbation programs for erythroid traits and converges with orthogonal genetic and functional evidence. (a) UMAP embedding of perturbation-specific meta-programs for 7,991 perturbations from a K562 whole-genome Perturb-seq screen for two red blood cell traits: mean corpuscular hemoglobin (MCH; left) and red blood cell count (RBC; right). Each perturbation is colored by its trait relevance score (TRS), with non-significant perturbations shown in gray. Top enriched pathways associated with the meta-programs of selected perturbations and the integrated MetaProg^(ALL)^ are highlighted. (b) Bar plots showing the marginal TRS of MetaProg^(ALL)^ and 7 individual perturbations with high TRS for MCH (left; the top 5—*PCBP4*, *SERBP1*, *FBXO22*, *INPP4A* and *WDR26*—together with *SIGLEC14* and *UBE2V1*, ranked 8^th^ and 92^nd^). Joint heritability analysis identified 3 independently informative annotations, MetaProg(ALL), MetaProg^(*SIGLEC*14)^ and MetaProg^(^*^UBE2V1^*^)^ (right). All analyses were conditioned on 53 baseline annotations comprising coding, conserved and epigenomic annotations. Error bars represent 95% confidence intervals and asterisks denote statistical significance (***P < 0.001, **P < 0.01, *P < 0.05). (c) Recall and enrichment (odds ratio) of genes implicated by rare-variant association analyses across different annotation masks—loss-of-function (LoF), missense and synonymous variants, individually and in combination—among (i) the top 500 MetaProg^(ALL)^ genes, (ii) the top 500 genes ranked by perturbation-level TRS and (iii) the top 500 PoPS genes. The dashed horizontal line denotes an odds ratio of 1. Solid colored points indicate significant enrichment (one-sided Fisher’s exact test, P < 0.05). (d) Violin plots showing the distribution of TRS estimates for MetaProg^(ALL)^ constructed from random subsets of 100–2,000 perturbations for MCH; for each subset size, 10 random perturbation subsets were sampled. Dashed lines indicate the TRS of MetaProg^(ALL)^ from the full 7,991-perturbation screen (blue), MetaProg^(ALL)^ from an independent screen of 414 essential-gene perturbations (gold), and the TRS corresponding to PoPS gene prioritization (red). (e) Violin plots showing the distribution of rare-variant enrichment (odds ratio, LoF + synonymous + missense mask) among the top MetaProg^(ALL)^ genes as a function of the number of random perturbations (100–2,000) used to construct MetaProg^(ALL)^ for MCH; for each subset size, 10 random perturbation subsets were sampled. Points are colored by nominal significance (one-sided Fisher’s exact test; black, significant; gray, non-significant). Dashed lines indicate the reference odds ratios for MetaProg^(ALL)^ from the full 7,991-perturbation screen (blue), MetaProg^(ALL)^ from the independent 414 essential-gene screen (gold), and PoPS (red); the dotted line denotes an odds ratio of 1. (f) Enrichment of experimentally linked enhancer target genes from three CRISPRi enhancer perturbation screens among the top two genes prioritized within a 1-Mb window of each enhancer using continuous MetaProg(ALL), PoPS and TRS scores, shown for both MCH (dark) and RBC (light). Enrichment is calculated relative to a control in which two genes are randomly selected from the same 1-Mb window. Error bars represent 95% confidence intervals. (g) Convergence of common-variant, enhancer perturbation and Perturbscape evidence at the *KIT* locus. The fine-mapped variant rs218265 (PIP = 0.99 for MCH, mean corpuscular volume (MCV) and RBC) overlaps enhancer E1 (chr4: 54,542,505–54,543,005), which is linked by K562 CRISPRi to a distal gene *KIT*, located 114.5 kb from the enhancer. Perturbscape independently prioritizes *KIT* for MCH, with MetaProg^(ALL)^ ranking *KIT* in the top 5% of genes and *KIT* CRISPRi perturbation showing a significant TRS (TRS^(^*^KIT^*^)^ = 0.42, P = 0.034). (h) Convergence of rare-variant burden, enhancer perturbation and Perturbscape evidence at the *CITED2* locus. Rare-variant burden-level association at enhancer E2 (chr6: 139,534,045–139,535,314) (STAAR-O P = 5.7e-08) coincides with a K562 CRISPRi enhancer–gene link to *CITED2*, located 158.8 kb from the enhancer. Perturbscape independently prioritizes *CITED2* for MCH, with MetaProg^(ALL)^ ranking the gene in the top 5% and *CITED2* CRISPRi perturbation showing a significant TRS (TRS^(^*^CITED2^*^)^ = 0.45, P = 0.004). Source data are provided as a Source Data file.

**Table 2:** Examples of perturbation–trait–pathway triplets. Selected perturbations whose meta programs are significantly associated with a complex trait or disease, each paired with a biologically relevant pathway enriched in the program. For each triplet, the table reports the dataset and technology, cell class, trait, perturbed gene, trait relevance score (TRS) and its p-value, and the pathway with its enrichment p-value and q-value.

| Dataset (technology) | cell class | trait | perturbation | TRS | TRS p-value | pathway | pathway p-value | pathway q-value |
|---|---|---|---|---|---|---|---|---|
| K562-GenomeWide (CRISPRi) | K562 | MCH | SIGLEC14 | 1.2489 | 1.7E-03 | CREB1 phosphorylation through the activation of Adenylate Cyclase | 2.8E-04 | 7.7E-02 |
| K562-GenomeWide (CRISPRi) | K562 | RBC | MYL6B | 0.9248 | 2.2E-03 | G beta:gamma signalling through PLC beta | 4.5E-05 | 1.2E-02 |
| HepG2-Essential (CRISPRi) | HepG2 | Cholesterol | ATP6AP2 | 1.7750 | 3.0E-02 | Cholesterol biosynthesis | 1.7E-06 | 6.0E-04 |
| HepG2-Essential (CRISPRi) | HepG2 | Alanine aminotransferase | HSPA5 | 1.0551 | 1.9E-05 | Plasma lipoprotein remodeling | 1.3E-04 | 2.8E-02 |
| HepG2-Essential (CRISPRi) | HepG2 | LDL-C | RABGGTA | 1.4510 | 4.6E-02 | Simvastatin Action Pathway (SMPDB) | 2.8E-11 | 1.6E-09 |
| Monocytes KD (CRISPRi) | THP-1 | Primary biliary cirrhosis | CCR6 | 1.0161 | 3.4E-03 | Interferon Signaling | 8.3E-10 | 1.1E-06 |
| Monocytes KO | THP-1 | Multiple sclerosis | STK4 | 1.4180 | 1.2E-03 | Cytokine Signaling in Immune system | 1.7E-04 | 8.7E-02 |
| T2D CRISPR KO - Day 18 | hPSC differentiation (SC-Islet - all cells) | T2D | TLE3 | 0.2953 | 2.5E-03 | Maturity onset diabetes of the young (KEGG) | 3.4E-04 | 1.3E-02 |
| T2D CRISPR KO - Day 18 | hPSC differentiation (SC-Islet - endocrine cells) | T2D | FOXA1 | 0.4612 | 8.7E-03 | Insulin secretion (KEGG) | 1.6E-05 | 1.9E-02 |
| T2D CRISPR KO - Day 18 | hPSC differentiation (SC-Islet - non-endocrine cells) | T2D | GATA6 HET | 0.3735 | 4.8E-02 | FOXA2 pathway | 3.4E-03 | 8.0E-02 |
| Whole Brain in vivo (CRISPR KO) - L4/5 IT CTX Glut | Mouse brain (L4/5 IT cortical glutamatergic) | Autism | Bin1 | 0.5145 | 3.2E-03 | Glutamate binding, activation of AMPA receptors and synaptic plasticity | 3.4E-04 | 7.9E-02 |
| Whole Brain in vivo (CRISPR KO) - Pvalb GABA | Mouse brain (Pvalb GABAergic) | Autism | Gad1 | 0.5031 | 4.5E-03 | Voltage gated Potassium channels | 1.6E-09 | 1.8E-06 |
| Whole Brain in vivo (CRISPR KO) - TH Prkcd Grin2c Glut | Mouse brain (thalamic glutamatergic) | Autism | Grin3a | 0.5256 | 5.6E-03 | Negative regulation of NMDA receptor-mediated neuronal transmission | 5.9E-04 | 4.8E-02 |
| Whole Brain in vivo (CRISPR KO) - MB Glut | Mouse brain (midbrain glutamatergic) | Autism | Grin2d | 0.9203 | 3.1E-05 | Unblocking of NMDA receptors, glutamate binding and activation | 3.4E-03 | 2.7E-02 |

We next benchmarked the disease-gene prioritization performance of Perturbscape against PoPS using rare-variant gene burden associations^33,34^ across 7 functional variant masks with alternate allele frequency (AAF) < 0.001 (**Methods**). For matched comparison with Perturbscape, we re-fit PoPS using its original feature set but restricted to the universe of expressed genes in the perturbation data. For MCH, the top 500 MetaProg^(ALL)^ genes exhibited 1.8-fold enrichment (P=1.2e-04) among burden genes implicated by the aggregate synonymous + missense + loss-of-function (LoF) mask, whereas top 500 PoPS genes showed no significant enrichment (1.2x enrichment, P=0.18) (**Figure 2c**). Alignment was strongest for missense-mask burden and weaker for LoF and synonymous masks. Because the common-variant MAGMA and rare-variant burden gene sets showed minimal direct overlap (Jaccard index < 0.063 across masks) (**Figure S2e**), these results indicate that Perturbscape meta-programs capture pathway-level mechanistic convergence between the common and rare genetic architectures of red blood cell traits. Notably, extending prior observations of common–rare pathway convergence^35–37^, perturbation-derived gene programs captured this convergence more strongly than the steady-state features in PoPS, demonstrating that, in this setting, experimentally induced gene programs can capture common–rare genetic convergence not recovered as strongly by steady-state PoPS features. By contrast, genes ranked directly by perturbation TRS were not enriched for burden genes (P = 0.56– 0.94 across masks), indicating that an upstream perturbed gene need not show direct genetic association for its downstream program to be disease relevant, as also noted in refs^12–14^. Perturbation TRS was not correlated with the MAGMA z-scores of the perturbed genes (**Figure S2f**), further suggesting that upstream perturbed genes driving disease-relevant programs need not themselves be GWAS-implicated genes.

To test the robustness of Perturbscape to perturbation coverage and resolution, we reconstructed MetaProg^(ALL)^ from 10 random subsets each of 100, 250, 500, 1,000, 1,500 and 2,000 perturbations. Both TRS^(ALL)^ and the corresponding heritability enrichment increased markedly with the number of perturbations for MCH and RBC count (**Figure 2d**, **S2g-h**). Statistical significance also generally increased with the number of perturbations, but with a weaker trend (**Figure S2h**). Importantly, this increase was not restricted to the common-variant signal used to construct disease-informed meta-programs: enrichment among independently identified WES rare-variant burden genes increased in parallel with perturbation coverage (**Figure 2e**, r=0.42, p=9e-04; **Figure S2i**). This orthogonal genetic validation supports the interpretation that broader perturbation coverage progressively captures additional disease-relevant biological programs rather than simply improving fit to the common-variant prediction task. An independent K562 essential-gene CRISPRi screen comprising 414 perturbations produced similar behavior to comparably sized subsets of the genome-wide screen. We next assessed robustness to the resolution at which perturbations were modeled. For 25 high-TRS MCH perturbations with two guides represented by ≥100 cells each, we repeated program learning and disease-conditioned meta-program analysis separately for each guide rather than after gene-level aggregation. 76% of perturbations had at least one guide with nominally significant positive TRS, and 56% had both guides with TRS > 0.2 (**Figure S2j**). These results indicate broad concordance between gene- and guide-level disease relevance while also revealing guide-to-guide heterogeneity.

We next tested convergence of Perturbscape results with orthogonal cis-regulatory evidence from multiple independent K562 CRISPRi enhancer screens^38–40^ harmonized in ref^29^. For each CRISPRi-significant enhancer, we asked whether its experimentally regulated target ranked among the top two MetaProg^(ALL)^ genes within 1 Mb. MetaProg^(ALL)^ achieved 1.71- and 1.72-fold enrichment over random selection for MCH and RBC count respectively, exceeding PoPS for MCH and matching it for RBC count (1.63- and 1.73-fold; one-sided Z-test P=9.1e-06 and 7.1e-01; **Figure 2f**). This is notable given that MetaProg^(ALL)^ is learned from perturbation responses within a single cellular system, whereas PoPS integrates a broad multi-tissue feature set. Ranking perturbed genes by TRS also recovered CRISPRi-linked targets with substantial enrichment (1.62-fold and 1.56-fold; P < 2.2e-16 for MCH and RBC count, respectively), although somewhat less strongly than MetaProg^(ALL)^ and PoPS. This demonstrates the convergence between cis-regulatory evidence from combining CRISPR and GWAS with the trans pathway level convergence from Perturbscape.

We highlight two loci supported by complementary genetic and perturbational evidence. First, an enhancer at chr4:54,542,505-54,543,005 (GRCh38), which contains a fine-mapped MCH GWAS variant (rs218265; PIP = 0.99), is linked to *KIT* in K562 by enhancer CRISPRi (β=-0.014, adjusted p=1.4e-02) (**Figure 2g**). *KIT* was the top-ranked gene by MetaProg^(ALL)^ (overall rank 323) within the 1Mb window around the enhancer and showed significant TRS for MCH (TRS=0.42, p=3.4e-02), thereby demonstrating convergence of fine-mapped GWAS, enhancer-perturbation and trans-perturbation evidence. *KIT* encodes the receptor tyrosine kinase for stem cell factor required for erythroid progenitor proliferation^41^ and severe *Kit* deficiency produces macrocytic anemia in mouse models, consistent with an MCH-relevant erythroid role^42^. MetaProg^(^*^KIT^*^)^ prioritized heme-biosynthesis genes like *FECH* (rank 177) and *PPOX* (rank 381)^43,44^. Next, we performed enhancer-level STAAR-O burden testing^45,46^ for MCH across all 3,670 CRISPRi-tested enhancers using UK Biobank whole-genome sequencing data, treating all variants within each enhancer as a single variant mask. An enhancer at chr6:139,534,045-139,535,314 linked by CRISPRi to *CITED2* (β=-0.09, p=2.4e-03) carried significant rare-variant burden for MCH (STAAR-O p = 5.7e-08); *CITED2* was again the top-ranked gene in the 1 Mb window by MetaProg^(ALL)^ (overall rank 357) and showed significant TRS for MCH (TRS=0.45, p = 4.4e-03) (**Figure 2h**). Consistent with this evidence, *CITED2* depletion in K562 reduces *GATA2* expression, while hematopoietic-specific *Cited2* loss in mice reduces erythroid progenitors and produces mild anemia^47–49^. An additional *EPB41*-linked enhancer example is provided in **Figure S2k**.

Collectively, in K562, one of the most deeply CRISPR-characterized cell lines, Perturbscape prioritizes RBC-trait genes and perturbations that converge across common-variant GWAS, rare-variant genetic associations and independent enhancer CRISPRi screens, while resolving the trans-regulatory programs through which these signals connect to erythroid biology.

### Perturbscape resolves lipid-relevant perturbation programs and therapeutic pathways in HepG2

We extended Perturbscape to a CRISPRi Perturb-seq screen targeting 255 essential genes in HepG2 hepatocellular carcinoma cells and evaluated perturbation programs across 10 lipid and hepatic traits (**Table 1, Table S2**). We focused on perturbations represented by ≥100 assigned cells to ensure sufficient cellular coverage for program learning. **Figure 3a** and **Figure S3a** present the UMAP embeddings of the per-perturbation meta-programs, colored by TRS. Among the traits, lipoprotein(a) exhibited both the highest average TRS^(*p*)^ across perturbations and the highest TRS^(ALL)^, followed by triglycerides, cholesterol and LDL cholesterol (**Figure 3b**, **S3b**). Unlike K562, the HepG2 meta-program landscape did not separate into prominent global clusters, reflective of the restricted essential-gene perturbation set. Nevertheless, for individual traits, high-TRS perturbations localized to discrete regions of the embedding. Across perturbations, the meta-analyzed TRS for lipid traits was on average 9.5-fold higher than that for hepatic traits (paired t-test p = 4.9e-26; **Figure S3c**). Perturbations showing the strongest relative lipid-trait relevance over hepatic traits included *EMC3* (rank 12) and *RAB4B* (rank 14), both with known lipid-related biological significance^50,51^ (**Figure S3d**). Several perturbation-pathway-trait triplets recovered established hepatic biology; notable examples include *ATP6AP2* → Cholesterol biosynthesis → Cholesterol^52^ and *HSPA5* → Plasma lipoprotein remodeling → Alanine aminotransferase^53^ (**Table 2**). Across both lipid and hepatic traits, MetaProg^(ALL)^ showed at least 2.2x and 2.0x higher meta-analyzed TRS than the strongest individual perturbation-specific programs, respectively (**Figure S3e**). Nevertheless, 7 perturbation-specific meta-programs remained nominally significant (P < 0.05) in a joint model with MetaProg^(ALL)^ for at least one trait (**Figure S3e**).

**Figure 3:**
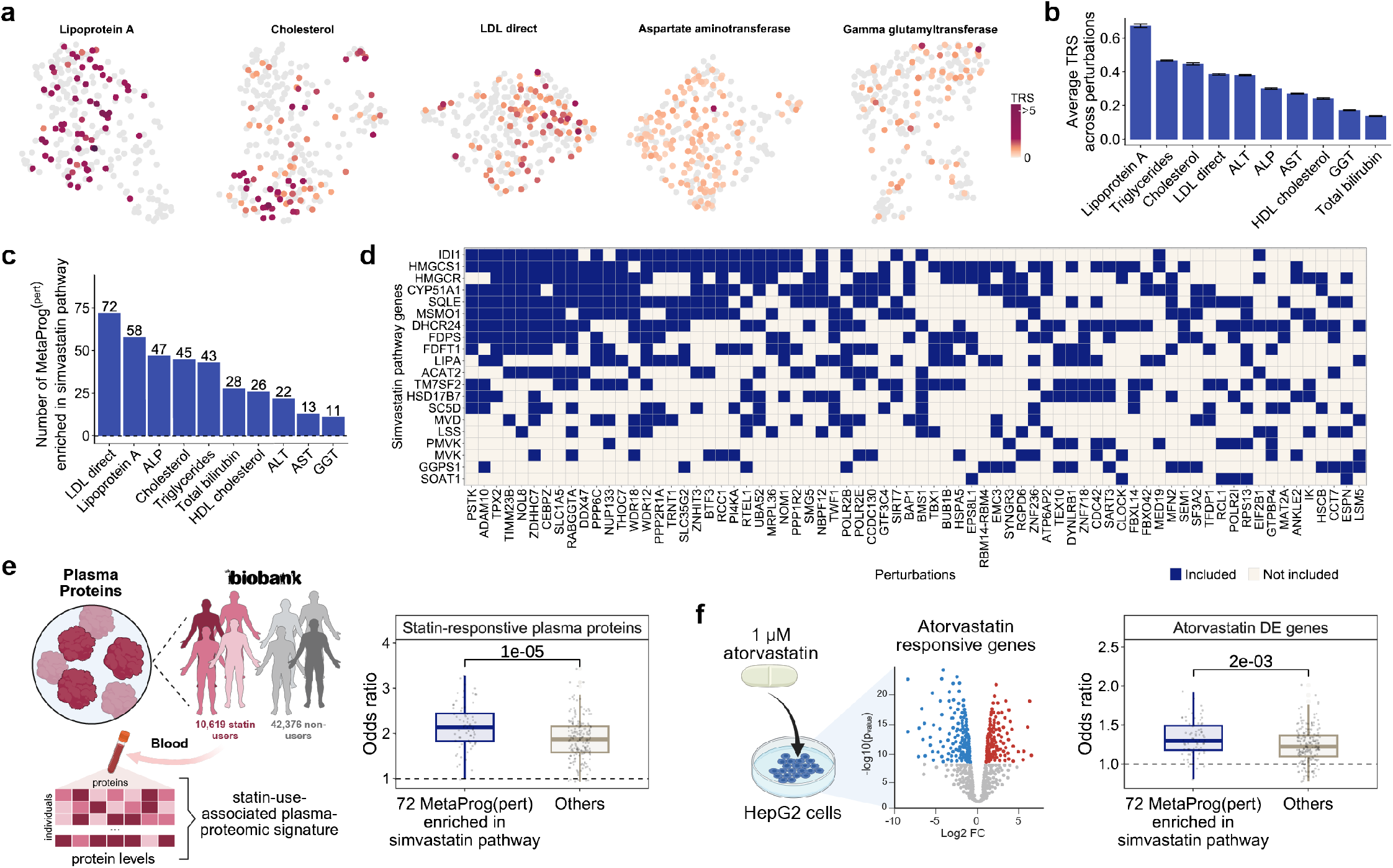
Perturbscape identifies trait-relevant perturbation programs in HepG2 and converges on statin-responsive biology. (a) UMAP visualization of perturbation-specific meta-programs from a HepG2 essential-gene Perturb-seq screen (n = 255) for representative lipid and hepatic traits, including Lipoprotein(a), Cholesterol, Direct LDL cholesterol, Aspartate Aminotransferase and gamma-glutamyltransferase (GGT). Each perturbation is colored by its trait relevance score (TRS), with non-significant perturbations shown in gray, highlighting distinct regions of the perturbation-response landscape associated with individual traits. (b) Bar plot showing the average TRS across perturbations for each of 10 lipid and hepatic quantitative traits (**Table S2**), ordered by descending average TRS. Error bars represent 95% confidence intervals. (c) Bar plot showing, for each of 10 lipid and hepatic quantitative traits, the number of essential-gene perturbations whose top 500 meta-program genes are significantly enriched for the simvastatin-response pathway (SMPDB). Traits are ordered by the number of perturbation-specific meta-programs showing significant pathway enrichment. (d) Binary heatmap showing the presence of simvastatin-response pathway genes among the top 500 genes of perturbation-specific direct LDL cholesterol meta-programs for 72 essential-gene perturbations significantly enriched for the simvastatin-response pathway. Columns represent perturbations and rows represent genes in the simvastatin-response pathway; values indicate presence (included) or absence (not included) in the top 500 meta-program genes. (e) Orthogonal validation of perturbations whose meta-programs are enriched for the simvastatin-response pathway, using plasma proteomics. Box plots show the enrichment odds ratio for overlap between the top 500 genes of the direct LDL cholesterol meta-programs of the 72 simvastatin-response-associated perturbations and 1,525 differentially abundant plasma proteins between UK Biobank participants reporting statin use (N = 10,619) and non-users (N = 42,376) in the White British UKB-PPP cohort, compared with all other perturbations (“Others”). Significance (P=1e-05) was assessed using a one-sided two-sample t-test. The dashed line denotes an odds ratio of 1. (f) Orthogonal validation using an *in vitro* atorvastatin perturbation. Box plots show the enrichment odds ratio for overlap between the top 500 genes of the direct LDL cholesterol meta-programs of the 72 simvastatin-response-associated perturbations and 2,166 genes differentially expressed in HepG2 cells following treatment with 1 µM atorvastatin, compared with all other perturbations (“Others”). Significance (P=2e-03) was assessed using a one-sided two-sample t-test. The dashed line denotes an odds ratio of 1. Source data are provided as a Source Data file.

We next asked whether some of the heritability-enriched perturbation pathways are informative for pharmacologic signatures. The simvastatin pathway (SMPDB) was enriched in LDL cholesterol (LDL-C) MetaProg^(^*^p^*^)^ meta-programs (top 500 genes) of 72 perturbations (padj < 0.1), representing a 1.2-to 6.5-fold greater number compared to other lipid and hepatic traits (**Figure 3c**). **Figure 3d** shows the representation of the statin pathway genes among the top 500 LDL-C meta-program genes for these 72 perturbations. Among the perturbations with the strongest enrichment was *RABGGTA* (OR=29.3, P=7.1e-12), consistent with its previously reported associations with statin pharmacology^54,55^. By contrast, the simvastatin pathway was not significantly enriched among the top 500 PoPS genes for LDL-C (padj = 0.12; PoPS re-fit to genes expressed in the HepG2 perturbation data). Because liver secretions are a major constituent of the plasma proteome, we next assessed whether these 72 perturbation meta-programs also aligned with statin-use-associated plasma-proteomic signatures in a population cohort. In the UKB-PPP cohort of White British individuals, we compared normalized plasma protein levels between 10,619 self-reported statin users and 42,376 non-users, adjusting for age, sex, 20 ancestry principal components, BMI, and standing height (**Methods**, **Figure 3e**). The 1,525 statin-use-associated proteins recovered were more strongly enriched among the top 500 LDL-C MetaProg^(^*^p^*^)^ genes of the 72 perturbations compared to the other perturbations (mean odds ratio=2.17 versus 1.87; 1-sided t-test P=1.5e-05) (**Figure 3e**). This difference was retained when restricting the analysis to the 500 proteins showing the strongest statin-use associations (**Figure S3f**).

Next, we tested whether these perturbation meta-programs align with a directly induced statin response *in vitro*. We derived an atorvastatin-response signature from RNA-seq of HepG2 cells treated with 1 μM atorvastatin versus controls using differential-expression analysis^56^. Atorvastatin-responsive genes were more strongly enriched among the top 500 LDL-C MetaProg^(^*^p^*^)^ genes of the 72 statin-pathway-associated perturbations than among those of the remaining perturbations (mean odds ratio = 1.33 versus 1.24, one-sided t-test P=2.0e-03; **Figure 3f**). Compared to other trait-conditioned meta-programs showing simvastatin-pathway enrichment, LDL-C meta-programs showed the strongest concordance with both atorvastatin-induced transcriptional responses and statin-use-associated plasma proteomic signatures (**Figure S3g**). These differences remained significant after removing simvastatin-pathway genes (**Figure S3h**). Together, these results connect perturbation-derived LDL-C programs to canonical statin biology, a directly induced atorvastatin response in HepG2, and population-scale plasma-proteomic signatures associated with statin use.

### Systematic benchmarking of Perturbscape establishes robustness and identifies key design features

Besides K562 and HepG2, we applied Perturbscape to perturbation screens in 2 other cell lines: (i) CRISPRi targeting 184 essential genes in Jurkat T-cell leukemia cells, analyzed for 19 blood-mediated traits; and (ii) CRISPRi targeting 577 expressed genes near CAD GWAS loci in TeloHAEC endothelial cells, analyzed for 3 cardiovascular traits (**Figure S4**, **S5**). Across the four cell systems, atlas-wide meta-programs showed preferential heritability relevance for matched traits (**Figure S6**). We therefore benchmarked Perturbscape against PoPS across these matched cell system–trait pairs, as both frameworks use a LOCO strategy to learn disease-specific feature weights and generate chromosome-held-out gene scores, while differing in the underlying features: perturbation-derived gene programs for Perturbscape versus steady-state multi-tissue features for PoPS. To account for differences in gene coverage, we compared the top 500 genes from Perturbscape MetaProg^(ALL)^ with top 500 genes from (i) the published PoPS scores and (ii) a refit PoPS model restricted to the genes expressed in the Perturb-seq screen (PoPS-matched).

Across the 4 cell systems and their matched traits, Perturbscape showed significant TRS (FDR-adjusted p < 0.05) for all 34 traits (**Figure 4a**); Perturbscape MetaProg^(ALL)^ achieved up to 4.3-fold and 3.4-fold higher TRS than PoPS-published and PoPS-matched, respectively, with higher average TRS across traits overall (**Figure S7a-b**). In joint S-LDSC models containing all three annotations (MetaProg^(ALL)^, PoPS-matched, PoPS-published), MetaProg^(ALL)^ exhibited significant association (FDR-adjusted p < 0.05) for all 34 traits, compared with 24 and 22 for PoPS-published and PoPS-matched, respectively (**Figure 4a**). Beyond the heritability evaluation, we benchmarked Perturbscape against PoPS-matched for recovery of 229 silver-standard fine-mapped credible-set–gene links, as defined previously^19,29^, assessing whether the linked gene ranked among the top 1 or 2 genes within a 500-kb window around each credible set (**Methods**). MetaProg^(ALL)^ achieved significantly higher recall than the random gene ranking in the window (one-sided z-test p = 5.8e-06 and 1.0e-08 for top 1 and top 2) and performed comparably to PoPS-matched (**Figure 4b**). These analyses show that perturbation-derived programs from matched cellular systems capture disease-genetic information that is competitive with, and in several benchmarks exceeds, that captured by the broad multi-tissue features incorporated by PoPS, despite using 3.4-fold fewer underlying features on average (**Figure S7c**). Beyond this aggregate performance, Perturbscape also prioritized complementary disease genes: among the top 50 MetaProg^(ALL)^ genes in each system and matched trait, 51–94% were absent from the corresponding PoPS-matched top 500 genes (**Figure S7d**). This Perturbscape-specific set included established trait-relevant genes, such as *NIX*, an essential regulator of mitochondrial clearance during erythroid maturation^57^, for RBC count in K562 (MetaProg^(ALL)^ rank 23; PoPS-matched rank 1,728), and *CD93*, a regulator of angiogenesis and vascular biology^58^, for CAD in TeloHAEC (MetaProg^(ALL)^ rank 37; PoPS-matched rank 2,611). We next tested reproducibility in a split-sample RBC-count GWAS comprising two non-overlapping UK Biobank subsets (N = 215K White British individuals each). MetaProg^(ALL)^ scores learned independently in the two splits were concordant (Spearman *ρ* = 0.55; **Figure S7e**) and cross-split S-LDSC evaluation produced TRS estimates similar to those from the full GWAS for both Perturbscape and PoPS-matched (**Figure S7f**; **Methods**). These results support generalization of disease-conditioned meta-programs to independent population samples.

**Figure 4:**
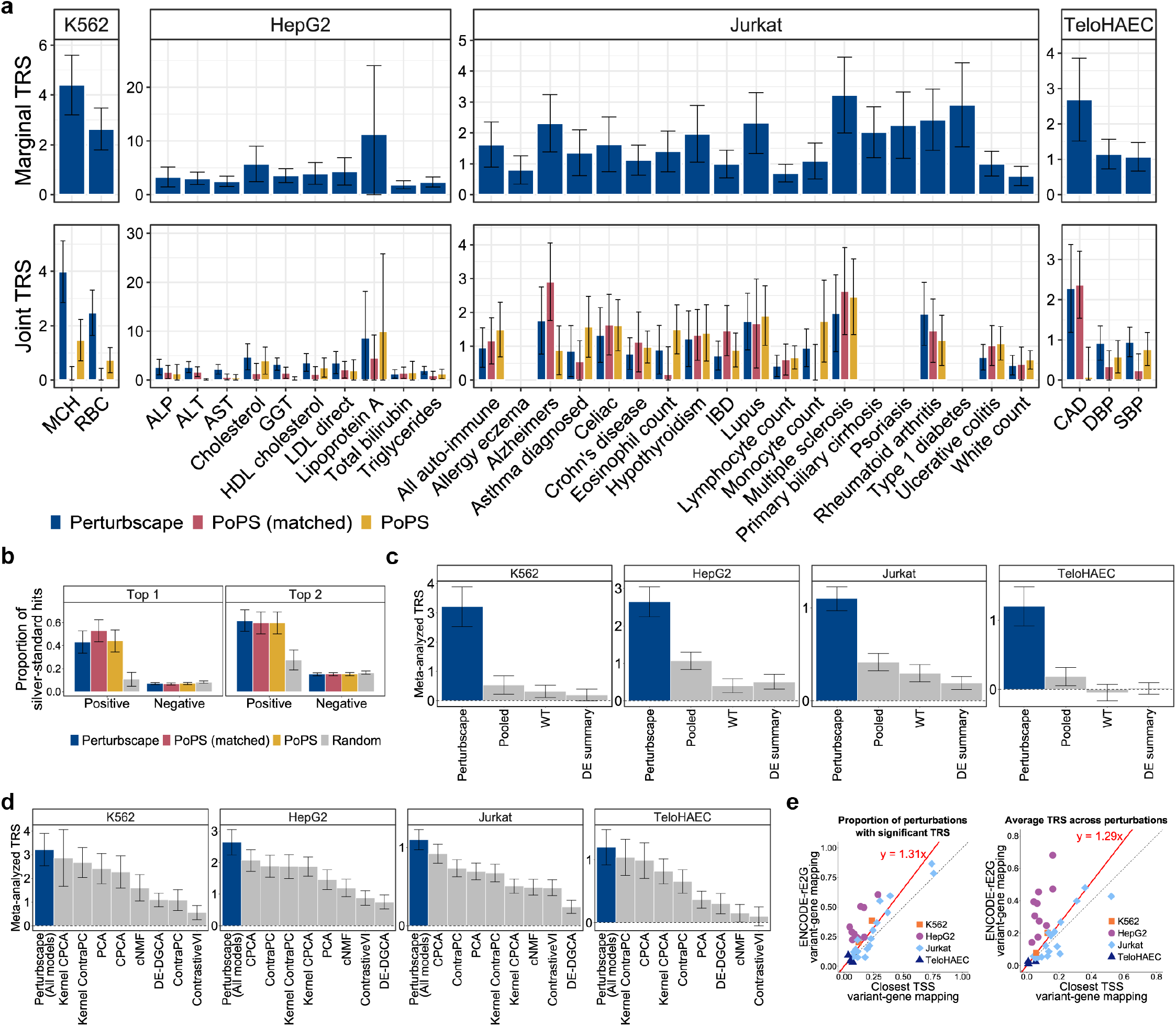
Perturbscape robustly captures disease-relevant perturbation signal across cellular systems and methodological benchmarks. (a) Bar plots showing marginal and joint trait relevance scores (TRS) across four Perturb-seq systems (K562, HepG2, Jurkat and TeloHAEC) and their matched quantitative traits and diseases, spanning erythroid, lipid/hepatic, immune and cardiovascular phenotypes. Top, marginal TRS of Perturbscape MetaProg^(ALL)^ annotations for each matched trait. Bottom, joint TRS comparing Perturbscape MetaProg(ALL), PoPS trained on the same gene universe as Perturbscape, and published PoPS annotations. All heritability analyses are conditioned on 53 baseline annotations comprising coding, conserved and epigenomic annotations, as in Weeks et al.19 Error bars represent 95% confidence intervals. (b) Evaluation against a silver-standard set of 229 fine-mapped credible-set–gene links across 12 UK Biobank traits matched to the cellular systems analyzed here, as previously used for benchmarking in Gschwind et al.29 and Weeks et al.19. Bar plots show the proportion of silver-standard links recovered when the top one (left) or top two (right) genes within a 500-kb window of each credible set were prioritized using Perturbscape MetaProg(ALL), PoPS trained on the corresponding gene universe, published PoPS scores or random gene prioritization. For each trait, Perturbscape predictions were derived exclusively from the biologically matched Perturb-seq system (for example, K562 for MCH and RBC). Error bars represent 95% confidence intervals. (c) Meta-analyzed marginal TRS across tissue-or cell type-matched traits within each of the four cellular systems, comparing Perturbscape MetaProg^(ALL)^ with alternative strategies for constructing meta-programs. Alternative representations were generated from (i) gene programs learned after pooling all perturbations into a single perturbation group (“Pooled”), (ii) gene programs derived from wild-type/control cells comprising non-targeting and safe-targeting controls (“WT”), and (iii) gene programs derived from differential-expression summary statistics for each perturbation relative to control (“DE summary”). See **Methods** for details of alternative gene-program construction. Error bars represent 95% confidence intervals. (d) Meta-analyzed marginal TRS across tissue-or cell type-matched traits within each of the four cellular systems, comparing Perturbscape MetaProg(ALL), which integrates programs across all representation learners, with MetaProg^(ALL)^ constructed separately from each individual representation learner. Individual program learners are ordered by their meta-analyzed TRS. Error bars represent 95% confidence intervals. (e) Comparison of regulatory variant-to-gene mapping using ENCODE-rE2G links versus assignment to the closest transcription start site (TSS). Scatter plots compare the proportion of perturbations with significant TRS (left) and the average TRS across perturbations (right) across traits analyzed in K562, HepG2, Jurkat and TeloHAEC. Each point represents a trait analyzed using both variant-to-gene mapping strategies, with colors indicating the corresponding cellular system. The dotted line represents y = x, and the solid line represents the fitted regression between TRS metrics obtained using ENCODE-rE2G and closest-TSS mapping with zero intercept. Source data are provided as a Source Data file.

We next evaluated key Perturbscape design choices. MetaProg^(ALL)^ achieved at least 2.5-, 3.7- and 5.3-fold higher meta-analyzed TRS than meta-programs derived from (i) pooled perturbed-versus-control cells, (ii) wild-type (WT) or control cells, or (iii) PCA and NMF applied to perturbation-level differential-expression effects, respectively (P < 8.5e-11 in all cases) (**Figure 4c, Methods**); this supports the value of preserving perturbation-level resolution. Among individual program learners, contrastive PC approaches, including cPCA/ContraPC and their kernel extensions, showed the highest average heritability relevance when tested separately (**Figure 4d**). However, removal of any single learner did not significantly reduce overall performance (**Figure S7g**), indicating complementary contributions across representations. Finally, tissue-matched ENCODE-rE2G linking identified 1.3-fold more perturbations with significant TRS than closest-TSS assignment and retained larger positive conditional TRS in 19 of 34 matched trait–system comparisons in joint S-LDSC models (**Figure 4e**, **S7h**). Both strategies produced null-centered TRS after shuffling MAGMA gene scores (**Figure S7i**), arguing against inflation under the null.

Together, these analyses establish the robustness of Perturbscape across cohorts and benchmarks and identify perturbation-resolved program learning, complementary representation learners and tissue-matched regulatory linking as key design features.

### Perturbscape disease signatures are modality-robust and align with state-resolved monocyte trans-eQTLs

We evaluated robustness of Perturbscape disease readouts across perturbation modalities using matched CRISPRi knockdown (KD) and Cas9 knockout (KO) screens targeting the same 598 immune-response genes in LPS-stimulated THP-1 cells (a human monocytic leukemia cell line), with 193 perturbations meeting our ≥100-cell inclusion criterion in both modalities^13^. Across 15 immune-related traits, perturbation-level TRS showed moderate KD-KO concordance (mean Pearson correlation = 0.37), approximately 2.3-to 7.4-fold higher than for blood-cell, brain and lipid/hepatic traits (**Figure 5a**). The two perturbation modalities also identified comparable numbers of disease-relevant perturbations (**Figure S8a**). Across autoimmune diseases and blood-cell traits, an average of 19.1 and 29.4 perturbations, respectively, showed significant overlap between the KD- and KO-derived meta-programs for the same perturbation (Fisher’s exact test, Padj < 0.05), substantially more than observed for brain and lipid/hepatic traits (**Figure S8b**). **Figure 5b** illustrates two autoimmune diseases with reproducible patterns of perturbation-level TRS across the KD and KO screens (results for other immune diseases are reported in **Figure S8c**). Psoriasis showed broadly elevated TRS across perturbations in both modalities, whereas rheumatoid arthritis exhibited a more localized signal concentrated within a shared subset of perturbations (overlap index = 0.51; **Methods**). **Figure 5c** further resolved 33 perturbations with strong cross-modality concordance (r > 0.5) of TRS across 15 immune-related traits. Among the most concordant perturbations were *CCR6*, *STK4*, *TSPAN4* and *IFI30*, genes with established immune-regulatory functions (**Figure S8d-e**).

**Figure 5:**
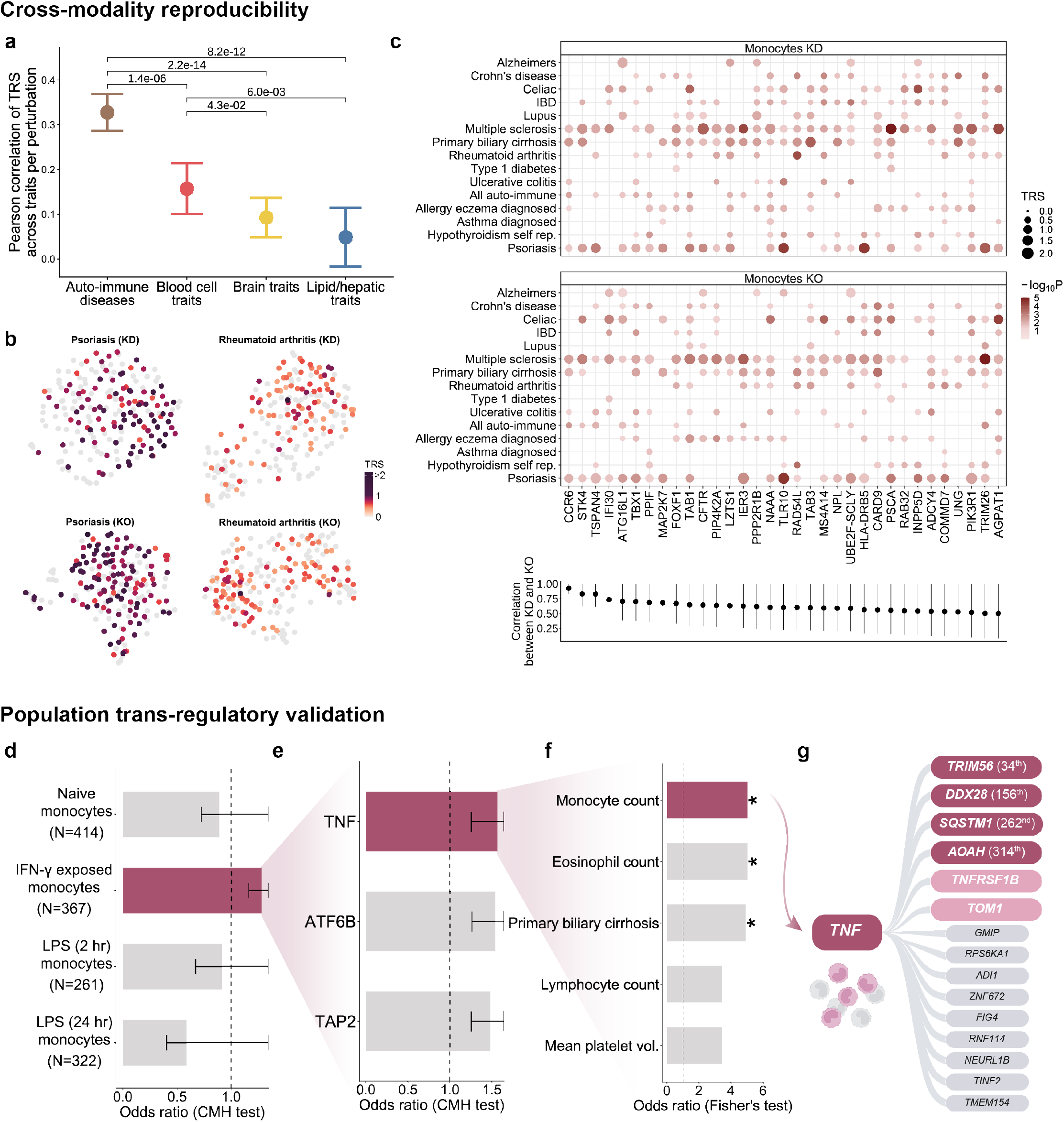
Perturbscape trait-relevance signatures are robust to perturbation modality and align with context-specific trans-regulatory effects in human monocytes. (a) Point estimates (means) with error bars showing the Pearson correlation between perturbation-level TRS profiles obtained from matched CRISPRi knockdown (KD) and Cas9 knockout (KO) perturbations in THP-1 monocytes across four broad trait categories: autoimmune diseases, blood cell traits, brain-related traits and lipid/hepatic traits. For each category, the marker summarizes the per-perturbation Pearson correlation between KD and KO TRS scores across all traits within that category (mean ± 1.96×standard error). Pairwise differences between trait categories were assessed using one-sided paired t-tests. (b) UMAP visualization of perturbation-specific meta-programs from matched KD and KO monocyte Perturb-seq screens for psoriasis (left) and rheumatoid arthritis (right). Perturbations are colored by their trait relevance score (TRS), with non-significant perturbations shown in gray, illustrating concordant disease-relevant structure across perturbation modalities. (c) Concordance of perturbation-specific disease relevance between CRISPRi knockdown (KD) and Cas9 knockout (KO) across 15 immune-related complex diseases in THP-1 monocytes. (Top) Magnitude (TRS, dot size) and significance (−log10P, dot color) of disease relevance for perturbation-specific meta-programs, shown separately for KD and KO. (Bottom) Pearson correlation of TRS scores between KD and KO across the 15 immune-related diseases for 33 perturbations with high correlation (r > 0.5), highlighting perturbations with reproducible disease-relevance signatures across modalities. Error bars represent 95% confidence intervals. (d) Alignment of Perturbscape-derived trans-regulatory links with population trans-eQTLs measured in primary human monocytes across four activation states: naive monocytes, IFN-γ-exposed monocytes, and monocytes profiled 2 h or 24 h following LPS stimulation. For each blood-related trait, perturbation– target links were defined as the top 500 genes from disease-conditioned perturbation-specific meta-programs across 23 blood traits and tested for enrichment among matched (cis-eGene, trans-eGene) pairs corresponding to variants with both cis- and trans-eQTL evidence. Bars show common enrichment odds ratios across perturbations, traits and modalities; the dashed line denotes an odds ratio of 1 and error bars represent 95% confidence intervals. Statistical significance of enrichment was assessed using the Cochran–Mantel–Haenszel (CMH) test. (e) Decomposition of the IFN-γ-exposed monocyte enrichment in (d) by individual cis-eGene perturbation. For each of the three cis-eGenes with both cis- and trans-eQTL evidence in IFN-γ-exposed primary monocytes (*TNF*, *ATF6B* and *TAP2*), the corresponding meta-program was tested for enrichment among its matched trans-eGenes. Bars show common enrichment odds ratios across traits and modalities; the dashed line denotes an odds ratio of 1 and error bars represent 95% confidence intervals. Statistical significance was assessed using the Cochran–Mantel–Haenszel (CMH) test. (f) For each of 5 traits (four blood cell traits—monocyte count, eosinophil count, lymphocyte count and mean platelet volume—and primary biliary cirrhosis), the top 500 genes from the *TNF* meta-program derived from the THP-1 CRISPRi knockdown perturbation screen were tested for enrichment among the trans-eGenes linked to *TNF* as a cis-eGene in IFN-γ-exposed primary monocytes. Bars show the enrichment odds ratios; the dashed line denotes an odds ratio of 1. Statistical significance was assessed using Fisher’s exact test, and asterisks indicate significant enrichment (P < 0.05). (g) Convergence of *TNF*-associated population trans-eQTLs with the *TNF* perturbation meta-program for monocyte count. 15 trans-eGenes linked to *TNF* as a cis-eGene in IFN-γ-exposed primary monocytes are ordered by their meta-program scores for monocyte count. Four trans-eGenes were among the top 500 *TNF* meta-program genes and are highlighted in dark red; two additional genes among the top 1,000 meta-program genes are highlighted in pink. Source data are provided as a Source Data file.

Next, we asked whether the blood-related trans-regulatory programs from Perturbscape correspond to trans-regulatory genetic effects observed in human populations. For each of 23 blood-related traits, we treated the perturbed gene and its top 500 disease-conditioned MetaProg^(p)^ genes as putative regulator-target pairs and tested their enrichment among population-level (cis-eGene, trans-eGene) pairs mapped in primary monocytes derived from 432 individuals across naive, IFN-γ-stimulated and 2-h or 24-h LPS-stimulated states^59^. Enrichment was strongest in IFN-γ-stimulated monocytes (common OR = 1.3; one-sided Cochran-Mantel-Haenszel (CMH) test p=1.6e-05) (**Figure 5d, Methods**). Disease-agnostic regulator–target pairs defined using each perturbation’s top 500 differentially expressed genes showed no significant enrichment in any activation state (p=0.23-0.88) (**Figure S8f**). Thus, disease conditioning enriches for perturbation-derived trans-regulatory relationships also supported by population genetic variation in a context-specific inflammatory state.

In the IFN-γ condition, three perturbed genes (*TNF*, *ATF6B*, *TAP2*) retained significant correspondence with population trans-eQTL relationships after multiple-testing correction (common OR across modalities and traits = 1.48-1.57; FDR-adjusted one-sided CMH test p < 1.8e-02; **Figure 5e**). *TNF* provided an illustrative example: its population trans-eGenes were significantly enriched among the top 500 KD-derived MetaProg^(^*^TNF^*^)^ genes for monocyte count, eosinophil count and primary biliary cirrhosis (OR = 5.03; P = 0.016, 1-sided Fisher’s exact test) (**Figure 5f**). Among the 15 population trans-eGenes associated with *TNF* as a cis-eGene in IFN-γ-stimulated monocytes, 4 genes (*TRIM56*, *DDX28*, *SQSTM1*, *AOAH*) ranked among the top 500 monocyte-count MetaProg^(^*^TNF^*^)^ genes, with *TNFRSF1B* and *TOM1* additionally ranking within the top 1,000 (**Figure 5g**). Several of these genes were also prioritized by MetaProg^(^*^TNF^*^)^ for other enriched diseases and traits (**Figure S8g**). *TRIM56* is a direct amplifier of TNF-induced NF-κB signaling, ubiquitinating *TAK1* (encoded by *MAP3K7*) to potentiate IKK activation^60^, and an interferon effector via the STING pathway^61^. Across traits and modalities, 85 cis-eGene–trans-eGene pairs were jointly supported by Perturbscape and IFN-γ population QTLs. Among KD-supported pairs, cis-regulatory genes were enriched for antigen-processing pathways, whereas their trans targets were enriched for TNF-receptor signaling relative to the corresponding IFN-γ cis- and trans-eGene backgrounds (**Figure S8h**).

Together, these analyses show that Perturbscape disease readouts are reproducible across independent perturbation modalities and recover disease-conditioned trans-regulatory relationships that are supported by context-matched population genetic variation.

### Perturbscape resolves the developmental and cellular emergence of T2D-relevant perturbation programs during β-cell differentiation

We applied Perturbscape to a time-course stem-cell CRISPR-knockout village Perturb-seq dataset comprising 79 barcoded hPSC lines (76 mutant clones and 3 wild-type controls) covering 36 genotypes across 30 key pancreatic developmental regulators during directed differentiation from embryonic stem cell (ESC) to pancreatic β-cell (SC-β)^62^. We analyzed single-cell RNA-seq profiles from four developmental stages: definitive endoderm (DE: day 3), posterior foregut (PFG: day 7), pancreatic progenitor (PP: day 11), and SC-islets (day 18) (**Figure 6a**). We primarily focused our Perturbscape disease-genetic analyses on type 2 diabetes (T2D) and hemoglobin A1c (HbA1c), a quantitative glycemic biomarker relevant to diabetes (average GWAS N = 252,846) (**Table S2**). Because developmental perturbations can alter both cell-intrinsic transcriptional states and lineage composition, we initially analyzed all cells within each stage and subsequently repeated Perturbscape within endocrine and non-endocrine compartments at day 18 (**Figure 6b**, **S9a**-**c**). Further stratification to individual cell types, such as SC-β and SC-*α* cells, was limited by the substantially smaller number of perturbations with sufficient cellular representation for Perturbscape analysis (**Figure S9d**).

**Figure 6:**
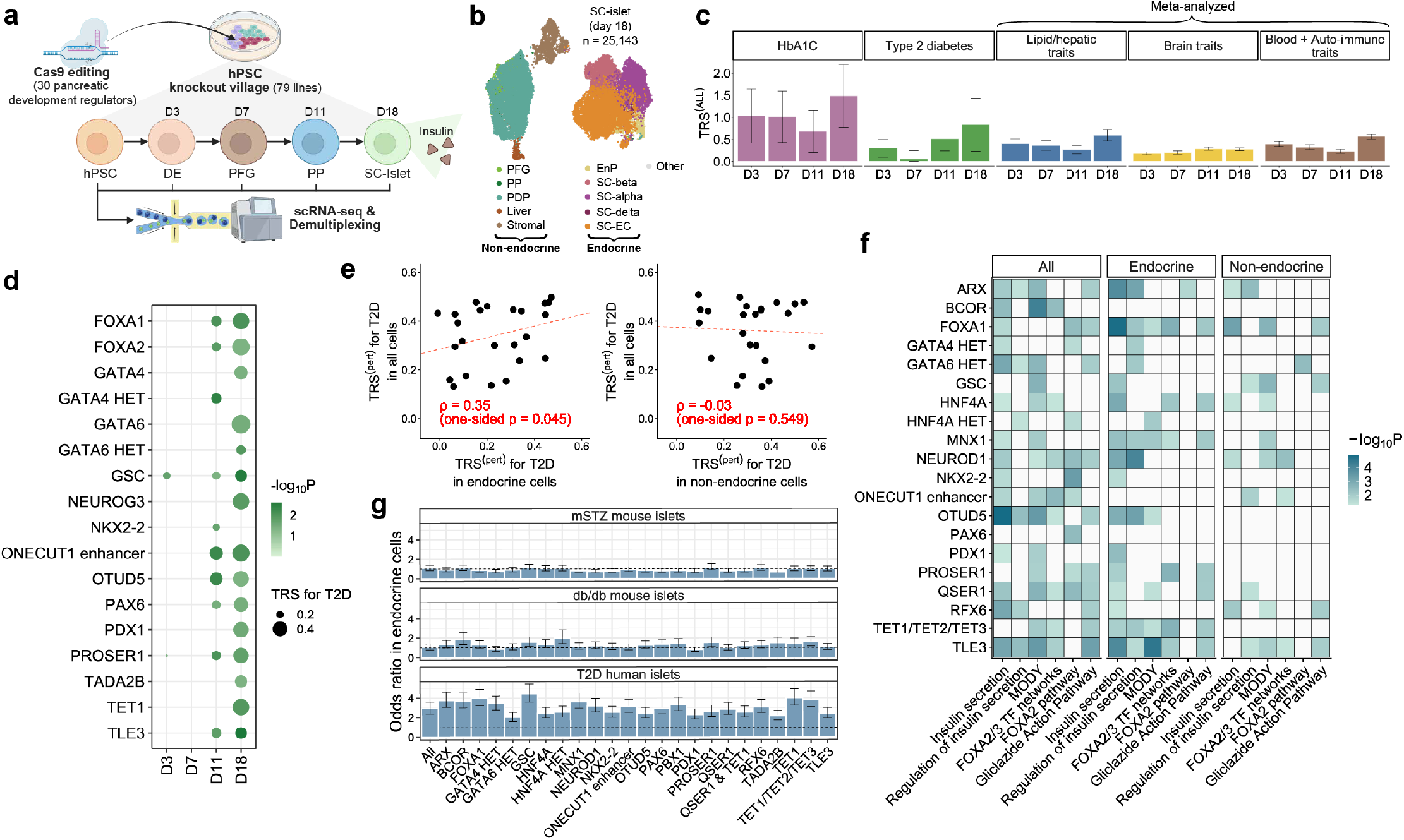
Perturbscape identifies developmental-stage-specific diabetes-relevant perturbation programs during pancreatic differentiation. (a) Schematic of the human pluripotent stem cell (hPSC) pancreatic differentiation CRISPR-KO Perturb-seq screen. Cas9 editing of 30 pancreatic development regulators in hPSCs, assembled in a village culture comprising 79 barcoded lines (76 mutant and 3 WT; 36 genotypes), and differentiated through definitive endoderm (DE; Day 3), posterior foregut (PFG; Day 7), pancreatic progenitor (PP; Day 11) and stem-cell islet (SC-islet; Day 18) stages. Cells were profiled by single-cell RNA sequencing at each developmental stage and demultiplexed across perturbations. (b) UMAP embedding of 24,843 single-cell transcriptomes from the stem-cell islet (SC-islet, Day 18) stage, colored by cell type. Cell types are grouped into non-endocrine populations—posterior foregut (PFG), pancreatic progenitor (PP), pancreatic–duodenal progenitor (PDP), liver and stromal cells—and endocrine populations—endocrine progenitor (EnP), SC-β, SC-α, SC-δ and SC–enterochromaffin (SC-EC) cells. Cell types with fewer than 100 cells at a stage are pooled as “Other”. (c) Bar plots showing MetaProg^(ALL)^ trait relevance scores (TRS) across the four pancreatic differentiation stages. TRS is shown separately for HbA1c and type 2 diabetes (T2D), and meta-analyzed across lipid/hepatic, brain-related, and blood/autoimmune trait categories. Bars correspond to the four developmental stages (Day 3, Day 7, Day 11 and Day 18), illustrating developmental-stage-specific variation in genetic trait relevance. Error bars represent 95% confidence intervals for the individual-trait TRS estimates (HbA1c and T2D) and for the meta-analyzed TRS estimates within each broad trait category. (d) Perturbation-specific TRS for T2D across pancreatic differentiation. Magnitude (TRS, dot size) and significance (−log10P, dot color) of T2D disease-heritability relevance of the meta-programs corresponding to perturbations at each of the four developmental stages (Days 3, 7, 11 and 18). Results are reported for 17 out of 30 perturbations with nominally significant (P < 0.05) TRS at one or more developmental stages. (e) Scatter plots comparing perturbation-level T2D TRS obtained from profiles of all cells (y-axis) with those obtained from endocrine cells (left) and non-endocrine cells (right) at the SC-islet stage. Each point represents a perturbation; the red dashed line indicates the linear fit, and the Spearman correlation coefficient (ρ) and one-sided p-value are shown. (f) Heatmap representing the statistical significance (−log10P) from pathway enrichment analysis of perturbation-specific T2D meta-programs constructed from all cells, endocrine cells or non-endocrine cells. Rows represent individual diabetes-related gene perturbations and columns represent selected diabetes- and endocrine-relevant pathways, including insulin secretion, regulation of insulin secretion, FOXA2/3 transcription-factor networks, MODY, FOXA2 pathway and the gliclazide-action pathway. (g) Enrichment of the top 500 genes from perturbation-specific T2D meta-programs in endocrine cells among genes differentially expressed (Padj < 5%) in islet cells between (i) mSTZ-treated and wild-type (WT) mice, (ii) db/db and WT mice, and (iii) T2D and healthy donors from HPAP. Bars show enrichment odds ratios for individual diabetes-related gene perturbations; error bars represent 95% confidence intervals; the dashed horizontal line denotes an odds ratio of 1. Across perturbations, enrichment for HPAP genes was significantly stronger than enrichment for either db/db or streptozotocin-responsive genes (one-sided paired Wilcoxon P = 3.0e-08 for both db/db and streptozotocin-responsive genes), with mean differences in log odds ratio of 0.85 and 1.18 (95% bootstrap CI, 0.25–1.36 and 0.79–1.52). Source data are provided as a Source Data file.

T2D informativeness of the perturbation meta-programs increased along the differentiation time course. TRS^(ALL)^, corresponding to MetaProg^(ALL)^, increased from *τ*^∗^ =0.30 (p=2.0e-03) at the DE stage to *τ*^∗^=0.83 (p=3.3e-03) at the SC-islet stage (**Figure 6c**); analogous results were observed when using heritability enrichment rather than *τ*^∗^ (**Figure S10a**). The number of perturbations with significant TRS for T2D paralleled this trajectory; among the 24 perturbations analyzed across all four stages, 2, 0, 10, and 15 showed nominally significant (p < 0.05) T2D TRS^(*p*)^ at the DE, PFG, PP and SC-islet stages, respectively (**Figure 6d**). HbA1c TRS profiles showed a correlated trend across stages with T2D TRS (average Spearman *ρ*=0.3 across perturbations) (**Figure S10b**). By comparison, TRS was lower for hepatic traits and substantially weaker for brain and hematologic/immune traits across differentiation (**Figure 6c**). Of the 30 perturbed regulators, 15 were selected based on established roles in monogenic or neonatal diabetes; Perturbscape identified significant T2D TRS for 8 of these 15 genes (53%) at day 18, indicating convergence of monogenic and polygenic diabetes biology within shared islet regulatory programs.

At day 18, atlas-wide T2D TRS^(ALL)^ and heritability enrichment were comparable across all-cell, endocrine and non-endocrine analyses (**Figure S10c**). Notably, per-perturbation TRS^(*p*)^ profiles were more strongly correlated between all cells and endocrine cells than between all cells and non-endocrine cells (Spearman *_ρ_* = 0.35 vs −0.03; **Figure 6e**). Consistent with this preferential endocrine localization, insulin-secretion pathways were significantly enriched in the meta-programs of 16 endocrine and 16 all-cell perturbations, compared with 9 non-endocrine perturbations among 20 perturbations represented in all three analyses (**Figure 6f**). Key perturbations showing this enrichment for endocrine cells include *PDX1, TLE3, OTUD5*, and *FOXA1*. Of these, *PDX1, TLE3,* and *FOXA1* (the latter’s meta-program was also enriched for the *FOXA2/3* network) are established regulators of pancreatic endocrine differentiation, β-cell identity and function^63–66^. By contrast, *OTUD5*, a deubiquitinase, has a less-characterized connection to β-cell developmental or secretory programs.

Finally, we compared top 500 T2D meta-program genes with transcriptional alterations in three islet datasets: human T2D versus healthy islets from HPAP (4,971 genes)^67^, db/db versus healthy mouse islets (7,446 genes)^68^, and multiple low-dose streptozotocin (mSTZ)-treated versus control mouse islets (5,987 genes)^68^. Across perturbations, enrichment was strongest for HPAP human T2D islet differentially expressed genes, with mean odds ratios of 3.06 and 3.05 in all-cell and endocrine analyses, respectively (P ranging from 2.2e-43 to 4.0e-09) (**Figure 6g, S10d**). Enrichment was substantially weaker for genes identified in the db/db model (with human orthologs), with average odds ratios of 1.24 and 1.30 (P = 1.8e-05 to 9.1e-01), and almost absent for the streptozotocin model, with average odds ratios of 1.03 and 0.93 (P = 4.0e-03 to 1.0e+00). These differences remained significant after down-sampling the three differential-expression gene sets to a common size (**Table S3**). Thus, in these comparisons, human perturbation-derived T2D programs more closely recapitulated transcriptional alterations observed in human T2D islets than those from the two mouse models.

Together, these analyses show that Perturbscape resolves the developmental and cellular contexts in which T2D-relevant perturbation programs emerge during β-cell differentiation and identifies programs that converge with transcriptional alterations in human T2D islets.

### Perturbscape applied to *in vivo* brain Perturb-seq resolves autism-relevant perturbations and linked pathways across neuronal subclasses

We applied Perturbscape to PerturbAI, a genome-scale *in vivo* Perturb-seq atlas of the mouse brain in which 1,947 disease-associated genes with broad neuronal expression were perturbed by AAV-delivered CRISPR–Cas9 and profiled by snRNA-seq (**Figure 7a**)^69^. Pooled gRNA libraries were retro-orbitally injected into 74 young (P16) Cas9-transgenic mice, and whole brains were harvested three to four weeks later (P37–P44), yielding 7.7M NeuN+ nuclei mapped onto the Allen Brain Cell (ABC) atlas taxonomy^70^. Because brain-trait heritability is partitioned across neuronal subclasses^16^, we analyzed four subclasses with >1,000 perturbations represented by ≥100 cells: L4/5 IT cortical glutamatergic neurons (L4/5 IT Glut), parvalbumin GABAergic interneurons (Pvalb Gaba), thalamic Prkcd/Grin2c glutamatergic neurons (TH Prkcd/Grin2c Glut), and midbrain glutamatergic neurons (MB Glut). Mouse genes were mapped to human orthologs, and perturbation programs were evaluated against 6 brain-related traits (**Table S2**, average GWAS N = 148,745).

**Figure 7:**
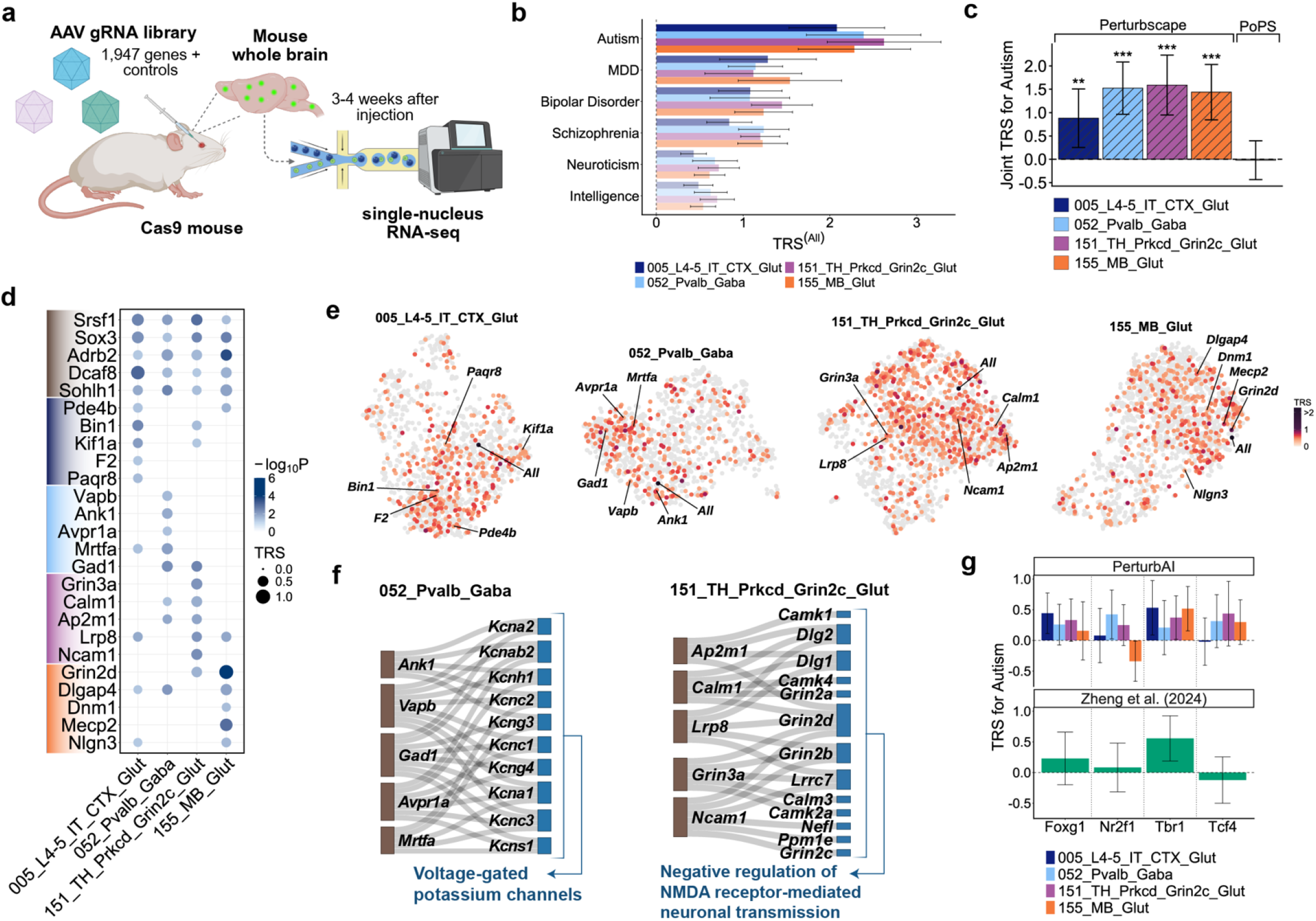
Perturbscape resolves autism-relevant perturbation programs across neuronal subclasses *in vivo*. (a) Schematic of the PerturbAI genome-scale *in vivo* mouse brain Perturb-seq screen. An AAV-delivered gRNA library targeting 1,947 genes together with controls was introduced into Cas9-transgenic mice, and whole-brain nuclei were profiled by single-nucleus RNA sequencing 3–4 weeks after injection. (b) Bar plots showing MetaProg^(ALL)^ trait relevance scores, TRS(ALL), for 6 brain-related traits across 4 neuronal subclasses. Autism shows the strongest heritability relevance across the four neuronal subclasses. Error bars represent 95% confidence intervals. (c) Joint heritability analysis for autism of 5 annotations corresponding to the four subclass-specific Perturbscape MetaProg^(ALL)^ annotations and a single published PoPS annotation. Bars show joint TRS estimates after conditioning on 53 other baseline annotations in the S-LDSC heritability model. Error bars represent 95% confidence intervals and asterisks denote statistical significance (***P < 0.001, **P < 0.01, *P < 0.05). (d) Perturbation-specific autism relevance across neuronal subclasses. Magnitude (TRS, dot size) and significance (−log10P, dot color) of autism disease-heritability information for perturbation-specific meta-programs. We show five perturbations with broadly shared autism TRS across all four neuronal subclasses together with five perturbations showing strong subclass-specific TRS for each of the four subclasses, yielding 25 perturbations in total. The selected perturbations are also highlighted in the UMAP visualization in panel (e). (e) UMAP embeddings of perturbation-specific autism meta-programs within each of the four neuronal subclasses. Perturbations are colored by their trait relevance score (TRS), with non-significant perturbations shown in gray. (f) Sankey diagrams showing pathway-level convergence of autism-relevant perturbations within neuronal subclasses. (Left) In Pvalb GABAergic neurons, perturbations with significant autism TRS whose top 500 meta-program genes were significantly enriched for the voltage-gated potassium channels pathway are linked to the subset of prioritized genes belonging to this pathway. (Right) In Prkcd/Grin2c glutamatergic neurons, perturbations with significant autism TRS whose top 500 meta-program genes were significantly enriched for the “negative regulation of NMDA receptor-mediated neuronal transmission” pathway are similarly linked to the prioritized genes belonging to this pathway. (g) Cross-study comparison of autism-related perturbation relevance across developmentally distinct *in vivo* Perturb-seq datasets. Autism TRS estimates are shown for four transcription-factor perturbations shared between PerturbAI and Zheng et al. (*Foxg1, Nr2f1, Tbr1* and *Tcf4*). Top, perturbation-level autism TRS across the four PerturbAI neuronal subclasses; bottom, corresponding autism TRS from the Zheng et al. developing-cortex Perturb-seq screen. Because the studies differ in perturbation timing, developmental stage and profiled cellular context, this comparison assesses qualitative concordance rather than direct replication. Error bars denote 95% confidence intervals. Source data are provided as a Source Data file.

Autism showed the strongest heritability relevance among the 6 brain-related traits across all 4 neuronal subclasses (**Figure 7b**). TRS^(ALL)^ ranged between 2.1 and 2.6 (p < 2.3e-12), averaging 2.7-fold higher than for the other brain-related traits; the same ordering was observed for heritability enrichment (**Figure S11a**). Autism had the smallest GWAS sample size of the six traits (**Table S2**), so this ordering is unlikely to reflect differences in GWAS statistical power. The stronger autism signal was not explained by disease-based target ascertainment in our analysis (**Figure S11b**). Because perturbations were introduced postnatally and profiled at P37–P44, however, we interpret the autism enrichment cautiously: these experiments may reveal neuronal regulatory programs enriched for autism genetic risk but do not necessarily recapitulate the earlier developmental windows during which many autism-associated genes exert their effects. Given the consistently strong signal across neuronal subclasses, we focused subsequent analyses on autism. In a joint S-LDSC model containing autism-related Perturbscape MetaProg^(ALL)^ programs across neuronal subclasses and PoPS-published, all four MetaProg^(ALL)^ showed joint heritability informativeness (FDR-adjusted p < 0.05) and absorbed the PoPS signal (**Figure 7c**, **S11c**); similar results were observed with PoPS-matched (**Figure S11d**). Meta-programs derived from matched control neurons in each subclass showed substantially weaker or non-significant heritability informativeness than MetaProg^(ALL)^ in both marginal and joint models (**Figure S11e-f**), indicating that the perturbation-derived programs captured additional disease-relevant genetic signal beyond that present in the corresponding baseline neuronal programs.

Perturbation-level autism TRS profiles were only weakly correlated across the four neuronal subclasses (pairwise r = 0.07-0.13), indicating substantial subclass specificity in disease-relevant perturbation responses. **Figure 7d** reports 25 perturbations with divergent TRS profiles across neuronal subclasses. *Srsf1,* haploinsufficiency of which causes a syndromic neurodevelopmental disorder^71^, showed broadly elevated autism TRS across subclasses. *Grin2d* showed the strongest signal in midbrain glutamatergic neurons; de novo variants in *GRIN2D* cause developmental and epileptic encephalopathy with autistic features^72^. *Gad1*, which encodes GAD67, the principal GABA-synthesizing enzyme of inhibitory interneurons^73^ and is recurrently downregulated in ASD postmortem cortex^74^, showed signal in parvalbumin GABAergic interneurons. **Figure 7e** presents the UMAP embedding of the autism perturbation meta-programs for each of the 4 neuronal subclasses, colored by TRS score. High-TRS perturbations converged on distinct neuronal pathways across subclasses (**Figure S11g-h**). In Pvalb GABAergic interneurons, multiple high-TRS perturbation meta-programs converged on voltage-gated potassium-channel genes, including *KCNC1*, *KCNC2*, *KCNC3*, *KCNA1* (**Figure 7f**); this is consistent with the specialized potassium-channel machinery that supports fast-spiking Pvalb interneuron physiology^75^. In thalamic Prkcd/Grin2c glutamatergic neurons, high-TRS perturbation meta-programs converged on NMDA-receptor-mediated neuronal transmission pathway genes, including *GRIN2A*-*D, DLG1/2* and calcium/calmodulin-dependent signaling genes (**Figure 7f**). Collectively, these results suggest that autism-relevant perturbations converge on distinct electrophysiological and synaptic regulatory modules across neuronal subclasses.

Finally, we compared four shared perturbations (*Foxg1*, *Nr2f1*, *Tbr1*, and *Tcf4*) across PerturbAI and an independent developmentally distinct *in vivo* Perturb-seq dataset from Zheng et al.^76^ (in utero delivery at E14.5, profiled at P7) (**Figure 7g**). Because the two studies perturb genes and profile transcriptional consequences at different developmental stages and in different cellular contexts, we treated this as a cross-study comparison rather than a direct replication analysis. *Tbr1* showed significant autism TRS in Zheng et al. and in 2 out of 4 PerturbAI neuronal subclasses (FDR-adjusted p < 0.05), whereas *Tcf4* and *Nr2f1* showed relatively weak signals in both datasets (FDR-adjusted p > 0.12 and 0.07 across subclasses respectively); *Foxg1* showed significant autism TRS specifically in L4/5 IT neurons in PerturbAI (FDR-adjusted p=0.03). Thus, despite substantial experimental and developmental differences, several shared perturbations showed qualitatively concordant autism-relevance patterns across datasets.

Together, these analyses establish that Perturbscape extends to genome-scale *in vivo* brain Perturb-seq, identifies autism as the most heritability-relevant brain trait, and resolves subclass-specific perturbations and downstream pathways associated with its genetic architecture.

## Discussion

Perturbscape provides a general framework for interpreting human disease genetics through transcriptional programs revealed by experimental perturbations. Rather than representing each perturbation by a single differential-expression signature or a small number of shared programs, Perturbscape captures multi-scale regulatory programs within each perturbation and uses human genetics to identify the combinations most relevant to disease. This formulation produces two complementary views of disease biology: MetaProg^(ALL)^ integrates disease-relevant gene programs distributed across an entire perturbation atlas to prioritize disease genes, whereas perturbation-specific meta-programs can assess the extent to which the downstream consequences of a single perturbation are enriched for disease heritability. Perturbscape complements recent work by Ota et al.^15^, which integrates quantitative rare-variant loss-of-function effects with Perturb-seq regulatory measurements to infer directional regulator–program–trait relationships for hematological traits. While the latter approach enables directional mechanistic inference, its reliance on sufficiently powered gene-level loss-of-function associations may limit its applicability across complex diseases and traits for which such associations remain sparse. In contrast, Perturbscape uses common-variant GWAS to systematically identify disease-relevant perturbations and downstream regulatory programs across diverse cellular contexts, without incorporating rare-variant associations into program learning or disease-conditioned meta-program construction. Instead, rare-variant burden associations provide an independent source of genetic validation, demonstrating convergence of common- and rare-variant disease signals within perturbation-induced regulatory programs. Across matched diseases and traits, MetaProg^(ALL)^ provided competitive or improved disease-gene prioritization relative to the integrative multi-tissue PoPS framework^19^ while prioritizing complementary disease genes. This demonstrates that perturbation atlases provide a complementary form of regulatory diversity to steady-state multi-tissue resources by sampling experimentally induced states within disease-relevant cellular systems. Concordance with common- and rare-variant associations, enhancer CRISPRi screens, pharmacologic responses and population trans-eQTLs further supports the utility of these perturbation-derived programs for connecting genetic variation to context-specific cellular biology.

Perturbscape has several downstream applications. First, the perturbation–pathway–disease maps can prioritize perturbations and downstream pathways for therapeutic investigation, as illustrated by the statin-associated programs in HepG2. Second, the multiscale perturbation-induced program atlas provides a structured representation of cellular responses to genetic interventions that could inform the development of virtual-cell^77–79^ and cellular world models^80,81^. In particular, perturbation-derived programs could serve as interpretable response representations for modeling cellular state transitions and predicting responses to unseen perturbations, while disease-conditioned meta-programs could provide genetically anchored objectives or evaluation criteria for assessing whether predicted cellular responses capture disease-relevant regulatory biology. Third, disease-relevant perturbations and genes within their meta-programs could guide subsequent experimental rounds, enabling iterative disease-genetics-guided map–perturb–predict^82^ or lab-in-the-loop^83^ active-learning frameworks. Finally, integrating perturbation-derived programs across multiple Perturb-seq datasets with complementary steady-state and functional annotations could further improve disease-gene prioritization, extending the previous works^19,84^.

The Perturbscape framework has several limitations. First, it is currently formulated primarily for single-perturbation designs, including low-MOI pooled screens and clonally engineered perturbations. In high-MOI screens, co-occurring perturbations complicate attribution of cellular responses to individual targets; extending Perturbscape to this setting will require counterfactual-guided programs that isolate each perturbation while accounting for co-perturbation composition. Second, Perturbscape’s ability to recover a perturbation-induced gene program depends on perturbation coverage, perturbation efficiency, the number of profiled cells and the cellular states represented in a screen. Third, the current framework focuses primarily on transcriptional consequences of single-gene perturbations at discrete endpoints and does not explicitly model time-resolved perturbation dynamics^85^, combinatorial genetic interactions^86^, or perturbations with multi-omic readouts^87,88^. Fourth, enrichment of a perturbation-specific meta-program for disease heritability establishes genetic relevance but should not be interpreted as causal. It does not by itself identify the causal variant or establish that the perturbed gene mediates the association; this is characteristic of most scalable program-to-disease mapping approaches in the literature^10,13,16,19^. Fifth, like PoPS, Perturbscape relies on sufficiently polygenic genome-wide association signal to learn stable relationships between gene programs and disease genetics and may therefore have reduced power for traits with sparse genetic architectures or underpowered GWAS. Sixth, Perturbscape currently summarizes multiple guides targeting the same gene into a gene-level perturbation, which increases cell numbers and robustness to guide-specific technical variation for program learning but can obscure biologically meaningful heterogeneity among guides. Future extensions that model guides hierarchically within genes could address this limitation. Despite these limitations, Perturbscape provides a scalable framework for connecting experimentally measured perturbation-response programs to human disease genetics and for mapping the regulatory pathways through which genetic risk influences cellular phenotypes.

## Methods

### Perturb-seq datasets, preprocessing and QC

We analyzed seven Perturb-seq systems spanning CRISPRi, CRISPR–Cas9 knockout, stem-cell differentiation and *in vivo* brain perturbation experiments (**Table 1**). We restricted analyses to cells or nuclei with an unambiguous assignment to a single genetic perturbation, including dual-guide constructs targeting the same gene where applicable, and excluded cells with assignments to multiple distinct target genes. Non-targeting and safe-targeting guides were retained as controls. Perturbations were analyzed only when represented by at least 100 cells, with a threshold of 200 cells applied to the K562 essential-gene and Jurkat datasets. No threshold was applied to the pancreatic differentiation dataset, in which all targeted lines were retained and the number of perturbations reported reflects those yielding at least one gene program. Guides targeting the same gene were aggregated to define a gene-level perturbation for the primary analyses; guide-level analyses were performed separately where indicated. This design therefore focuses on single-perturbation experiments, including low-MOI pooled CRISPR screens and clonally engineered single-target perturbations.

For all datasets, expression values were normalized to counts per 10,000 per cell and log-transformed (log CP10k + 1). The single exception was the genome-wide K562 Perturb-seq dataset, for which we used the z-score-normalized expression representation provided by the original study^9^. For the genome-wide K562 Perturb-seq dataset, control cells greatly outnumbered the cells assigned to any individual perturbation. We therefore randomly subsampled 10% of control cells, stratified by GEM group so that each group contributed in proportion to its representation in the full control population. This subsampled control set was used for all K562 analyses. Raw count matrices were available for every dataset and were retained alongside the normalized values, providing the input for the models that operate directly on counts. Gene programs were derived within a dataset-specific gene universe capped at 10,000 genes; gene sets below this threshold were retained in full, and larger sets were reduced to the 10,000 most highly variable genes. For the linear contrastive methods, each population is standardized to zero mean and unit variance per gene. For the kernel methods, each population is min–max scaled to the unit interval, satisfying the bounded-domain assumption required by random-feature approximations^89,90^. Before transformation, the workflow checks whether the input already satisfies the required scaling and, if so, passes the values through unchanged to avoid repeated transformation.

For datasets containing multiple cell types, developmental stages or perturbation modalities, analyses were stratified by the relevant biological context, with perturbation programs learned relative to control cells from the corresponding experimental stratum. The pancreatic differentiation dataset comprised 79 barcoded hPSC lines (76 mutant clones and 3 wild-type controls) covering 36 genotypes across 30 pancreatic developmental regulators using CRISPR–Cas9 editing62. We analyzed cells at four developmental stages: definitive endoderm (Day 3), PFG (Day 7), PP (Day 11) and SC-islet (Day 18). Perturbscape was first applied to all cells within each stage and was subsequently repeated within the endocrine and non-endocrine compartments at day 18. For the *in vivo* brain analysis, we used the PerturbAI AAV–CRISPR–Cas9 single-nucleus RNA-seq atlas^69^. Although the original dataset contained nuclei with single and multiple guide assignments, we restricted Perturbscape analysis to nuclei with a single unambiguously assigned perturbation. Analyses were performed separately within 4 Allen Brain Cell atlas neuronal subclasses with sufficient perturbation coverage: layer 4–5 intratelencephalic cortical glutamatergic neurons (L4/5 IT Glut), parvalbumin-positive GABAergic interneurons (Pvalb Gaba), thalamic *Prkcd/Grin2c*-expressing glutamatergic neurons (TH Prkcd/Grin2c Glut), and midbrain glutamatergic neurons (MB Glut). Mouse genes were mapped to their human orthologs before construction of gene programs and subsequent integration with human GWAS data. The independent developing-cortex Perturb-seq dataset of Zheng et al.^76^ was processed analogously and used only for cross-screen validation of the four transcription-factor perturbations shared with PerturbAI. Dataset-specific numbers of cells, genes, perturbations and relatively independent gene programs per perturbation (defined below) are provided in **Table 1**.

Perturbscape is implemented as a modular, workflow-managed computational framework within Snakemake (see **Code Availability**). It takes as input an AnnData object^91^, containing normalized expression as the main matrix of the object (.X). The dataset further includes an observation column (obs_column, default “gene”) which encodes perturbation identity of a cell, and a designated layer for raw unique-molecular-identifier counts without normalization (counts_layer, default “counts”). A single label within this column is nominated as the control population (background_label, default “non-targeting”) to define the background for applying contrastive methods.

### Perturbation program library construction

Perturbscape constructs a multi-resolution library of gene programs describing the transcriptional response associated with each perturbation. Let **X** ∈ ℛ*^N^*^×*G*^ denote the normalized expression matrix for *N* cells and *G* genes; the current implementation therefore assumes that each analyzed cell can be attributed to a single perturbation or control state and is not designed for high-MOI screens in which individual cells carry multiple perturbations. For each perturbation *p*, we define **X**^(*p*)^ as the expression matrix for cells assigned to that perturbation and let **X**^(0)^ correspond to the background population comprising non-targeting and safe-targeting control cells. Program learning is performed separately for each perturbation, thereby preserving perturbation-specific transcriptional structure rather than first aggregating responses across perturbations.

A gene program is represented as a vector over the common gene universe,

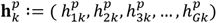

where 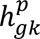 denotes the weight of gene *g* in program *k* associated with perturbation *p*. Depending on the program learner, 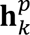 is either a continuous vector of gene weights or a binary vector indicating membership in a set of locally co-modulated genes capturing a coordinated cellular process. The collection of programs learned for perturbation *p* is

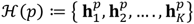

and the complete candidate program space used by Perturbscape is

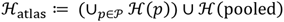

where *P* denotes the set of perturbations analyzed in the screen. ℋ(pooled) represents the set of programs obtained by modeling all perturbed cells as a single target population relative to the same control population. The class of programs per perturbation is determined using several complementary program learners.

#### Differential and canonical latent programs

Direct perturbation-associated transcriptional changes were represented using gene-level differential-expression statistics from a per-gene ordinary least squares regression of normalized expression on a binary perturbation indicator, comparing cells corresponding to perturbation *p* with control cells. The estimated coefficient was retained as the effect size, together with its standard error and P value. We additionally applied differential gene-correlation analysis using DGCA^21^ to capture perturbation-dependent changes in gene co-expression. Pearson correlations were computed separately within the perturbed and control populations across all gene pairs, and the significance of their difference was assessed against a permutation null. Pair-level statistics were then aggregated to the gene level by taking the median across all pairs involving each gene, yielding a continuous per-gene program. To capture dominant multivariate transcriptional structure, we further generated programs using principal-component analysis (PCA) and consensus non-negative matrix factorization (cNMF)^22^. Unlike the contrastive learners, PCA and cNMF were applied to the perturbed cells alone, without reference to the control population, and therefore describe dominant transcriptional structure within the perturbed state rather than variation specifically enriched relative to control. We select 10 latent factors for both PCA and cNMF to capture a broad set of latent states per method.

#### Contrastive learning approaches

We next used contrastive learners to explicitly identify transcriptional variation enriched in cells carrying a given perturbation relative to control cells. For perturbation *p*, the cells in **X**^(*p*)^ were treated as the target population and non-targeting and safe-targeting cells in **X**^(0)^ as the background population. We considered the standard difference-based contrastive PCA (cPCA)^23^, and a ratio-based formulation, which we term ContraPC, as well as nonlinear kernel extensions of both approaches. These methods can be represented within a common contrastive-program formulation. Let *c* ∈ {*t*, *b*} denote the target or background population and let **X***_c_* be the normalized expression matrix for index *c*, **X***_c_* ∈ ℛ^*N_c_*×*G*^. We define **C***_c_* as the empirical gene-gene covariance matrix for the target or background population.

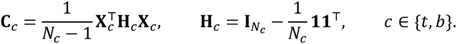

For the kernel formulations, let *X̃_c_* denote the corresponding min-max-scaled matrix, with each entry mapped to [0,1]. We define the gene-gene Gaussian radial-basis-function (RBF) kernel as follows.

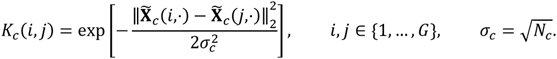

See **Supplementary Note** for details regarding the selection of the kernel parameters. The covariance matrix **C***_c_* captures linear co-variation along global axes, whereas an RBF kernel **K***_c_* captures local and potentially nonlinear similarity structure, including curved or clustered patterns that may not be well represented by any single linear axis. Because direct computation of the RBF kernel matrices becomes expensive as the number of genes grows, we used the Fastfood approximation^90^. Fastfood approximates Gaussian random Fourier features using structured Hadamard transforms, substantially reducing the computational and memory cost (**Supplementary Note**).

To write the linear and kernel versions in a common form, define

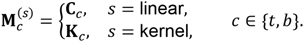

We propose two types of objective functions within the contrastive PCA regime. For the difference-based objective, the cPCA family identifies a direction with high variation in the target population while penalizing variation that is also present in the background:

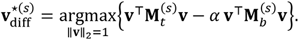

For the ratio-based objective, our proposed ContraPC-based gene programs can be defined as follows.

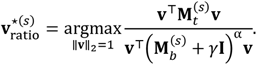

Thus, the four contrastive program learners span two orthogonal methodological choices: difference-versus ratio-based contrast, and linear covariance-versus nonlinear kernel-based representation. In the ContraPC formulation, *γ* is a small regularization term (default 0.001) that stabilizes the denominator when a candidate contrastive direction lies close to the null space of the background matrix, such that 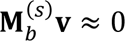. The contrast parameter *α* in both cPCA and ContraPC controls the degree to which variation shared with the background is suppressed. Larger values of *α* specifically emphasize directions showing strong contrast between target and background variation, whereas smaller values retain more subtle perturbation-associated variation that may nevertheless have downstream biological relevance. We therefore evaluated multiple contrast levels, selecting 4 representative values of *α* over a grid, spaced logarithmically from 10^−1^ to 10^3^ for cPCA and linearly over [0, 1] for ContraPC, using a spectral clustering strategy^92^ as proposed in Abid et al.^23^.

#### Deep-generative contrastive representation learning

In addition to the covariance- and kernel-based contrastive learners described above, we applied ContrastiveVI^24^ as a probabilistic deep-generative approach for learning perturbation-specific cellular variation. For each perturbation, ContrastiveVI decomposes gene expression into a salient latent space representing variation specific to perturbed cells and a background latent space representing variation shared with control cells. Both latent spaces were specified with dimension 10. The model was fit to raw counts with library size inferred from the data, using explicit perturbed- and control-cell indices to constrain the salient latent space to variation absent from the control population. Models were trained for up to 500 epochs with early stopping based on the validation objective, and only the salient representation of perturbed cells was retained for downstream analysis.

Unlike the cPCA, ContraPC and corresponding kernel formulations, ContrastiveVI does not provide a closed-form gene-loading vector for each salient latent dimension because the mapping between gene expression and latent space is learned through a nonlinear neural-network encoder. We therefore used ContrastiveVI as a cell-level contrastive representation learner rather than a direct gene-program learner. Gene programs associated with the ContrastiveVI representation were obtained by applying Hotspot to the salient-cell embedding, as described below.

#### Local co-modulated gene programs

Global latent and contrastive components capture dominant axes of transcriptional variation, but perturbation-associated programs may also be restricted to local regions of cellular state space and therefore need not align with a single global component. To identify such locally coordinated gene programs, we applied Hotspot^25^, a graph-based method that detects genes and gene modules exhibiting coordinated local variation across neighboring cells in a predefined cellular embedding. In the Perturbscape framework, Hotspot was applied independently across multiple embeddings generated by the canonical and contrastive representation learners, thereby allowing local gene programs to be detected under complementary representations of perturbation-associated cellular state. Hotspot constructs a neighborhood graph over cells in the cell embedding space generated by each global learner and evaluates each gene for local autocorrelation, testing whether cells that are nearby in the representation space show more similar expression of that gene than expected under the null. Genes exhibiting significant local autocorrelation were subsequently evaluated for pairwise local correlation, and groups of locally co-modulated genes were identified as gene modules using the Hotspot module-detection procedure. Each resulting module ℎ was represented as a binary gene program,

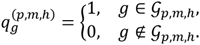

where G*_p_*_,*m*,ℎ_ denotes the set of genes assigned to Hotspot module ℎ for perturbation *p* in embedding method *m*. Hotspot was run with 20 nearest neighbors under exact neighbor search, a false-discovery-rate threshold of 0.05, and a minimum of 30 genes per module, retaining only core module members. When we refer to the program atlas generated by a given method, we include both the continuous gene programs learned directly by that method and the binary Hotspot modules derived from its corresponding cellular embedding. For each perturbation, we combined the programs from all program learners and collapsed redundant ones by greedy leader clustering: programs were visited in random order, and each unassigned program became a representative and absorbed all remaining programs with similarity ≥ 0.7. Similarity was the absolute Pearson correlation across genes for continuous programs and the overlap coefficient (|A∩B| / min(|A|,|B|)) for binary Hotspot gene modules. Because the result depends on visit order, we repeated the procedure over 500 random orders and took the representatives from a permutation achieving the median count; these constitute the relatively independent gene programs for that perturbation. The resulting number of relatively independent gene programs per perturbation is reported for each dataset in **Table 1**.

### GWAS variant and gene-level summary statistics

We analyzed publicly available GWAS variant-level summary statistics for 53 unique diseases and traits with genetic correlation <0.9 that are well-matched to the cell line or tissue of interest (**Table S2**). Each trait passed the filter of being well powered enough for heritability studies (z-score for observed heritability >5 as in previous work including Finucane et al.^93^). We used the summary statistics for SNPs with minor allele count >5 in a 1000 Genomes Project European reference panel^94^. We generated the gene-level summary statistics from the GWAS data using the MAGMA method^26^. MAGMA v.1.08 was run using a 0-kb window around each gene to link SNPs to genes, using all default MAGMA parameters for running the gene-level analysis, and using the 1000 Genomes reference panel for the genotype LD reference.

### Disease-conditioned meta-program construction

Perturbscape identifies the combination of programs in ℋ(*p*) that is most concordant with the genetic architecture of the trait. To do so, we apply the PoPS framework^19^; however, whereas PoPS uses genome-wide multi-tissue expression, pathway, and protein-interaction features, we instead use the perturbation-induced gene programs in ℋ(*p*) as features.

#### Selection of programs

For each perturbation *p* and disease or trait *d*, we first regress MAGMA gene-level association *z*-scores26, **y***_d_*, against each perturbation-induced gene program in ℋ(*p*), accounting for LD-induced correlation among gene-level statistics:

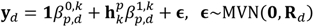

where 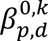 and 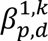 are the intercept and slope parameters and **R***_d_* is a covariance matrix that accounts for projected LD between nearby genes due to the LD among mapped variants in the MAGMA formulation. The model is fit by generalized least squares (GLS). We obtain a p-value for testing the null hypothesis of zero slope against the alternative that 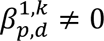 and retain features passing a nominal significance threshold (p<0.05) for the subsequent joint model.

#### Combined meta-program of selected programs

Using the features retained in the previous step, we fit the joint model

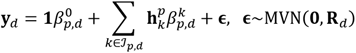

where ℐ*_p_*_,*d*_ ⊆ {1, …, *K*} is the set of indices of the features selected in the previous step. Each feature was standardized to have mean 0 and standard deviation 1 before fitting. Because the model can include many features, we fit it by GLS with L2 regularization. We used a leave-one-chromosome-out (LOCO) scheme: for each held-out chromosome, program coefficients were estimated from genes on the remaining chromosomes, and the fitted model was then applied to genes on the held-out chromosome. Concatenating these predictions across chromosomes yields an optimally weighted, trait-conditioned combination of ℋ(*p*), which we term a “meta-program.” Formally, for gene *g*, the meta-program score is

MetaProg^(*p*,*d*)^ 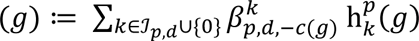 where *c*(*g*) denotes the chromosome containing gene *g*, 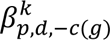 is the GLS coefficient for program *k* estimated while excluding genes on chromosome *c*(*g*), and 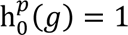. This chromosome-level cross-fitting limits direct reuse of local GWAS association signal during meta-program weight estimation, following the general LOCO strategy used by PoPS and related integrative approaches^19,27,28^.

We analogously construct an atlas-wide meta-program, MetaProg^(ALL,*d*)^, by applying the same procedure to the complete perturbation-program library ℋ*_atlas_*. Because programs learned from different representation methods and perturbations can be correlated, we fit the multivariable model with L2 regularization to stabilize coefficient estimates and limit variance inflation arising from feature collinearity. As a sensitivity analysis, we additionally repeated the analysis after restricting the program library to relatively uncorrelated features (**Supplementary Note**). We observed comparable performance in both *in vitro* and *in vivo* Perturb-seq datasets (**Figure S12a-b**), indicating that the atlas-wide results were not driven by redundancy among highly correlated programs. For the genome-wide K562 screen and the PerturbAI screen, we adopted the atlas-wide meta-program built from these relatively uncorrelated features as the default in the corresponding analyses, as both datasets comprise a large number of perturbations (> 1,000).

For most Perturb-seq datasets, the number of perturbation-derived program features was substantially smaller than the number of features used by PoPS (**Figure S7c**). The genome-wide K562 screen was an exception because the large number of perturbations generated a correspondingly larger program library. For this dataset, prior to multivariable disease-conditioned fitting, we therefore retained the 12,648 programs showing the strongest marginal association with disease gene-level statistics for MCH in Step 1, yielding a feature set of the same size as that used in the matched PoPS model. We evaluated the robustness of this feature-selection strategy in additional sensitivity analyses (**Figure S12a**).

To visualize the perturbation landscape for a given disease, we embedded all per-perturbation and atlas-level meta-programs in a shared UMAP. We first constructed a binary gene-by-meta-program matrix indicating whether each gene ranked among the top 500 by each meta-program’s scores and then fit a UMAP to this matrix using 15 neighbors and the Jaccard index as the distance metric. We used the Python package umap-learn (v0.5.7).

### Trait Relevance Score (TRS) and heritability analysis

Perturbscape quantifies the disease relevance of each perturbation by testing whether genomic regions linked to genes prioritized by its disease-conditioned meta-program are enriched for disease heritability. For each perturbation *p* and disease or trait *d*, we ranked genes according to their meta-program MetaProg^(*p*,*d*)^, and selected the top 500 genes; we do the same for the atlas-wide MetaProg^(ALL,*d*)^. These gene sets were mapped to regulatory elements using the union of ENCODE-rE2G enhancer-gene predictions^29^ across biosamples matched to the relevant tissue (**Table S4**). We then generate a binary 0/1 annotation for 9,991,229 variants with minor allele count >5 from the 1000 Genomes Project European samples^94^, based on whether the variant was linked to a gene in the program. For meta-program *c*, we define the annotation as follows.

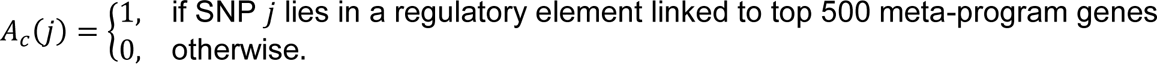

We quantified the heritability informativeness of each meta-program-derived annotation using stratified LD score regression (S-LDSC)^17,30^. For SNP *j*, S-LDSC models the expected GWAS association statistic as

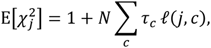

where *N* is GWAS sample size, *τ_c_* is the per-SNP heritability contribution associated with annotation *c*, and

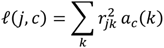

is the LD score of SNP *j* with respect to annotation *c*, with *a_c_*(*k*) denoting the annotation value of SNP *k*. Each Perturbscape annotation was evaluated conditional on 53 baseline coding, conserved and epigenomic annotations similar to Weeks et al.^19^, so that its estimated coefficient reflects heritability information beyond these baseline genomic features. Following previous analyses of cell-type-specific functional annotations^19,95^, we did not additionally condition on the LD-related annotations included in the baseline-LD model^96^ which capture aspects of negative selection and may attenuate cell-type-specific heritability signals^97^.

We then define the Trait Relevance Score (TRS) of perturbation *p* for disease *d* as the standardized S-LDSC effect size, *τ*^∗^, of its corresponding meta-program annotation:

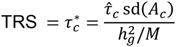

where *τ̂_c_* is the conditional S-LDSC coefficient of the Perturbscape annotation, sd(*A_c_*) is the standard deviation of the annotation across SNPs, 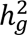 is the SNP heritability of the trait and *M* is the number of SNPs used in the S-LDSC model (equal to 5,961,159 in our analyses). *τ*^∗^ represents the proportionate change in per-SNP heritability associated with a 1 SD increase in the value of the annotation. We use *τ*^∗^ metric as our TRS score because standardizing by annotation variance facilitates comparisons across perturbation-derived annotations with different genomic coverage. The construction of TRS parallels the normalized *τ* statistic used by Benchmarker^28^, and by the subsequent related gene prioritization method PoPS in Weeks et al.^19^. TRS here is more targeted towards the conditional heritability relevance of perturbation-derived disease meta-programs. Standard errors for TRS are estimated using a genomic block-jackknife with 200 blocks.

As a secondary measure, we report the S-LDSC heritability enrichment (ENR),

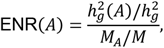

where 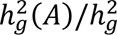 denotes the proportion of SNP heritability attributable to the annotation and *M_A_*/*M* is the proportion of analyzed SNPs contained in the annotation. We use the TRS as the primary statistic because, unlike enrichment, it is less dependent on annotation size, directly quantifies the per-SNP heritability contribution conditional on the baseline model and can be evaluated in joint models containing multiple correlated annotations.

For the joint heritability analysis, we first considered perturbation-specific annotations with marginal evidence of positive heritability informativeness. Among them, we selected MetaProg^(ALL)^ together with the 10 meta-programs that had the highest marginal TRS while being relatively non-overlapping (overlap index of top 500 genes < 0.1) with MetaProg^(ALL)^ and with one another. We then applied stepwise elimination to obtain a reduced set of annotations retaining independent conditional signal. The resulting TRS represents the conditional disease-relevant heritability signal associated with each meta-program after accounting for the other included meta-programs and the 53 baseline annotations. We used analogous joint models when directly comparing atlas-wide Perturbscape annotations, MetaProg^(ALL)^, with PoPS-derived annotations.

To assess calibration of the complete disease-conditioning and heritability-analysis pipeline, we additionally repeated the procedure after shuffling MAGMA gene-level disease statistics before meta-program construction.

### Split-sample reproducibility analysis

To assess whether disease-conditioned meta-programs are reproducible across non-overlapping cohorts—and whether a meta-program learned in one cohort retains its heritability informativeness in another—we performed a split-sample analysis of RBC count. We partitioned the UK Biobank cohort into two non-overlapping, equally sized subsets (N = 215K White British individuals each) and computed GWAS summary statistics for each using REGENIE^34^. We used the UK Biobank Research Analysis Platform for this analysis. From these, we derived MAGMA scores^26^ and, in turn, the K562 Perturb-seq–derived MetaProg^(ALL)^ for each subset. We then performed S-LDSC heritability analysis of the MetaProg^(ALL)^ learned in one subset using the GWAS summary statistics from the other and compared the resulting TRS between the two cross-split assignments.

### Clustering perturbations and characterizing clusters

We partitioned the perturbations into clusters using Leiden clustering. We first constructed a k-nearest-neighbor graph from the binarized meta-programs—binary vectors indicating whether each gene ranked among the top 500 by each meta-program’s scores—using 15 neighbors and the Jaccard index as the distance metric. We then applied Leiden clustering with the Python package leidenalg (v0.10.2), calling find_partition with RBConfigurationVertexPartition as the partition type. Because Leiden clustering results depend on both the resolution parameter and the random seed, we performed a grid search over resolution values of 0.3, 0.4, …, 1.0 and 10 random seeds. For each resulting clustering, we computed the silhouette score using the silhouette_score function from the Python package scikit-learn (v1.5.2) with the Jaccard distance and selected the configuration that maximized it.

To characterize each cluster, we performed pathway enrichment on the genes preferentially ranked highly by that cluster’s meta-programs relative to the others. For each gene, we compared the proportion of meta-programs whose top 500 genes included the focal gene, between meta-programs with significant TRS inside the target cluster and those outside it. We assessed whether the within-cluster proportion exceeded the outside-cluster proportion using a one-sided two-proportion z-test (R function prop.test). We then corrected for multiple testing across all genes using the Benjamini– Hochberg procedure, retained genes with an adjusted p-value below 0.05, and performed pathway enrichment analysis on the resulting gene set.

### Rare-variant burden testing for MCH

Rare-variant, gene-based burden testing for rank-based inverse-normal transformed (RINT) mean corpuscular hemoglobin (MCH) was performed on UK Biobank whole-exome sequencing (WES) data in the White British cohort using REGENIE (v4.1.2)^34^. In Step 1, a whole-genome ridge-regression model was fit using directly genotyped array variants to estimate genome-wide polygenic effects and generate chromosome-specific leave-one-chromosome-out (LOCO) predictions for each individual. These LOCO predictions capture polygenic background, relatedness and population structure while excluding information from the chromosome subsequently tested, thereby limiting proximal contamination. In Step 2, rare coding variants from the WES data were aggregated within genes according to 7 prespecified functional masks: loss-of-function (LoF), missense, synonymous, LoF+missense, LoF+synonymous, missense+synonymous, and LoF+missense+synonymous. For each mask, qualifying variants were collapsed into a gene-level burden genotype and tested for association with MCH using REGENIE’s quantitative-trait association model. For quantitative traits, REGENIE first residualizes both the phenotype and tested genotype with respect to covariates and removes the corresponding chromosome-specific LOCO prediction from the phenotype; association is then tested using linear regression of the residualized phenotype on the burden genotype. Thus, the rare-variant association test is performed conditional on the genome-wide polygenic predictor estimated in Step 1. We consider variants with AAF filter < 0.001; restricting the analysis to variants below 0.1% frequency focuses the test on rare variation and reduces the possibility that burden associations are driven by lower-frequency variants tagging common-variant association signals through linkage disequilibrium^98^. All analyses were adjusted for age, sex, age^2^, age-by-sex interaction, the first 10 genetic principal components, fasting time and genotyping array. To distinguish rare-variant aggregate associations from local common-variant effects, we additionally included as a covariate the genome-wide significant common-variant GWAS signal nearest the transcription start site of the tested gene. We used the UK Biobank Research Analysis Platform for this analysis. The significant burden genes were determined based on Storey’s q-value < 0.05^99^ on the p-values generated across genes within a specific mask.

### Enhancer-level rare-variant association testing

We performed enhancer-level rare-variant association testing for MCH using UK Biobank whole-genome sequencing data (WGS)^100^ and STAARpipeline^46^. We tested each of the 3,670 enhancers assayed in the K562 CRISPRi enhancer screens, aggregating variants with AAF < 0.001 within the genomic interval of each enhancer into a single variant set. STAAR-O was then used to test the association between rare genetic variation within each enhancer and MCH. Enhancer-level rare-variant association analyses were adjusted for age, sex, age², age-by-sex interaction, the first 10 genetic principal components, fasting time and genotyping array, matching the covariates used in the gene-level burden analysis. To distinguish rare-variant aggregate associations from local common-variant effects, we additionally conditioned on the most proximal common-variant GWAS hit within the 1Mb window around the enhancer. STAARpipeline was used to perform multiple set-based rare-variant association tests, including burden^101^, SKAT^102^ and the STAAR-O omnibus test^45^. We report both the burden-test and STAAR-O P values for each enhancer.

### Benchmarking against enhancer-directed CRISPRi screen

For each K562 CRISPRi-significant enhancer, we tested whether its experimentally regulated target genes ranked among the top two genes (out of all genes within 1 Mb of the enhancer) by each of three metrics: (1) PoPS-matched score, (2) MetaProg^(ALL)^, and (3) TRS score. Because the PoPS-matched score and MetaProg^(ALL)^ are defined over program genes whereas the TRS is defined over perturbation genes, we restricted the PoPS-matched score and MetaProg^(ALL)^ to only those genes matching the perturbation genes, ensuring a fair comparison. For each metric, we computed enrichment as the ratio of an observed fraction to a random fraction:

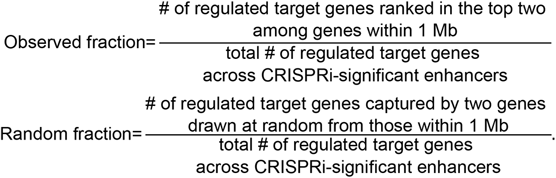

We evaluated this enrichment across 100 random seeds and report the mean and standard error of the resulting 100 values.

### Benchmarking gene prioritization against fine-mapped credible-set–gene links

We first obtained fine-mapped credible-set–gene links as described below, following the approach of refs^19,29^. Fine-mapping was performed on 94 UK Biobank traits (application 31063) using SuSiE^103^ on up to 361,194 individuals of White British ancestry, restricted to variants meeting INFO > 0.8, minor allele frequency > 0.01%, and Hardy–Weinberg equilibrium P > 1 × 10^-10^. Associations were estimated using BOLT-LMM^104^ (for quantitative traits, inverse-rank transformed) or SAIGE^105^ (for binary traits), controlling for principal components, sex, age, and related covariates. SuSiE allowed up to ten causal variants per region (± 1.5 Mb around lead variants), pruned credible sets at r^2^ > 0.25, and excluded both the MHC region (chr. 6, 25–36 Mb) and credible sets containing variants with fewer than 100 minor allele counts.

For gene-linking evaluation, we identified 560 noncoding credible sets located near exactly one gene carrying an independent coding variant (protein-truncating or damaging missense, PIP ≥ 0.5), after deduplication across genetically correlated traits. From the 31 traits or diseases examined in ref^19^, we selected the 12 traits or diseases associated with the cell lines we analyzed, yielding 229 of the 560 links.

To benchmark Perturbscape against PoPS and PoPS-matched for recovery of these 229 silver-standard fine-mapped credible-set–gene links, we ranked, for each credible set, the genes within 500 kb using the MetaProg^(ALL,*d*)^ scores derived from the cell line associated with the corresponding trait (e.g., HepG2 for HDL-C). We then computed the fraction of positive genes (those at loci with an independent, fine-mapped coding variant for the trait) ranked first or among the top two by the MetaProg^(ALL,*d*)^ score and compared this with the fractions obtained using PoPS and PoPS-matched, as well as with a baseline in which genes were ranked at random. As a further comparison, we computed the corresponding top-one and top-two fractions for negative genes -those at loci without an independent, fine-mapped coding variant for the trait -under each score.

### Benchmarking using statin-related in-vivo proteomics and in-vitro stress experiments

In the UKB-PPP cohort of White British individuals^106,107^, we performed differential expression (DE) testing with the Wilcoxon rank-sum test (wilcox.test, R stats package) to identify plasma proteins whose normalized levels differed significantly between 10,619 self-reported statin users and 42,376 non-users. Normalized protein levels were adjusted for age, sex, the first 20 ancestry principal components, BMI, and standing height. We corrected the resulting p-values using the Bonferroni method, for consistency with the DE analyses performed below with the FindMarkers() function in Seurat, and classified the corresponding genes with adjusted P < 0.05 as significantly differentially expressed genes (DEGs).

We similarly performed DE testing with the Wilcoxon rank-sum test to identify genes whose normalized counts differed between HepG2 cells treated with 1 μM atorvastatin and controls. Counts were normalized using the NormalizeData function in the Seurat R package108. Genes were retained if they showed a minimum effect size (|log₂ FC| > 0.1) and sufficient expression (detected in >1% of cells in at least one condition). We applied Bonferroni correction based on the total number of genes and classified genes with adjusted P < 0.05 as DEGs, following the default implementation of the FindMarkers() function in Seurat108.

For three islet datasets—human T2D versus healthy islets from HPAP^67^, db/db versus healthy mouse islets^68^, and streptozotocin-treated versus control mouse islets^68^—we performed DE testing with the Wilcoxon rank-sum test to identify genes whose expression differed significantly between each condition and its baseline. As above, genes were retained if they showed a minimum effect size (|log₂ FC| > 0.1) and sufficient expression (detected in >1% of cells in at least one condition), and we applied Bonferroni correction based on the total number of genes and classified genes with adjusted P < 0.05 as DEGs, following the default implementation of FindMarkers() in Seurat^108^.

### Monocyte trans-eQTL comparison of the Perturbscape results

To assess the enrichment of blood-related trans-regulatory programs identified by Perturbscape in LPS-stimulated THP-1 cells for population-level (cis-eGene, trans-eGene) pairs nominated from QTL analyses of primary monocytes from 432 individuals across four activation states—naive, IFN-γ stimulation, and 2 h or 24 h after LPS stimulation^59^—we treated the perturbed gene and each of the corresponding top 500 disease-conditioned MetaProg^(^*^p^*^,*d*)^ genes as a putative (regulator, target) pair, spanning 23 blood-related traits and 2 modalities. We then tested these Perturbscape (regulator, target) pairs for enrichment of (cis-eGene, trans-eGene) pairs at three levels of increasing specificity. First, we assessed the overall enrichment across perturbations, traits and modalities within each of the four activation states. Second, focusing on the activation state with the strongest overall enrichment (IFN-γ stimulation), we tested whether the trans-eGenes associated with each cis-eGene were enriched among the top 500 MetaProg^(^*^p^*^,*d*)^ genes of the corresponding perturbation, aggregated across the 23 blood-related traits and 2 modalities. Third, focusing on the cis-eGene with the strongest enrichment in the second analysis (*TNF*), we examined the enrichment of its trans-eGenes among the top 500 MetaProg^(^*^p^*^,*d*)^ genes separately for each disease and each modality.

In the first analysis, for each activation state, we performed a stratified enrichment analysis using the Cochran–Mantel–Haenszel (CMH) framework. For each cis-eGene (and its corresponding perturbation *p*), disease *d* and modality *m*, we constructed a 2×2 contingency table quantifying the overlap between the associated trans-eGenes and the top 500 MetaProg^(^*^p^*^,*d*)^ genes. The four cells were defined as follows: *A_p_*_,*d*,*m*_, the number of genes that were both trans-eGenes of the cis-eGene corresponding to *p* and among the top 500 MetaProg^(^*^p^*^,*d*)^ genes of modality *m*; *B_p_*_,*d*,*m*_, the number of trans-eGenes not among the top 500 MetaProg^(^*^p^*^,*d*)^ genes of modality *m*; *C_p_*_,*d*,*m*_, the number of top 500 MetaProg^(^*^p^*^,*d*)^ genes of modality *m* that were not trans-eGenes; and *D_p_*_,*d*,*m*_, the number of genes that were neither trans-eGenes nor among the top 500 MetaProg^(^*^p^*^,*d*)^ genes of modality *m*. We combined these tables across cis-eGenes, the 23 blood-related traits and the 2 modalities, treating each disease– perturbation–modality combination as a stratum, to obtain a common odds ratio:

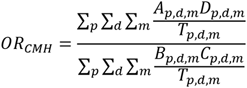

where *T_p_*_,*d*,*m*_ = *A_p_*_,*d*,*m*_ + *B_p_*_,*d*,*m*_ + *C_p_*_,*d*,*m*_ + *D_p_*_,*d*,*m*_ denotes the total number of observations in the corresponding stratum. This stratified approach accounts for differences in the baseline prevalence of trans-eGenes across modalities and diseases, as well as perturbation-specific gene composition, while yielding a global measure of enrichment across the full set of modality–disease–perturbation combinations. The statistical significance of the enrichment—that is, whether the common odds ratio was significantly greater than 1—was evaluated with the CMH test using the mantelhaen.test function in the R stats package.

In the second analysis, we again applied a CMH test to evaluate the overall enrichment across the 23 blood-related traits and 2 modalities, now separately for each cis-eGene (and its corresponding perturbation *p*). The perturbation-specific common odds ratio was computed as:

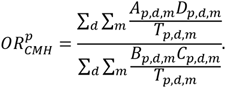

As above, statistical significance—whether the common odds ratio was significantly greater than 1— was assessed with the mantelhaen.test function in the R stats package.

In the third analysis, we quantified the enrichment of trans-eGenes among the top 500 MetaProg^(^*^p^*^,*d*)^ genes separately for each cis-eGene, each blood-related trait and each modality using the odds ratio. Statistical significance—whether the odds ratio was significantly greater than 1—was assessed with Fisher’s exact test using the fisher.test function in the R stats package.

Finally, we carried out an analogous analysis to assess the enrichment of trans-eGenes among the top 500 differentially expressed genes associated with each cis-eGene and its corresponding perturbation in each modality. For each activation state, overall enrichment was evaluated with a CMH test in which the contingency tables were stratified by cis-eGene/perturbation and modality only.

### Statistical assessment of gene-set overlap

To quantify the degree of overlap between two distinct gene sets, we considered two metrics, the Jaccard index and the overlap index, defined as

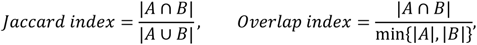

where *A* and *B* denote the two gene sets.

To quantify the overlap between two gene sets relative to the full set of genes, we computed the odds ratio (OR) from a 2×2 contingency table. To ensure numerical stability, we added a pseudo-count of 1 to each cell whenever any off-diagonal entry was zero prior to computing the OR. To assess whether the observed enrichment was statistically significant (i.e., whether the OR was significantly greater than 1), we performed Fisher’s exact test using the R function fisher.test (stats package).

### Pathway enrichment analysis

To identify pathways enriched among the top 500 genes of each meta-program —MetaProg^(*p*,*d*)^ or MetaProg^(ALL,*d*)^ — we performed over-representation analysis using ConsensusPathDB^109^. Unless a specific pathway was of interest (e.g., simvastatin pathway from SMPDB^110^, insulin-related pathways), we restricted the analysis to pathways from Reactome and Wikipathways. We used the default thresholds of a minimum overlap of 2 genes with the input list and a p-value cutoff of 0.01. To correct for multiple testing, we report q-values^99^, which we refer to as adjusted p-values throughout.

## Data Availability

All perturbation-level and atlas-wide meta-program scores for each gene across every cell class and trait, together with all TRS scores and top enriched pathways for the top 500 meta-program genes for each perturbation, cell class, and trait, have been deposited at https://zenodo.org/records/22959154. The MAGMA outputs used in meta-program construction, the results of rare-variant burden testing for MCH performed on UK Biobank whole-exome sequencing (WES) data using REGENIE, and the differential-expression testing results comparing plasma protein levels between statin users and non-users in the UKB-PPP cohort of White British individuals are available at the same Zenodo record.

This study made use of the following publicly available datasets. The K562 CRISPRi Perturb-seq dataset is available at https://doi.org/10.25452/figshare.plus.20029387. The HepG2 and Jurkat CRISPRi Perturb-seq datasets are available from the Gene Expression Omnibus (GEO) under accession GSE264667. The TeloHAEC CRISPRi Perturb-seq dataset is available at the IGVF Data Portal (https://data.igvf.org/matrix-files/IGVFFI8645YMGX/). The THP-1 CRISPRi Perturb-seq dataset is available from GEO under accession GSM6858449, and the THP-1 CRISPR-knockout Perturb-seq dataset under accession GSM6858447. Type 2 diabetes (T2D) datasets are available at https://zenodo.org/records/17824935. The PerturbAI whole-brain CRISPR atlas is available at https://huggingface.co/datasets/perturbai/wholebrain_crispr_atlas. The *in vivo* Perturb-seq dataset from Zheng et al.^76^ is available from GEO under accession GSE249416.

The published PoPS scores are available at https://www.finucanelab.org/data. Data from the Human Pancreas Analysis Program (HPAP) are available at https://hpap.pmacs.upenn.edu. The db/db and streptozotocin-treated mouse islet dataset is available from GEO under accession GSE211799, and the atorvastatin-treated HepG2 cell dataset under accession GSE179347. The monocyte trans-eQTL dataset is available in Supplementary Tables 2 and 3 of Fairfax et al.^59^ (https://www.science.org/doi/10.1126/science.1246949).

ENCODE-rE2G predictions are available on the ENCODE portal (https://www.encodeproject.org). Closest-TSS genome annotations are available at https://github.com/EngreitzLab/CRISPR_comparison/tree/v1.0.0/resources/genome_annotations.

This work used summary statistics from the UK Biobank study (http://www.ukbiobank.ac.uk). The UK Biobank summary statistics used in this paper are available at https://data.broadinstitute.org/alkesgroup/UKBB. The 1000 Genomes Project Phase 3 data are available at ftp://ftp.1000genomes.ebi.ac.uk/vol1/ftp/release/20130502. The baseline annotations are available at https://data.broadinstitute.org/alkesgroup/LDSCORE.

Supplementary Tables S1–S4 accompany this paper. **Table S1** lists the gene-program learners used in Perturbscape, with their references, software and source code. **Table S2** lists the complex traits and diseases used in the disease-informativeness analysis, with GWAS sample sizes and data sources. **Table S3** reports the enrichment of T2D meta-program genes against down-sampled differentially expressed genes in mouse and human islet datasets. **Table S4** lists the ENCODE-rE2G biosamples and the tissues they were matched to for enhancer-gene mapping.

Source data are provided with this paper.

## Code Availability

Perturbscape was implemented in Python 3.11 and orchestrated with Snakemake 9.15. Core numerical dependencies comprised NumPy, SciPy, scikit-learn, scikit-learn-extra (kernel modules only), pandas, and statsmodels, with Scanpy and AnnData used for single-cell data handling^91^, hotspotsc for spatial autocorrelation analysis, cnmf for consensus factorization, and scvi-tools for variational contrastive inference^111^. Differential co-expression analysis was performed in R using DGCA, with sparse matrix handling via Matrix and data.table. Complete, version-pinned environment specifications for each module are distributed with the source code.

Source code is available at https://github.com/Deylab999MSKCC/perturbscape

cPCA: https://github.com/abidlabs/contrastive

ContrastiveVI: https://github.com/suinleelab/contrastiveVI

cNMF: https://github.com/dylkot/cNMF

Hotspot: https://hotspot.readthedocs.io/en/latest/

sc-linker: https://github.com/kkdey/GSSG, https://github.com/karthikj89/scgenetics

LDSC software: https://github.com/bulik/ldsc/wiki

PoPS: https://github.com/FinucaneLab/pops

REGENIE software for performing burden

WES: https://rgcgithub.github.io/regenie/

Large scale genetic analysis using the DNAnexus platform: https://platform.dnanexus.com/

ENCODE-rE2G enhancer-gene linking: https://www.encodeproject.org/software/distal-regulation-encode_re2g/

Seurat: https://satijalab.org/seurat/

## Acknowledgements

We thank all members of the Dey laboratory. We also thank the Impact of Genomic Variation on Function (IGVF) DACC and Network Working Group for helpful discussions.

## Supplementary Figures

**Supplementary Figure 1:**
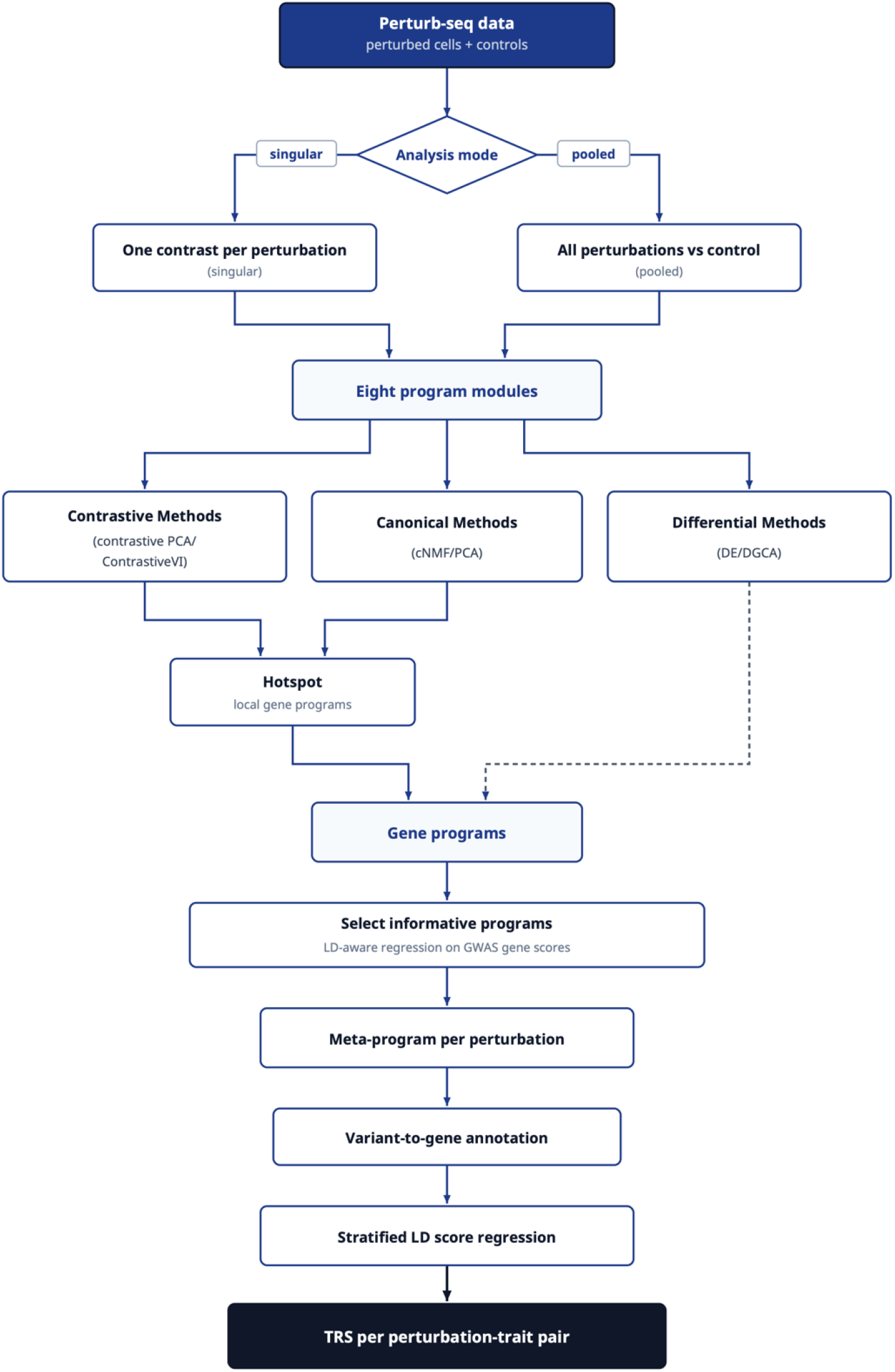
The Perturbscape workflow. Perturb-seq data are analyzed either singly (one contrast per perturbation) or pooled (all perturbations against control). Eight modules define gene programs. Contrastive (cPCA, ContraPC and their kernel forms) and canonical (PCA, cNMF) methods yield programs directly, as gene loadings or spectra, and also pass their cell-level embeddings to Hotspot, which derives further programs from genes co-varying across that space; ContrastiveVI contributes through Hotspot alone, DE/DGCA directly only (dashed arrow). Per GWAS trait, informative programs are selected by LD-aware regression on gene-level association scores, combined into one meta-program per perturbation, converted to SNP annotations via variant-to-gene links and scored by stratified LD score regression, giving a trait relevance score (TRS) per perturbation–trait pair.

**Supplementary Figure 2:**
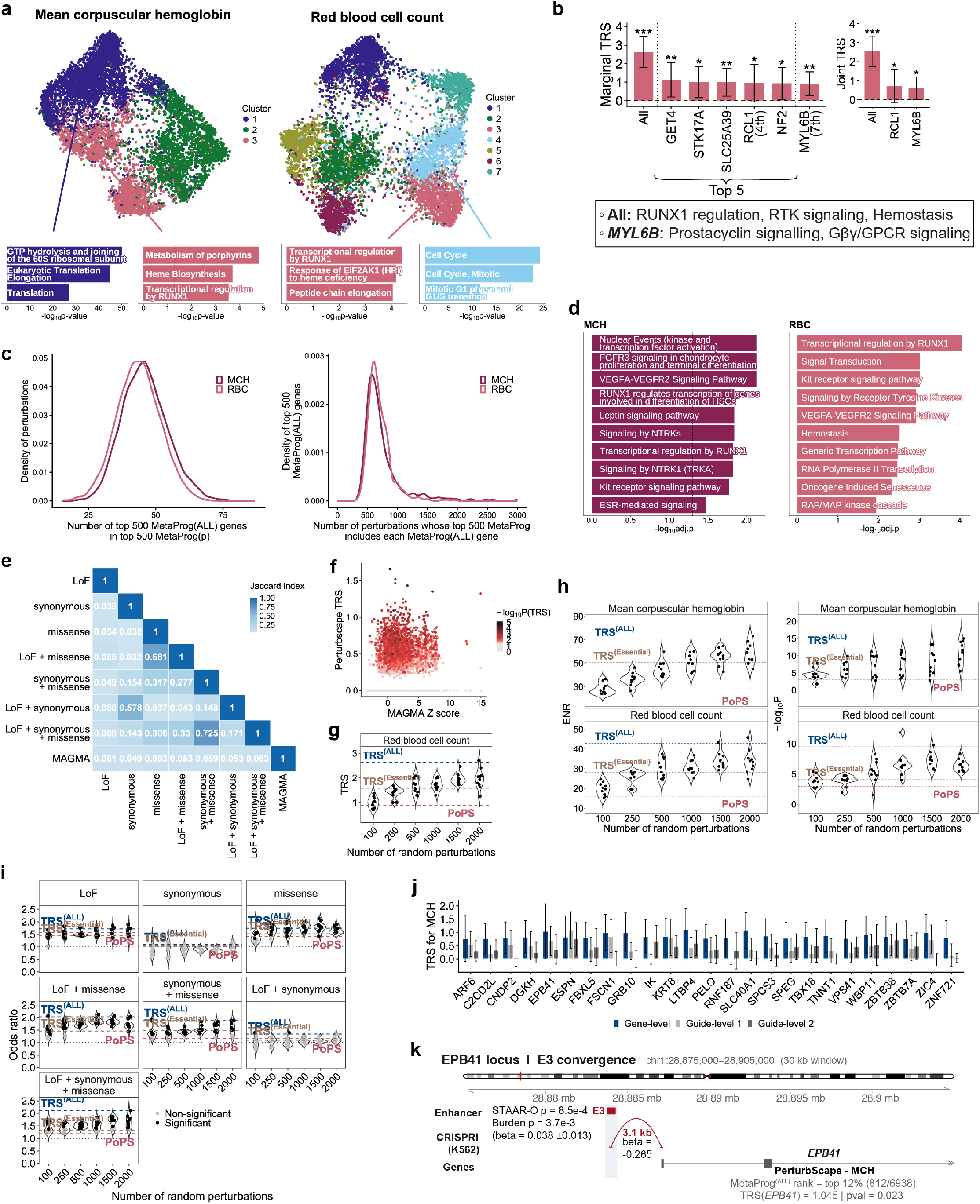
Additional characterization and validation of Perturbscape disease meta-programs for erythroid traits in K562 screen. (a) UMAP embedding of perturbation-specific meta-programs from the K562 whole-genome Perturb-seq screen for mean corpuscular hemoglobin (MCH; left) and red blood cell count (RBC; right), colored by unsupervised cluster assignment (3 clusters for MCH, 7 clusters for RBC). Bar plots below each embedding show representative enriched pathways for selected clusters. (b) Bar plots showing the marginal TRS of MetaProg^(ALL)^ and 6 individual perturbations with high TRS for RBC (left; the top 5—*GET4*, *STK17A*, *SLC25A39*, *RCL1* and *NF2*—together with *MYL6B*, ranked 7th). Joint heritability analysis identified 3 independently informative annotations, MetaProg(ALL), MetaProg(*RCL1*) and MetaProg(*MYL6B*) (right). All analyses were conditioned on 53 baseline annotations comprising coding, conserved and epigenomic annotations. Error bars represent 95% confidence intervals and asterisks denote statistical significance (***P < 0.001, **P < 0.01, *P < 0.05). Top enriched pathways of MetaProg^(ALL)^ and MetaProg(*MYL6B*) are highlighted in the box. (c) Density plots characterizing the overlap between the MetaProg^(ALL)^ and individual perturbation meta-programs, shown for MCH and RBC. Left: distribution, across perturbations, of the number of top-500 MetaProg^(ALL)^ genes recovered within each perturbation’s top-500 MetaProg^(*p*)^. Right: distribution, across the top-500 MetaProg^(ALL)^ genes, of the number of perturbations whose top-500 MetaProg^(*p*)^ includes each gene. (d) Top enriched pathways of MetaProg^(ALL)^ for MCH (left) and RBC (right); the x-axis shows −log10 adjusted p-value. (e) Pairwise Jaccard index between the gene sets defined by the different rare-variant annotation masks—loss-of-function (LoF), missense and synonymous variants and their combinations—and MAGMA gene set for MCH. Color indicates the Jaccard index. (f) Scatter plot comparing perturbation-level TRS (y-axis) with MAGMA z-scores (x-axis) for MCH. Each point represents a perturbation, colored by significance of TRS (−log_10_P). (g) Violin plots showing the distribution of TRS estimates for MetaProg^(ALL)^ constructed from random subsets of 100–2,000 perturbations for RBC; for each subset size, 10 random perturbation subsets were sampled. Dashed lines indicate the values for MetaProg^(ALL)^ from the full 7,991-perturbation screen (blue), MetaProg^(ALL)^ from an independent screen of 414 essential-gene perturbations (gold), and PoPS gene prioritization (red). (h) Violin plots showing the distribution of heritability enrichment (left) and significance of TRS (−log10P; right) for MetaProg^(ALL)^ constructed from random subsets of 100–2,000 perturbations for MCH or RBC; for each subset size, 10 random perturbation subsets were sampled. Dashed lines indicate the values for MetaProg^(ALL)^ from the full 7,991-perturbation screen (blue), MetaProg^(ALL)^ from an independent screen of 414 essential-gene perturbations (gold), and PoPS gene prioritization (red). (i) Odds ratio of rare-variant gene recovery across all annotation masks (the seven combinations of LoF, missense and synonymous variants) for MetaProg^(ALL)^ constructed from random subsets of 100– 2,000 perturbations for MCH. Points are colored by nominal significance (one-sided Fisher’s exact test; black, significant; gray, non-significant). Dashed lines indicate the reference odds ratios for MetaProg^(ALL)^ from the full 7,991-perturbation screen (blue), MetaProg^(ALL)^ from the independent 414 essential-gene screen (gold) and PoPS (red); the dotted line denotes an odds ratio of 1. (j) Bar plot of the TRS for MCH of 25 high-TRS perturbations with two guides represented by ≥100 cells each, computed at the gene level (all guides aggregated; dark blue) and separately for two individual guides (guide-level 1 and guide-level 2; gray). Error bars represent 95% confidence intervals. (k) Convergence of rare-variant burden, enhancer perturbation and Perturbscape evidence at the *EPB41* locus. Rare-variant burden-level association at enhancer E3 (chr1:28,875,000–28,905,000 window; STAAR-O P = 8.5e-4, burden P = 3.7e-3, beta = 0.038 ± 0.013) coincides with a K562 CRISPRi enhancer–gene link to *EPB41*, located 3.1 kb from the enhancer (beta = −0.265). Perturbscape independently prioritizes *EPB41* for MCH, with MetaProg^(ALL)^ ranking the gene in the top 12% and *EPB41* CRISPRi perturbation showing a significant TRS (TRS(*EPB41*) = 1.045, P = 0.023). Source data are provided as a Source Data file.

**Supplementary Figure 3:**
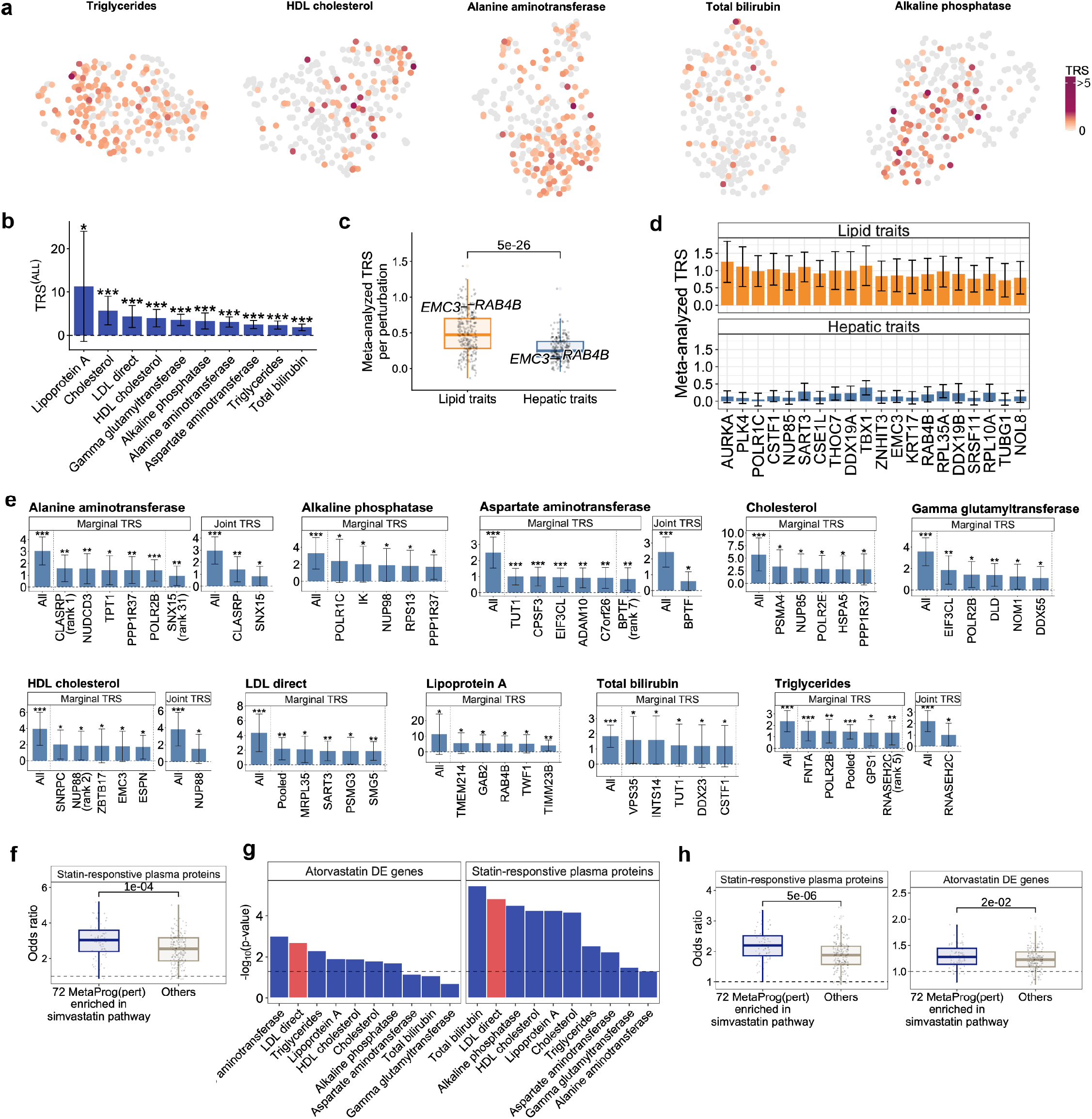
Additional characterization and extended statin-response validation of HepG2 perturbation meta-programs. (a) UMAP visualization of perturbation-specific meta-programs from the HepG2 essential-gene Perturb-seq screen (n = 255) for the remaining five lipid and hepatic traits not shown in **Figure 3a**: Triglycerides, HDL cholesterol, Alanine aminotransferase, Total bilirubin and Alkaline phosphatase. Each perturbation is colored by its trait relevance score (TRS), with non-significant perturbations shown in gray. (b) Bar plot showing the TRS^(ALL)^ for each of 10 lipid and hepatic quantitative traits, ordered by descending TRS(ALL). Asterisks denote significance of TRS^(ALL)^ (*P < 0.05, **P < 0.01, ***P < 0.001); error bars represent 95% confidence intervals. (c) Box plots showing the distribution of meta-analyzed perturbation-level TRS for lipid traits versus hepatic traits. Each point represents a HepG2 essential-gene perturbation, and representative perturbations (*EMC3*, *RAB4B*) are labeled. Perturbations show significantly higher meta-analyzed TRS for lipid than for hepatic traits (one-sided paired t-test; P = 5e-26). (d) Bar plots showing, for the top perturbations ranked by difference in meta-analyzed TRS between lipid and hepatic trait-categories, their TRS for lipid traits (top) and the TRS of the same perturbations for hepatic traits (bottom). Error bars represent 95% confidence intervals. (e) Marginal and joint TRS for each of the 10 lipid and hepatic traits. For each trait, bar plots show the marginal TRS of MetaProg^(ALL)^ and the individual perturbations with the highest TRS (left) and, when the joint model identified annotations other than MetaProg^(ALL)^ as independently informative, the joint TRS of the independently informative annotations (right). All analyses were conditioned on the baseline annotations. Asterisks denote significance (*P < 0.05, **P < 0.01, ***P < 0.001); Error bars represent 95% confidence intervals. (f) Box plot showing the enrichment odds ratio for overlap between the top 500 genes of the direct LDL cholesterol meta-programs of the 72 simvastatin-response-associated perturbations and top 500 plasma proteins showing the strongest statin-use associations, compared with all other perturbations (“Others”). Significance (P=1e-04) was assessed using a one-sided two-sample t-test. The dashed line denotes an odds ratio of 1. (g) Per-trait significance (−log10 p-value) of the difference in the odds ratio for overlap between the top 500 genes of the meta-programs for each trait of the 72 simvastatin-response-associated perturbations and atorvastatin-responsive differentially expressed genes in HepG2 cells (left) or statin-responsive plasma proteins (right), compared with all other perturbations (“Others”). Traits are ordered by decreasing significance; direct LDL cholesterol (red) ranks among the most significant for both experiments. Significance was assessed using a one-sided two-sample t-test. The dashed line denotes the significance threshold (P = 0.05). (h) Box plot showing the enrichment odds ratio for overlap between the top 500 genes of the direct LDL cholesterol meta-programs of the 72 simvastatin-response-associated perturbations and statin-responsive plasma proteins (left; P=5e-06) or atorvastatin DE genes (right; P=2e-02), compared with all other perturbations (“Others”), where the known simvastatin-pathway genes were removed from each meta-program. Significance was assessed using a one-sided two-sample t-test. The dashed line denotes an odds ratio of 1. Source data are provided as a Source Data file.

**Supplementary Figure 4:**
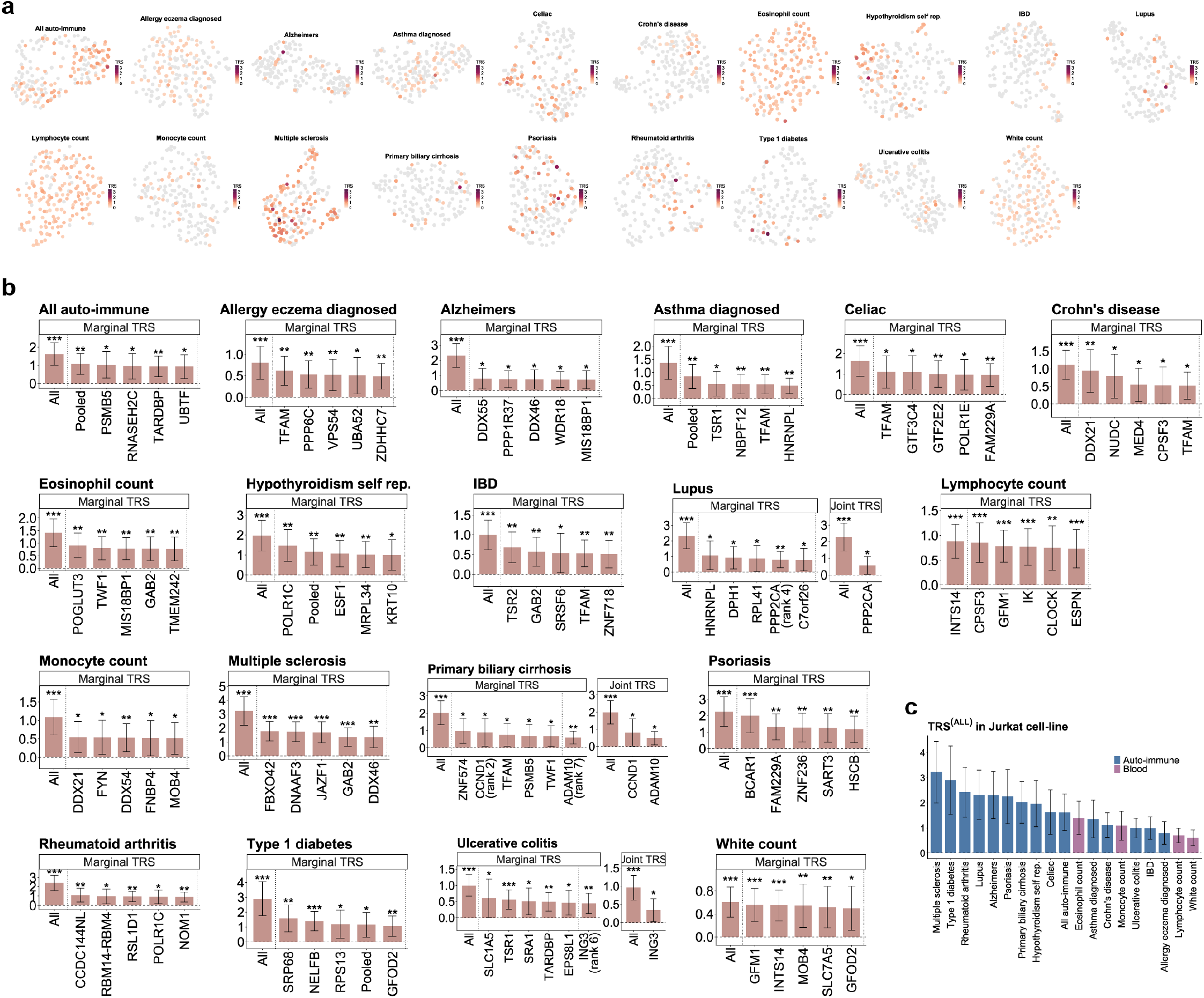
Perturbscape analysis for autoimmune and blood traits in Jurkat Perturb-seq screen. (a) UMAP visualization of perturbation-specific meta-programs from a Jurkat essential-gene Perturb-seq screen for 19 autoimmune and blood-biomarker traits. Each perturbation is colored by its trait relevance score (TRS), with non-significant perturbations shown in gray. (b) Marginal and joint TRS for each of the 19 autoimmune and blood traits. For each trait, bar plots show the marginal TRS of MetaProg^(ALL)^ and the individual perturbations with the highest TRS (left) and, when the joint model identified annotations other than MetaProg^(ALL)^ as independently informative, the joint TRS of the independently informative annotations (right). Asterisks denote significance (*P < 0.05, **P < 0.01, ***P < 0.001); error bars represent 95% confidence intervals. (c) Bar plot showing the TRS(ALL), for each trait in the Jurkat screen, ordered by descending TRS^(ALL)^ and colored by trait category (autoimmune, blue; blood, pink). Error bars represent 95% confidence intervals. Source data are provided as a Source Data file.

**Supplementary Figure 5:**
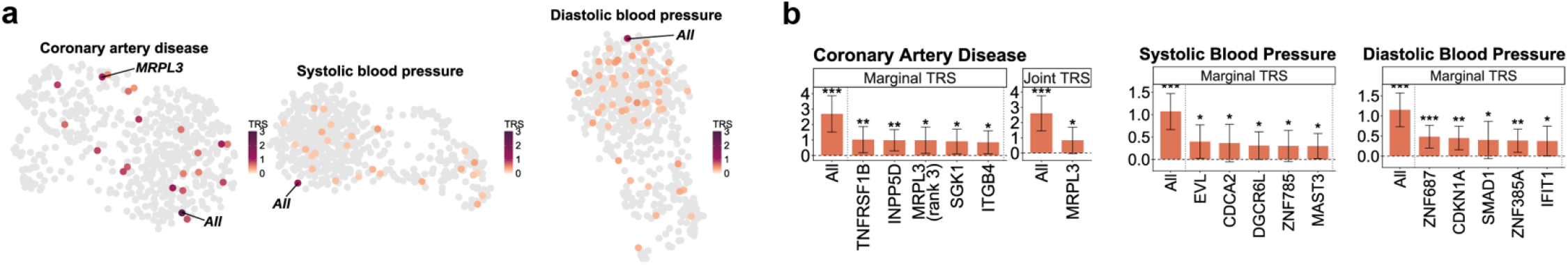
Perturbscape analysis for cardiovascular traits in TeloHAEC Perturb-seq screen. (a) UMAP visualization of perturbation-specific meta-programs from a TeloHAEC Perturb-seq screen for three cardiovascular traits: coronary artery disease, systolic blood pressure and diastolic blood pressure. Each perturbation is colored by its trait relevance score (TRS), with non-significant perturbations shown in gray. (b) Marginal and joint TRS for the three cardiovascular traits. For each trait, bar plots show the marginal TRS of MetaProg^(ALL)^ and the individual perturbations with the highest TRS (left) and, when the joint model identified annotations other than MetaProg^(ALL)^ as independently informative, the joint TRS of the independently informative annotations (right). Asterisks denote significance (*P < 0.05, **P < 0.01, ***P < 0.001); error bars represent 95% confidence intervals. Source data are provided as a Source Data file.

**Supplementary Figure 6:**
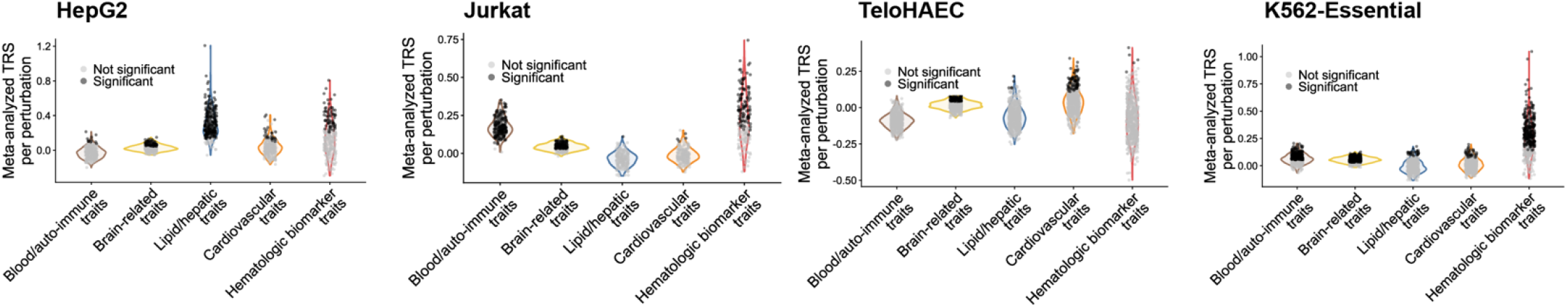
Cell-type-matched specificity of meta-analyzed perturbation-level TRS across four Perturb-seq screens. Violin plots showing the distribution of meta-analyzed perturbation-level TRS across traits within each of five broad trait categories—blood/autoimmune, brain-related, lipid/hepatic, cardiovascular and hematologic biomarker traits—for four Perturb-seq screens: HepG2, Jurkat, TeloHAEC and K562-Essential. Each point represents a perturbation; significant perturbation–trait-category associations are shown in black and non-significant associations in gray. Source data are provided as a Source Data file.

**Supplementary Figure 7:**
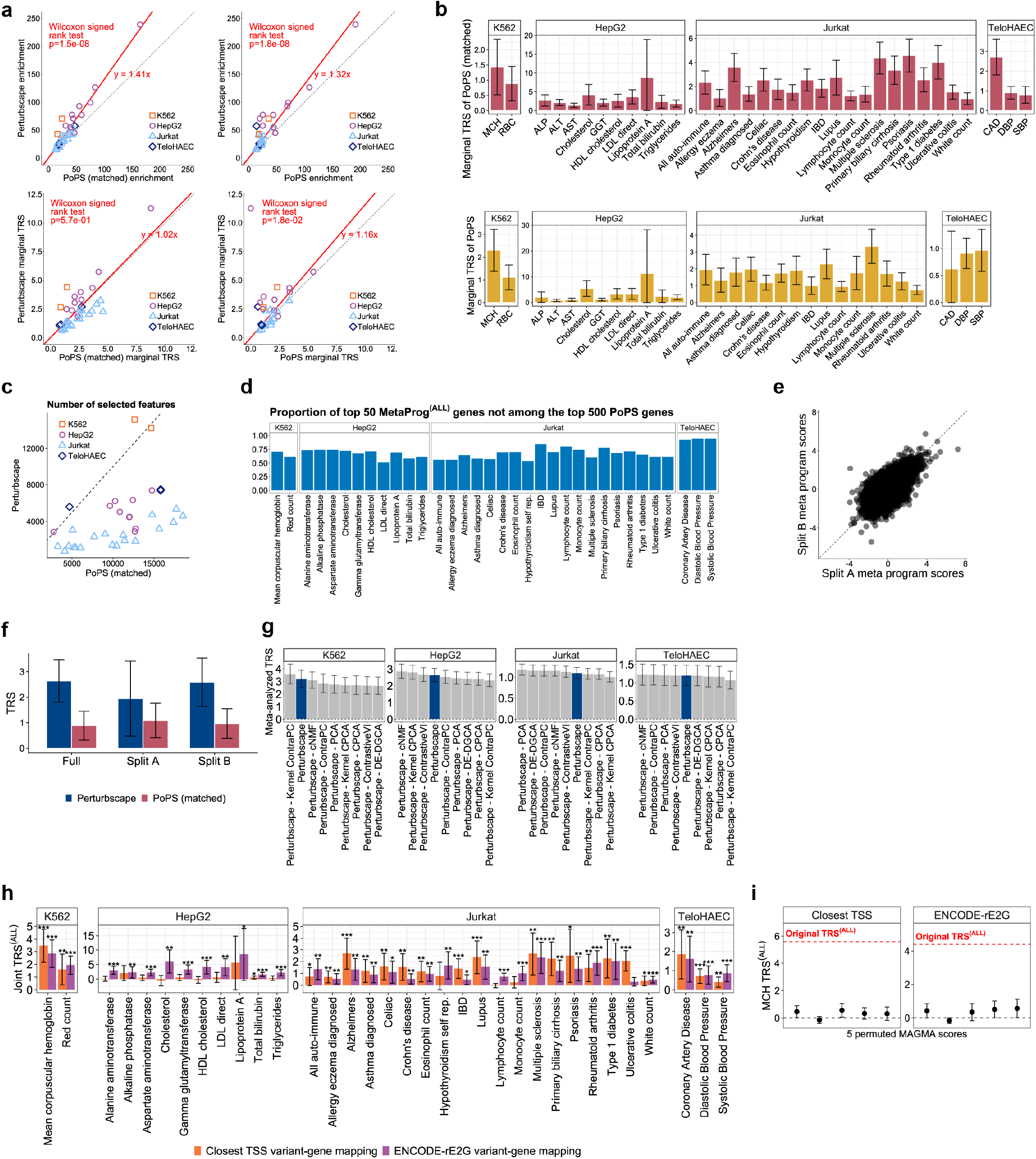
Additional benchmarking of Perturbscape relative to PoPS and across methodological choices. (a) Scatter plots comparing Perturbscape with PoPS across traits from all four Perturb-seq systems (K562, HepG2, Jurkat and TeloHAEC). Perturbscape heritability enrichment is compared with that of PoPS trained on the matched gene universe (first) and published PoPS (second), and Perturbscape marginal TRS is compared with the corresponding marginal TRS of matched PoPS (third) and published PoPS (fourth). Each point represents a trait, colored by cellular system; the dotted line denotes y = x and the solid red line the fitted regression (slope indicated) with zero intercept. P-values are from a one-sided Wilcoxon signed-rank test. (b) Bar plots showing the marginal TRS of PoPS trained on the perturbation-gene universe (“PoPS (matched)”, left) and of published PoPS (“PoPS”, right) for each trait within the four cellular systems. Error bars represent 95% confidence intervals. (c) Scatter plot comparing the number of selected features (programs) in the Perturbscape MetaProg^(ALL)^ versus the matched PoPS model for each trait, colored by cellular system. The dotted line denotes y = x. (d) Bar plot showing, for each trait across the four cellular systems, the proportion of the top 50 Perturbscape MetaProg^(ALL)^ genes that are not among the top 500 PoPS (matched) genes. (e) Scatter plot showing the reproducibility of MetaProg^(ALL)^ gene scores between two independent random splits (Split A and Split B) of the population cohorts; each point represents a gene. The dotted line denotes y = x. (f) Bar plot showing the TRS estimated from the full dataset (Full) and from each of two independent data splits (Split A, Split B) for Perturbscape MetaProg^(ALL)^ (blue) and matched PoPS (red), for RBC. Error bars represent 95% confidence intervals. (g) Ablation analysis of representation learners. Bar plots show the meta-analyzed TRS of the full Perturbscape MetaProg^(ALL)^ (blue) and of MetaProg^(ALL)^ reconstructed after removing each individual representation learner in turn (“Perturbscape − <learner>”; gray), for each of the four cellular systems. Bars are ordered by meta-analyzed TRS. Error bars represent 95% confidence intervals. (h) Bar plots comparing the joint TRS^(ALL)^ obtained using closest-TSS (orange) versus ENCODE-rE2G (purple) variant-to-gene mapping for each trait across the four cellular systems. Asterisks denote significance (*P < 0.05, **P < 0.01, ***P < 0.001); error bars represent 95% confidence intervals. (i) MCH TRS^(ALL)^ computed from five independent permutations of the MAGMA gene-level disease scores (points) is shown for closest-TSS (left) and ENCODE-rE2G (right) variant-to-gene mapping; the red dashed line indicates the original (unpermuted) TRS(ALL). Error bars represent 95% confidence intervals. Source data are provided as a Source Data file.

**Supplementary Figure 8:**
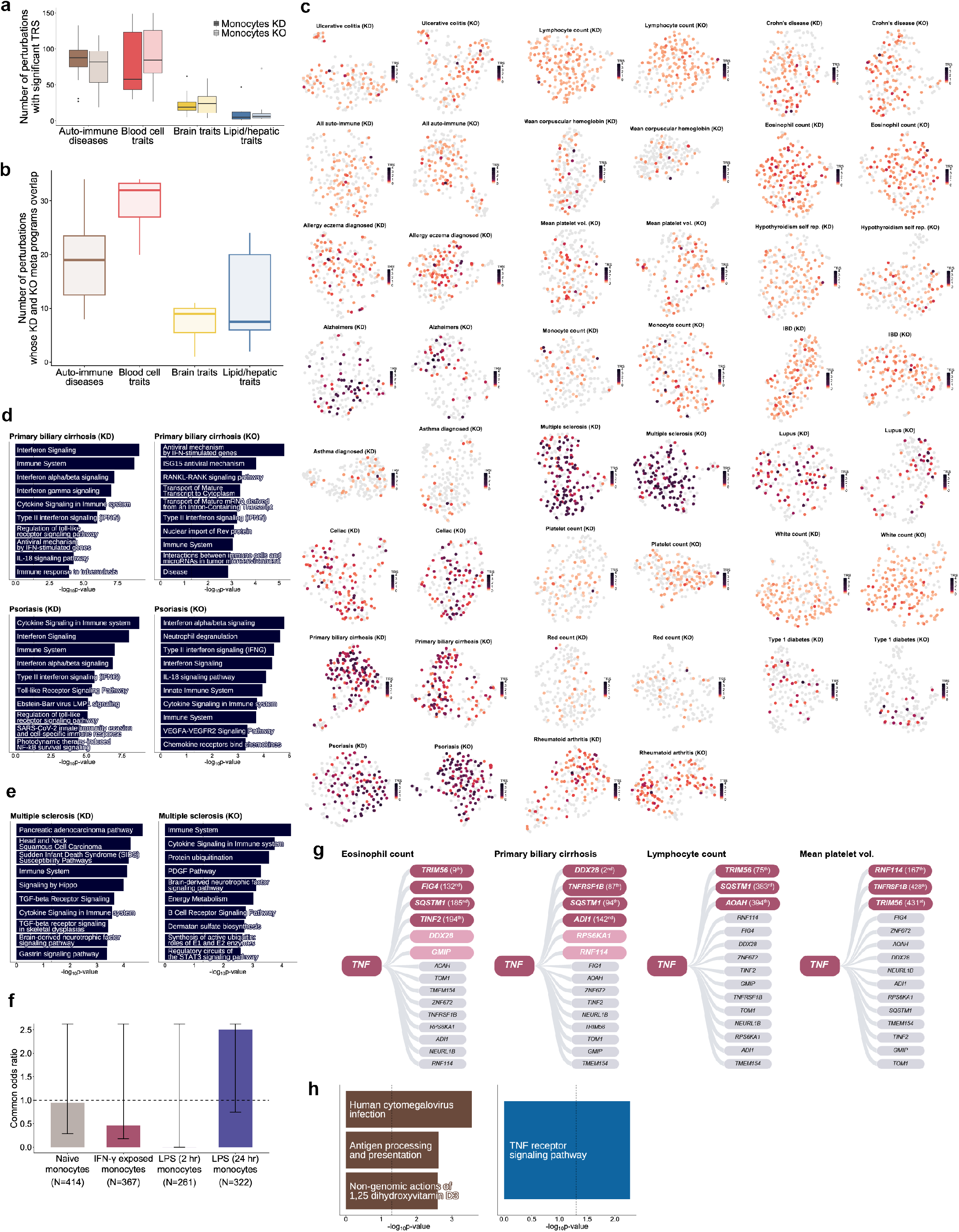
Cross-modality (KD vs KO) reproducibility of monocyte perturbation programs and population trans-eQTL validation. (a) Box plots showing the number of perturbations with significant TRS, grouped into four broad trait categories (autoimmune diseases, blood cell traits, brain-related traits and lipid/hepatic traits) and shown separately for CRISPRi knockdown (Monocytes KD, dark) and Cas9 knockout (Monocytes KO, light). Each box summarizes the distribution, across traits within the category, of the number of significant perturbations (Box: median, 25th/75th percentiles; whiskers: 1.5×interquartile range; points: outliers). (b) Box plots showing, for each trait across the four broad trait categories, the number of perturbations for which the top KD and KO meta-programs overlap, quantifying the reproducibility of perturbation-level signal across modalities. Overlap was assessed using a one-sided Fisher’s exact test, with p < 0.05 considered significant. Each box summarizes the distribution across traits within the category (Box: median and 25th/75th percentiles; whiskers: 1.5×interquartile range). (c) UMAP visualization of perturbation-specific meta-programs for each immune and blood trait analyzed in the monocyte screens, shown separately for KD and KO. Each perturbation is colored by its trait relevance score (TRS), with non-significant perturbations shown in gray (complementing **Figure 5b**). (d) Top enriched pathways (−log10p-value) of the *CCR6* meta-programs for primary biliary cirrhosis (top) and psoriasis (bottom), shown separately for KD and KO. *CCR6*, which encodes a chemokine receptor central to Th17-cell trafficking, showed reproducible disease relevance across KD and KO for primary biliary cirrhosis and psoriasis; these diseases have been linked to Th17-associated inflammatory responses, including IL-23–Th17 signaling. (e) Top enriched pathways (−log10p-value) of the *STK4* meta-programs for multiple sclerosis, shown separately for KD and KO. *STK4* (*MST1*), a Hippo-pathway serine/threonine kinase involved in lymphocyte survival, trafficking and cytokine signaling, showed strong disease relevance across modalities for multiple sclerosis; its multiple-sclerosis meta-program was enriched for cytokine signaling in immune system. (f) Common enrichment odds ratio (Cochran–Mantel–Haenszel test) of DE-derived regulatory links with population trans-eQTLs across four primary-monocyte activation states (naive, N = 414; IFN-γ-exposed, N = 367; LPS 2 h, N = 261; LPS 24 h, N = 322). The dashed line denotes an odds ratio of 1 and error bars represent 95% confidence intervals. (g) Convergence of *TNF*-associated population trans-eQTLs with the *TNF* perturbation meta-program for four additional traits: eosinophil count, primary biliary cirrhosis, lymphocyte count and mean platelet volume (complementing monocyte count in **Figure 5g**). For each trait, the 15 trans-eGenes linked to *TNF* as a cis-eGene in IFN-γ-exposed primary monocytes are ordered by their meta-program scores; genes among the top 500 *TNF* meta-program genes are highlighted, with ranks indicated. (h) Pathway enrichment (−log10p-value) of the cis-eGene (left) and trans-eGene (right) sets enriched in Perturbscape-derived trans-regulatory links across traits and modalities. Source data are provided as a Source Data file.

**Supplementary Figure 9:**
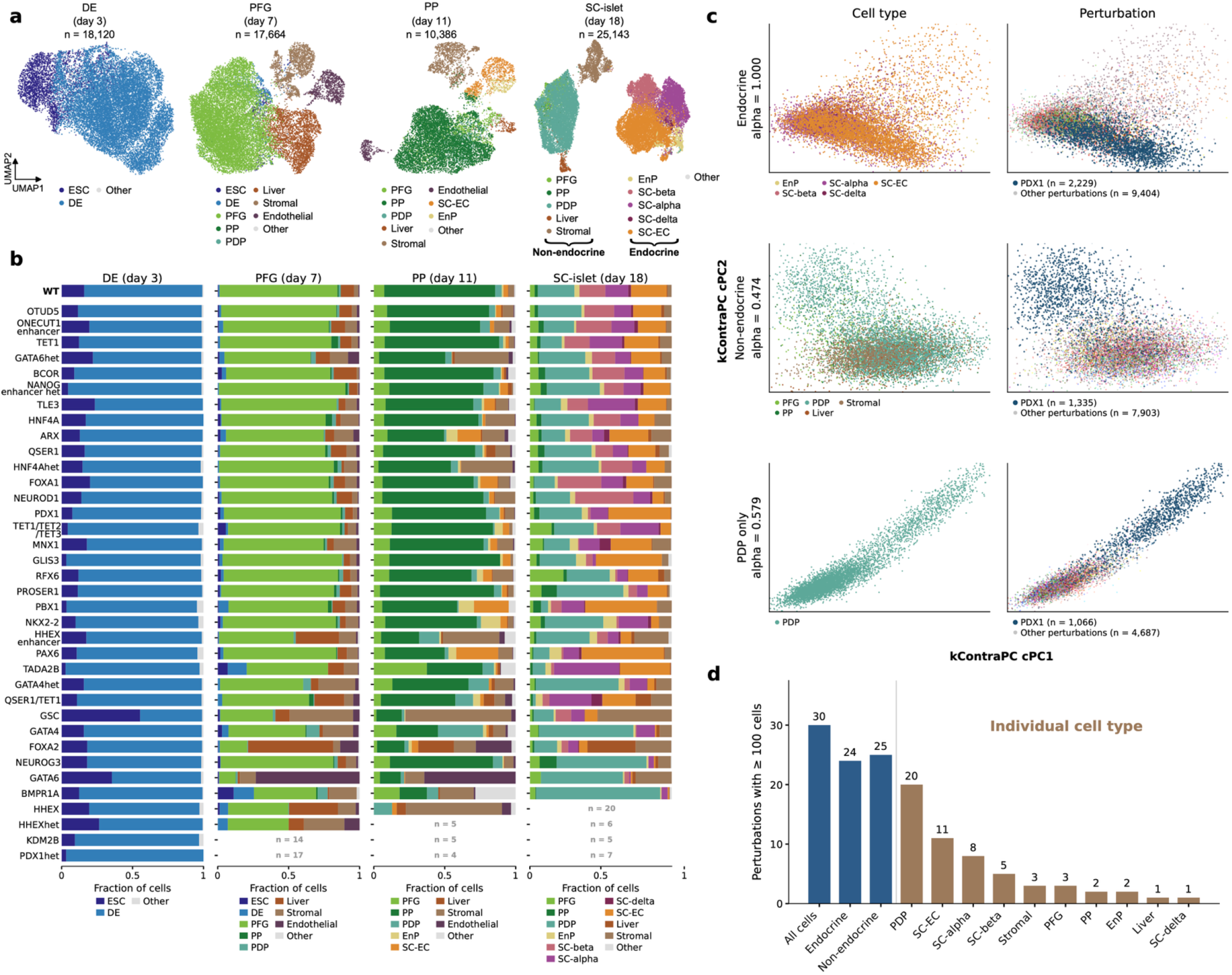
Perturbscape applied to a time-course stem-cell CRISPR-knockout village Perturb-seq dataset. (a) UMAP embeddings of cells at four differentiation stages, colored by cell type. Embeddings were computed independently per stage. Cell types with fewer than 100 cells at a stage are pooled as “Other”. n, cells per stage. (b) Cell type composition of each perturbation at each stage. Perturbation–stage combinations with fewer than 25 cells are blank and annotated with the cell count. (c) Kernel ContraPC (kContraPC) embeddings of day-18 cells, colored by cell type (left) and perturbation (right). Rows show progressively restricted populations: endocrine cells, non-endocrine cells, and pancreatic–duodenal progenitor (PDP) cells alone. Contrastive strength α is indicated per row. PDX1-perturbed cells separate in all three, including within PDP cells alone where cell type is held constant. Axis limits span the 0.5th–99.5th percentile. (d) Number of perturbations meeting the Perturbscape minimum of 100 cells at day 18, for compartments (blue) and individual cell types (brown), out of 36 perturbed genotypes (WT excluded). Source data are provided as a Source Data file.

**Supplementary Figure 10:**
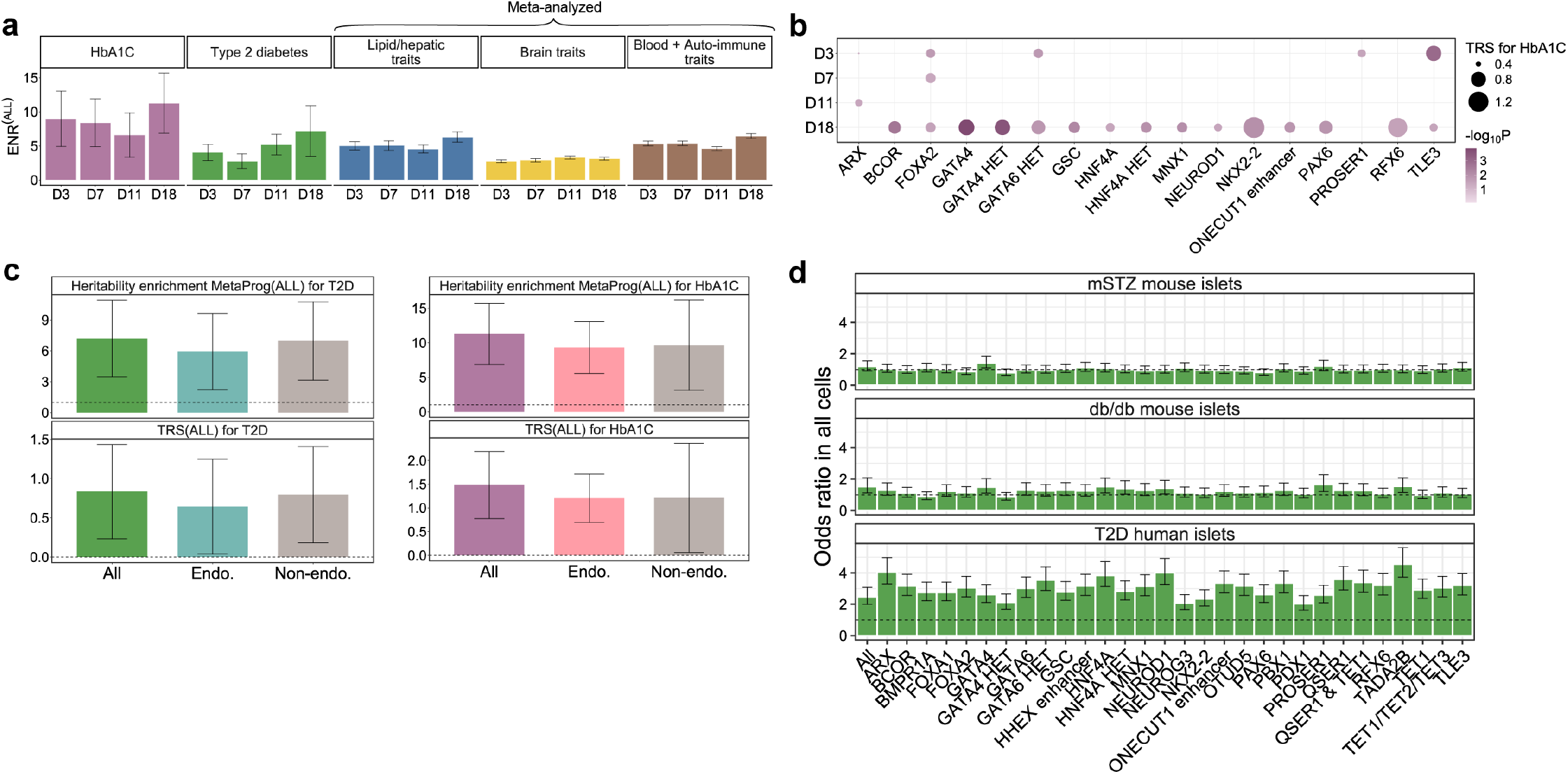
Additional analyses of pancreatic-differentiation perturbation programs. (a) Bar plots showing the heritability enrichment of MetaProg^(ALL)^ across the four pancreatic differentiation stages (Day 3, Day 7, Day 11 and Day 18). Enrichment is shown separately for HbA1c and type 2 diabetes (T2D), and meta-analyzed across lipid/hepatic, brain-related, and blood/autoimmune trait categories. Error bars represent 95% confidence intervals. (b) Perturbation-specific TRS for HbA1c across pancreatic differentiation. Magnitude (TRS, dot size) and significance (−log10P, dot color) of HbA1c heritability relevance of the meta-programs corresponding to perturbations at each of the four developmental stages (Days 3, 7, 11 and 18). Results are shown for the perturbations with nominally significant (P < 0.05) HbA1c TRS at one or more developmental stages. (c) Bar plots comparing MetaProg^(ALL)^ constructed from perturbation profiles of all cells (All), endocrine cells (Endo.) or non-endocrine cells (Non-endo.) at the SC-islet stage. Top, heritability enrichment of MetaProg^(ALL)^ for T2D (left) and HbA1c (right); bottom, corresponding trait relevance scores, TRS(ALL), for T2D (left) and HbA1c (right). Dashed lines denote an enrichment of 1 (top) and a TRS of 0 (bottom). Error bars represent 95% confidence intervals. (d) Enrichment of the top 500 genes from perturbation-specific T2D meta-programs in all cells among genes differentially expressed (Padj < 5%) in islet cells between (i) mSTZ-treated and wild-type (WT) mice, (ii) db/db and WT mice, and (iii) T2D and healthy donors from HPAP. Bars show enrichment odds ratios for individual diabetes-related gene perturbations; error bars represent 95% confidence intervals; the dashed horizontal line denotes an odds ratio of 1. Across perturbations, enrichment for HPAP genes was significantly stronger than enrichment for either db/db or streptozotocin-responsive genes (one-sided paired Wilcoxon P = 4.7e-10 for both db/db and streptozotocin-responsive genes), with mean differences in log odds ratio of 0.90 and 1.08 (95% bootstrap CI, 0.45–1.13 and 0.61–1.50). Source data are provided as a Source Data file.

**Supplementary Figure 11:**
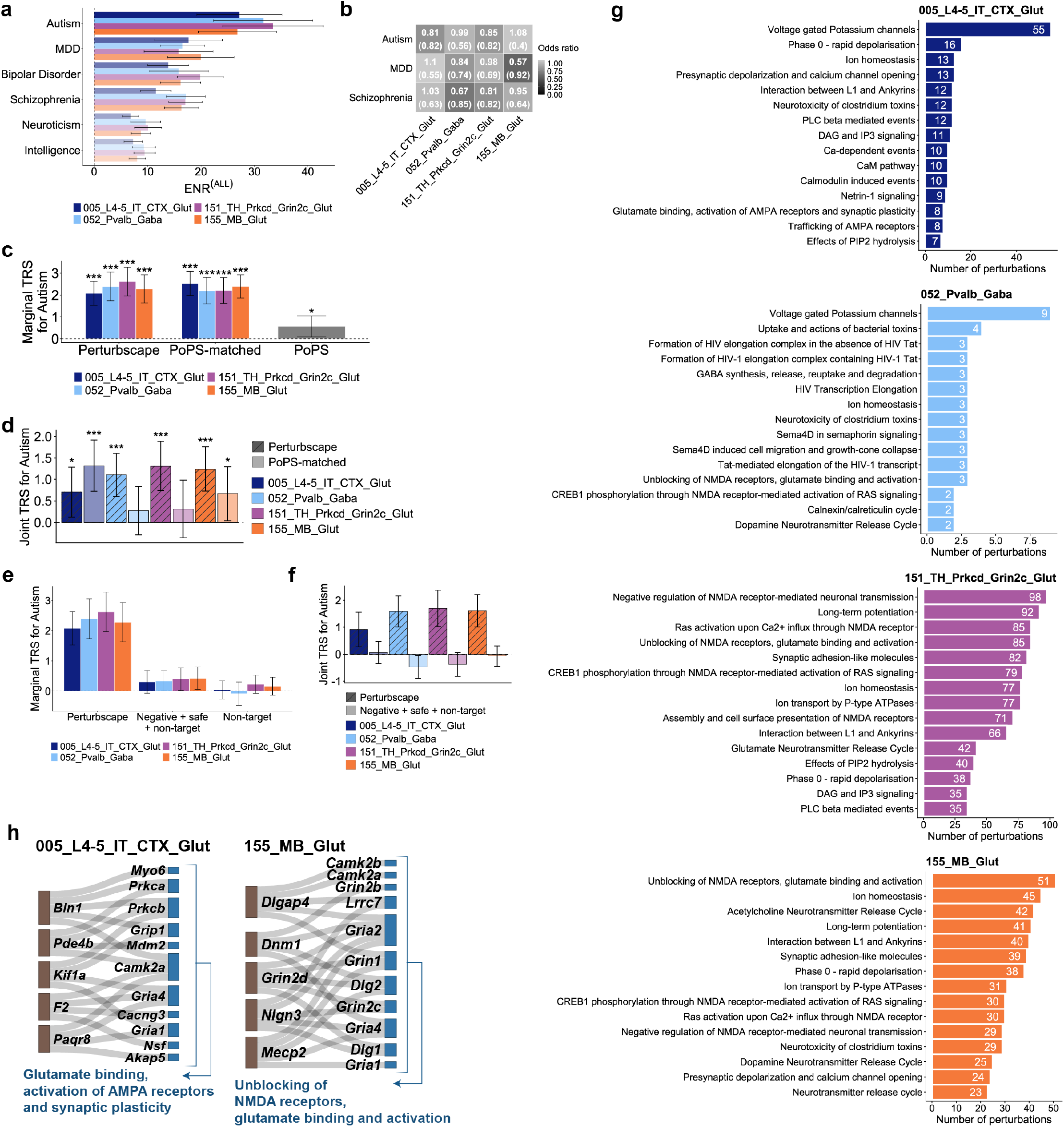
Additional analyses of autism-relevant perturbation programs across neuronal subclasses. (a) Bar plots showing the heritability enrichment of MetaProg^(ALL)^ for 6 brain-related traits across the 4 neuronal subclasses. Error bars represent 95% confidence intervals. (b) Heatmap showing, for each of four psychiatric traits (autism, bipolar disorder, MDD and schizophrenia) and each of the four neuronal subclasses, the enrichment odds ratio of overlap between the subclass-specific top 500 MetaProg^(ALL)^ genes and the panel genes selected in PerturbAI for each trait, with the corresponding p-value shown in parentheses. (c) Bar plots showing the marginal TRS for autism, comparing Perturbscape MetaProg(ALL), PoPS trained on the matched gene universe (“PoPS-matched”) and published PoPS, across the four neuronal subclasses. Asterisks denote statistical significance; error bars represent 95% confidence intervals. (d) Bar plots showing the joint TRS for autism, comparing Perturbscape MetaProg^(ALL)^ with PoPS-matched, across the four neuronal subclasses. Asterisks denote statistical significance; error bars represent 95% confidence intervals. (e) Bar plots showing marginal TRS for autism across the four neuronal subclasses, comparing Perturbscape MetaProg^(ALL)^ with meta-programs combining cNMF- and PCA-derived programs from negative, safe-target, and non-target guide cells (denoted “Negative + safe + non-target”) or only non-target guide cells (denoted “Non-target”). Error bars represent 95% confidence intervals. (f) Bar plots showing joint TRS for autism across the four neuronal subclasses, comparing Perturbscape MetaProg^(ALL)^ with meta-programs combining cNMF- and PCA-derived programs from negative, safe-target, and non-target guide cells (denoted “Negative + safe + non-target”) or only non-target guide cells (denoted “Non-target”). Error bars represent 95% confidence intervals. (g) Top enriched pathways for each of the four neuronal subclasses. Bars show the number of perturbations with significant TRS (p<0.05) whose top 500 meta-program genes are significantly enriched for each pathway; pathways are ordered by the number of perturbations. (h) Sankey diagrams showing pathway-level convergence of autism-relevant perturbations for the remaining two neuronal subclasses. (Left) In L4/5 IT cortical glutamatergic neurons (005_L4-5_IT_CTX_Glut), perturbations with significant autism TRS whose top 500 meta-program genes were significantly enriched for the “glutamate binding, activation of AMPA receptors and synaptic plasticity” pathway are linked to the prioritized genes belonging to this pathway. (Right) In midbrain glutamatergic neurons (155_MB_Glut), perturbations similarly enriched for the “unblocking of NMDA receptors, glutamate binding and activation” pathway are linked to the prioritized genes belonging to this pathway. Source data are provided as a Source Data file.

**Supplementary Figure 12:**
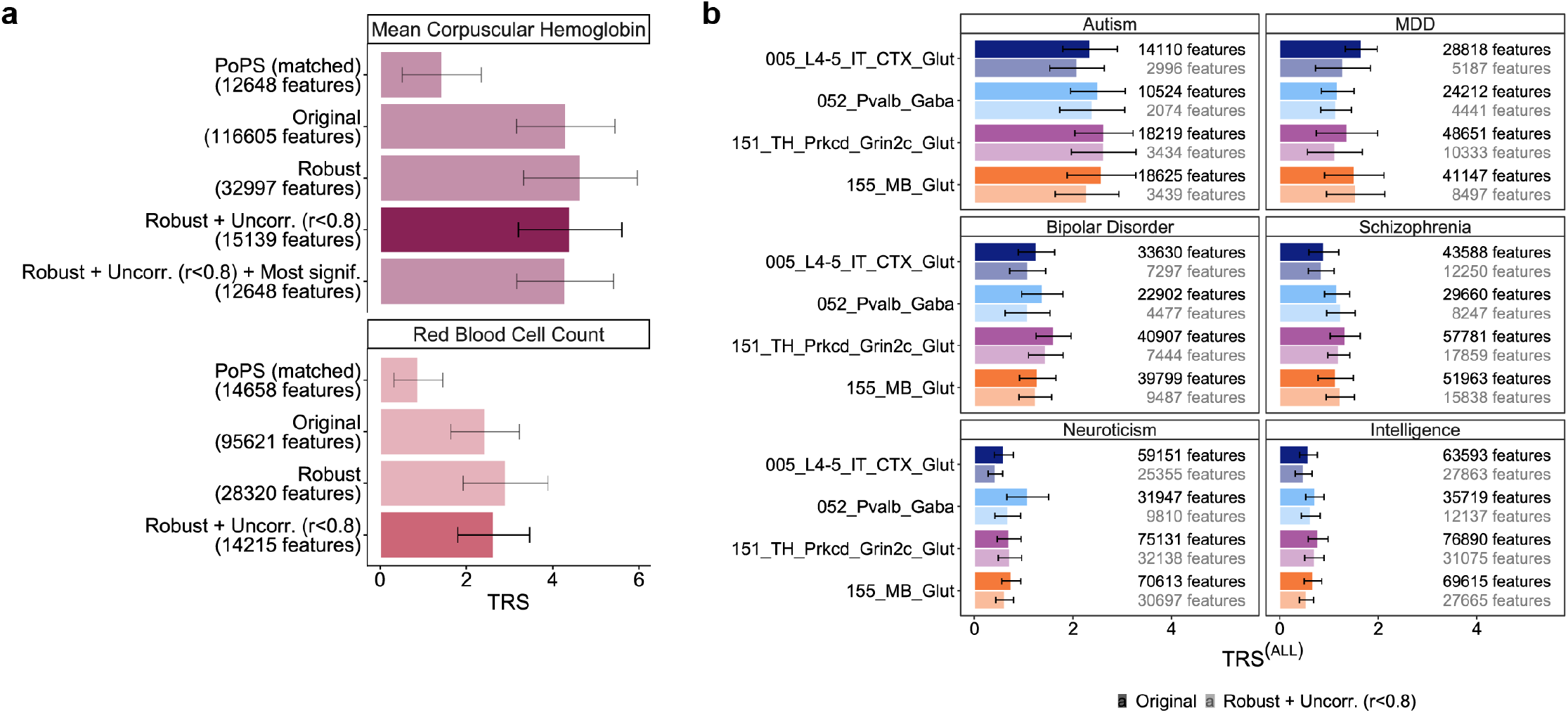
The atlas-wide meta-program is robust to restricting the program library to relatively uncorrelated features. (a) Bar plots showing the trait relevance score (TRS) of MetaProg^(ALL)^ for the two K562 erythroid traits— mean corpuscular hemoglobin (top) and red blood cell count (bottom)—when the meta-program is constructed under different feature-selection strategies applied to the perturbation-program library: the full library (“Original”); a reproducibility-filtered subset (“Robust”); the robust subset further restricted to relatively uncorrelated features (“Robust + Uncorr., r < 0.8”); and, for mean corpuscular hemoglobin, that subset additionally restricted to the most significant features (“Robust + Uncorr. (r < 0.8) + Most signif.”). TRS of PoPS trained on the matched gene universe (“PoPS (matched)”) is also compared. The number of features (programs) retained by each strategy is indicated in parentheses. Error bars represent 95% confidence intervals. The highlighted bars correspond to the MetaProg^(ALL)^ used as the default in all other analyses. (b) Bar plots showing the trait relevance scores (TRS) of MetaProg^(ALL)^ for 6 brain-related traits (autism, MDD, bipolar disorder, schizophrenia, neuroticism and intelligence) across the four PerturbAI neuronal subclasses, comparing the meta-program built from the full feature library (“Original”, darker bars) with that built from the relatively uncorrelated feature library (“Robust + Uncorr., r < 0.8”, lighter bars). The number of features retained under each strategy is indicated next to each bar. Error bars represent 95% confidence intervals. The uncorrelated-feature meta-program was adopted as the default for this high-perturbation-count screen (> 1,000 perturbations). Source data are provided as a Source Data file.

