## Supplementary Figure 1 for "Perturbation-induced gene programs map the hidden regulatory architecture of complex diseases"

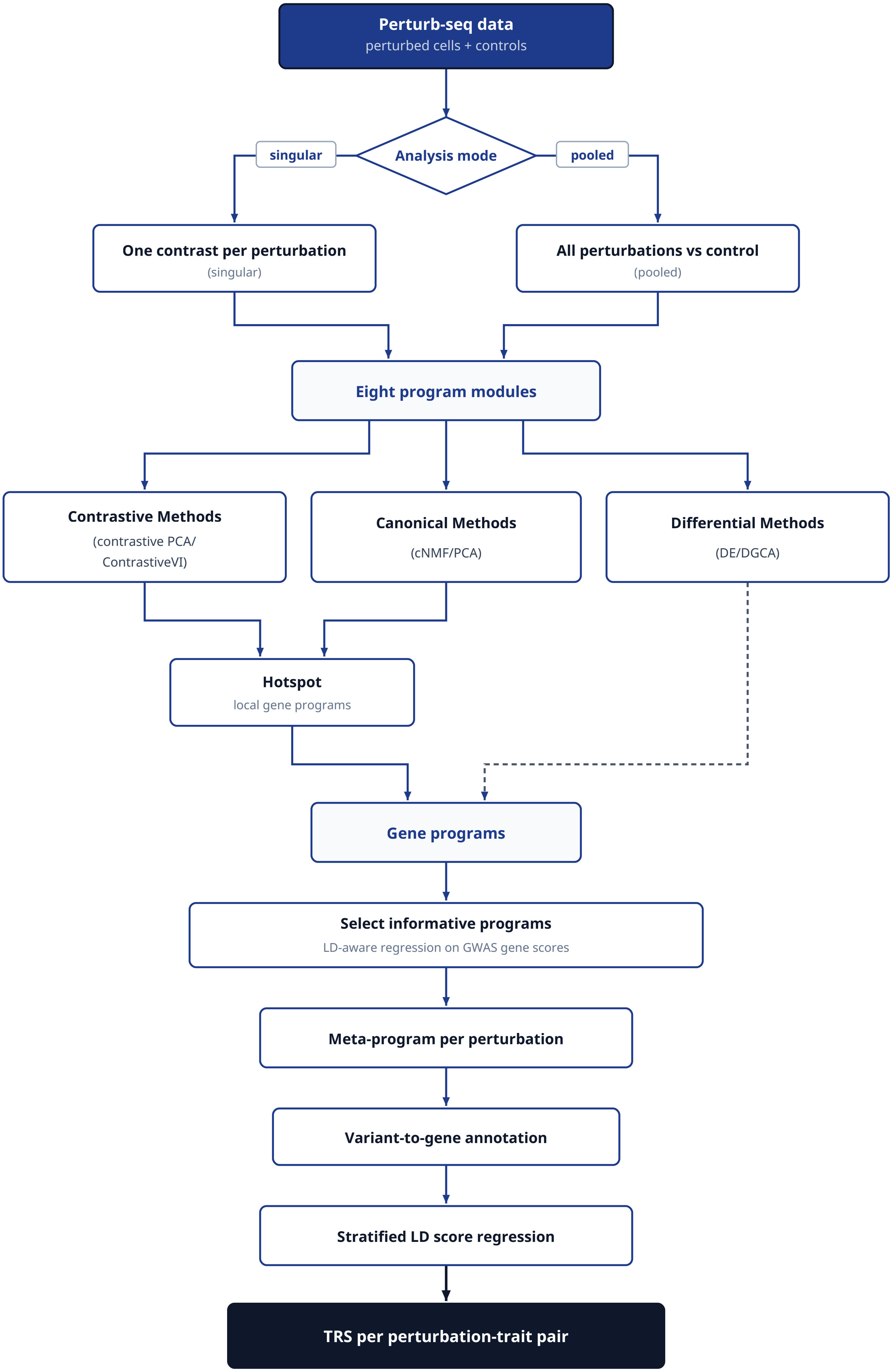

Perturb-seq data
perturbed cells + controls
Analysis mode
singular
pooled
One contrast per perturbation
(singular)
All perturbations vs control
(pooled)
Eight program modules
Contrastive Methods
(contrastive PCA/
ContrastiveVI)
Canonical Methods
(cNMF/PCA)
Differential Methods
(DE/DGCA)
Hotspot
local gene programs
Gene programs
Select informative programs
LD-aware regression on GWAS gene scores
Meta-program per perturbation
Variant-to-gene annotation
Stratified LD score regression
TRS per perturbation-trait pair
