## Supplementary Figure 2 for "Perturbation-induced gene programs map the hidden regulatory architecture of complex diseases"

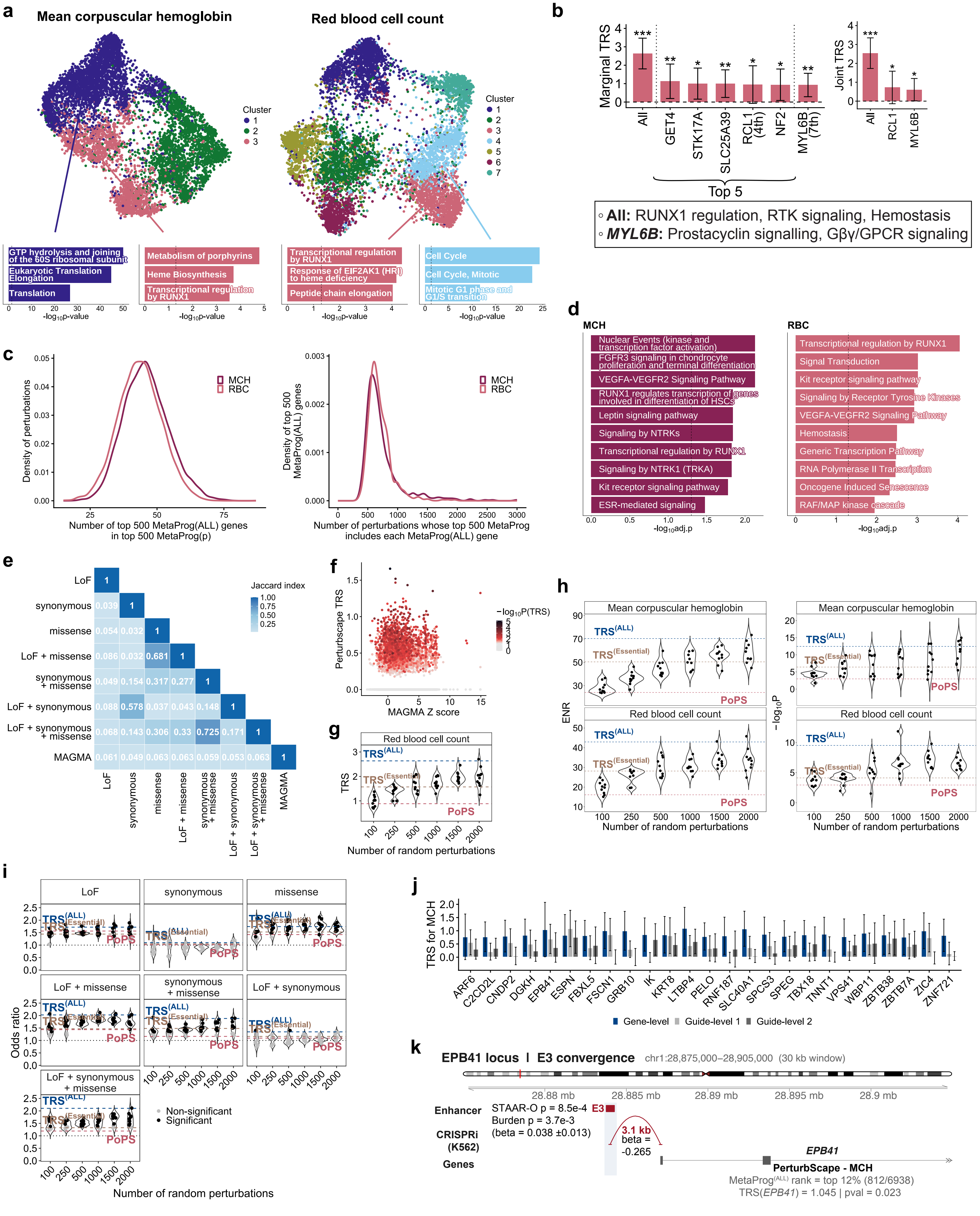
a
Mean corpuscular hemoglobin
Red blood cell count
Cluster
GTP hydrolysis and joining of the 60S ribosomal subunit
Eukaryotic Translation Elongation
Translation
Metabolism of porphyrins
Heme Biosynthesis
Transcriptional regulation by RUNX1
Transcriptional regulation by RUNX1
Response of EIF2AK1 (HRI) to heme deficiency
Peptide chain elongation
Cell Cycle
Cell Cycle, Mitotic
Mitotic G1 phase and G1/S transition
-log10p-value
b
Marginal TRS
All
GET4
STK17A
SLC25A39
RCL1 (4th)
NF2
MYL6B (7th)
Top 5
Joint TRS
All
RCL1
MYL6B
All: RUNX1 regulation, RTK signaling, Hemostasis
MYL6B: Prostacyclin signalling, Gβγ/GPCR signaling
c
MCH
RBC
Density of perturbations
Number of top 500 MetaProg(ALL) genes in top 500 MetaProg(p)
Density of top 500 MetaProg(ALL) genes
Number of perturbations whose top 500 MetaProg includes each MetaProg(ALL) gene
d
MCH
Nuclear Events (kinase and transcription factor activation)
FGFR3 signaling in chondrocyte proliferation and terminal differentiation
VEGFA-VEGFR2 Signaling Pathway
RUNX1 regulates transcription of genes involved in differentiation of HSCs
Leptin signaling pathway
Signaling by NTRKs
Transcriptional regulation by RUNX1
Signaling by NTRK1 (TRKA)
Kit receptor signaling pathway
ESR-mediated signaling
-log10adj.p
RBC
Transcriptional regulation by RUNX1
Signal Transduction
Kit receptor signaling pathway
Signaling by Receptor Tyrosine Kinases
VEGFA-VEGFR2 Signaling Pathway
Hemostasis
Generic Transcription Pathway
RNA Polymerase II Transcription
Oncogene Induced Senescence
RAF/MAP kinase cascade
-log10adj.p
e
Jaccard index
LoF
synonymous
missense
LoF + missense
synonymous + missense
LoF + synonymous
LoF + synonymous + missense
MAGMA
f
Perturbscape TRS
MAGMA Z score
−log10P(TRS)
g
Red blood cell count
TRS
TRS(ALL)
TRS(Essential)
PoPS
Number of random perturbations
h
Mean corpuscular hemoglobin
Red blood cell count
ENR
−log10P
TRS(ALL)
TRS(Essential)
PoPS
Number of random perturbations
i
LoF
synonymous
missense
LoF + missense
synonymous + missense
LoF + synonymous
LoF + synonymous + missense
Odds ratio
TRS(ALL)
TRS(Essential)
PoPS
Non-significant
Significant
Number of random perturbations
j
TRS for MCH
ARF6
C2CD2L
CNDP2
DGKH
EPB41
ESPN
FBXL5
FSCN1
GRB10
IK
KRT8
LTBP4
PELO
RNF187
SLC40A1
SPCS3
SPEG
TBX18
TNNT1
VPS41
WBP11
ZBTB38
ZBTB7A
ZIC4
ZNF721
Gene-level
Guide-level 1
Guide-level 2
k
EPB41 locus | E3 convergence
chr1:28,875,000–28,905,000 (30 kb window)
28.88 mb
28.885 mb
28.89 mb
28.895 mb
28.9 mb
Enhancer
STAAR-O p = 8.5e-4
Burden p = 3.7e-3
(beta = 0.038 ±0.013)
E3
CRISPRi (K562)
3.1 kb
beta = -0.265
Genes
EPB41
PerturbScape - MCH
MetaProg(ALL) rank = top 12% (812/6938)
TRS(EPB41) = 1.045 | pval = 0.023
