## Supplementary Figure 3 for "Perturbation-induced gene programs map the hidden regulatory architecture of complex diseases"

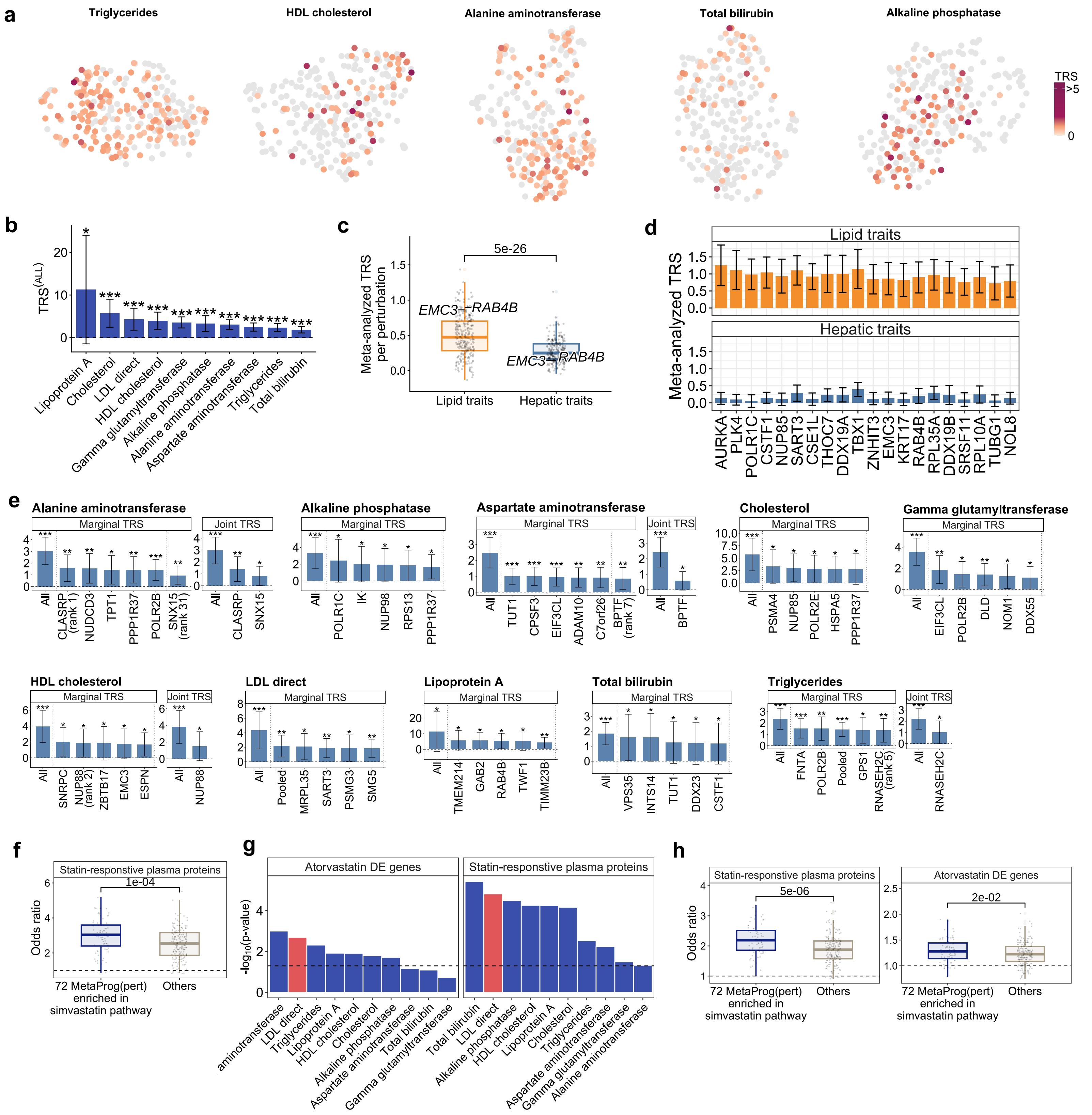
a
Triglycerides
HDL cholesterol
Alanine aminotransferase
Total bilirubin
Alkaline phosphatase
TRS
>5
0
b
TRS(ALL)
Lipoprotein A
Cholesterol
LDL direct
HDL cholesterol
Gamma glutamyltransferase
Alkaline phosphatase
Alanine aminotransferase
Aspartate aminotransferase
Triglycerides
Total bilirubin
c
Meta-analyzed TRS per perturbation
5e-26
EMC3—RAB4B
Lipid traits
Hepatic traits
d
Meta-analyzed TRS
Lipid traits
Hepatic traits
AURKA
PLK4
POLR1C
CSTF1
NUP85
SART3
CSE1L
THOC7
DDX19A
TBX1
ZNHIT3
EMC3
KRT17
RAB4B
RPL35A
DDX19B
SRSF11
RPL10A
TUBG1
NOL8
e
Alanine aminotransferase
Marginal TRS
Joint TRS
All
CLASRP (rank 1)
NUDCD3
TPT1
PPP1R37
POLR2B
SNX15 (rank 31)
Alkaline phosphatase
POLR1C
IK
NUP98
RPS13
Aspartate aminotransferase
TUT1
CPSF3
EIF3CL
ADAM10
C7orf26
BPTF (rank 7)
BPTF
Cholesterol
PSMA4
NUP85
POLR2E
HSPA5
Gamma glutamyltransferase
EIF3CL
DLD
NOM1
DDX55
HDL cholesterol
SNRPC
NUP88 (rank 2)
ZBTB17
EMC3
ESPN
NUP88
LDL direct
Pooled
MRPL35
SART3
PSMG3
SMG5
Lipoprotein A
TMEM214
GAB2
RAB4B
TWF1
TIMM23B
Total bilirubin
VPS35
INTS14
TUT1
DDX23
CSTF1
Triglycerides
FNTA
GPS1
RNASEH2C (rank 5)
RNASEH2C
f
Statin-responsive plasma proteins
1e-04
Odds ratio
72 MetaProg(pert) enriched in simvastatin pathway
Others
g
Atorvastatin DE genes
Statin-responsive plasma proteins
-log10(p-value)
h
5e-06
2e-02
