## Supplementary Figure 4 for "Perturbation-induced gene programs map the hidden regulatory architecture of complex diseases"

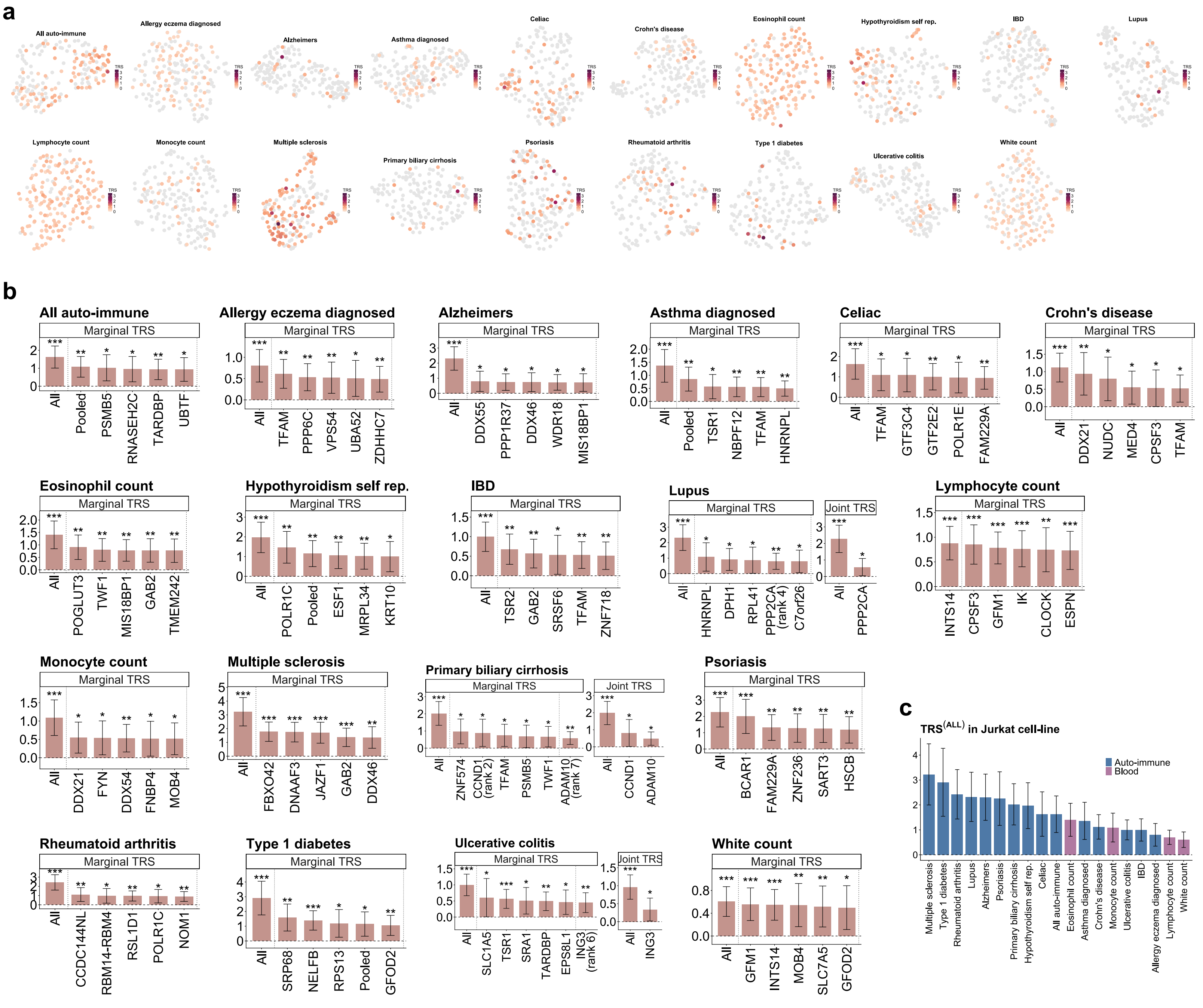
a
All auto-immune
Allergy eczema diagnosed
Alzheimers
Asthma diagnosed
Celiac
Crohn's disease
Eosinophil count
Hypothyroidism self rep.
IBD
Lupus
Lymphocyte count
Monocyte count
Multiple sclerosis
Primary biliary cirrhosis
Psoriasis
Rheumatoid arthritis
Type 1 diabetes
Ulcerative colitis
White count
b
Marginal TRS
Joint TRS
All
Pooled
PSMB5
RNASEH2C
TARDBP
UBTF
TFAM
PPP6C
VPS54
UBA52
ZDHHC7
DDX55
PPP1R37
DDX46
WDR18
MIS18BP1
TSR1
NBPF12
HNRNPL
GTF3C4
GTF2E2
POLR1E
FAM229A
DDX21
NUDC
MED4
CPSF3
POGLUT3
TWF1
GAB2
TMEM242
POLR1C
ESF1
MRPL34
KRT10
TSR2
SRSF6
ZNF718
DPH1
RPL41
PPP2CA (rank 4)
C7orf26
PPP2CA
INTS14
GFM1
IK
CLOCK
ESPN
FYN
DDX54
FNBP4
MOB4
FBXO42
DNAAF3
JAZF1
ZNF574
CCND1 (rank 2)
ADAM10 (rank 7)
CCND1
ADAM10
BCAR1
ZNF236
SART3
HSCB
CCDC144NL
RBM14-RBM4
RSL1D1
NOM1
SRP68
NELFB
RPS13
GFOD2
SLC1A5
SRA1
EPS8L1
ING3 (rank 6)
ING3
INTS14
SLC7A5
c
TRS(ALL) in Jurkat cell-line
Auto-immune
Blood
