## Supplementary Figure 5 for "Perturbation-induced gene programs map the hidden regulatory architecture of complex diseases"

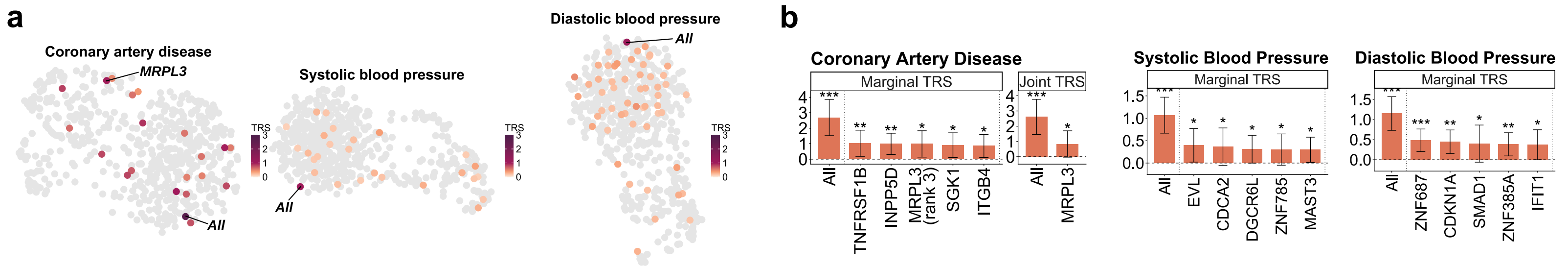
a
Coronary artery disease
MRPL3
All
TRS
3
2
1
0
Systolic blood pressure
All
TRS
3
2
1
0
Diastolic blood pressure
All
TRS
3
2
1
0
b
Coronary Artery Disease
Marginal TRS
Joint TRS
All
TNFRSF1B
INPP5D
MRPL3 (rank 3)
SGK1
ITGB4
All
MRPL3
Systolic Blood Pressure
Marginal TRS
All
EVL
CDCA2
DGCR6L
ZNF785
MAST3
Diastolic Blood Pressure
Marginal TRS
All
ZNF687
CDKN1A
SMAD1
ZNF385A
IFIT1
