## Supplementary Figure 6 for "Perturbation-induced gene programs map the hidden regulatory architecture of complex diseases"

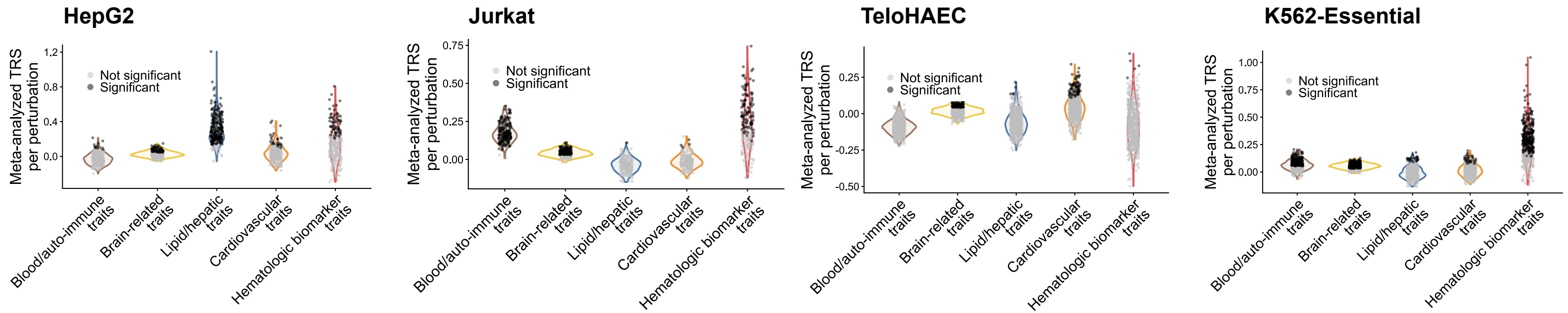
HepG2
Jurkat
TeloHAEC
K562-Essential
Meta-analyzed TRS per perturbation
Not significant
Significant
1.2
0.8
0.4
0.0
0.75
0.50
0.25
0.00
-0.25
-0.50
1.00
Blood/auto-immune traits
Brain-related traits
Lipid/hepatic traits
Cardiovascular traits
Hematologic biomarker traits
