## Supplementary Figure 7 for "Perturbation-induced gene programs map the hidden regulatory architecture of complex diseases"

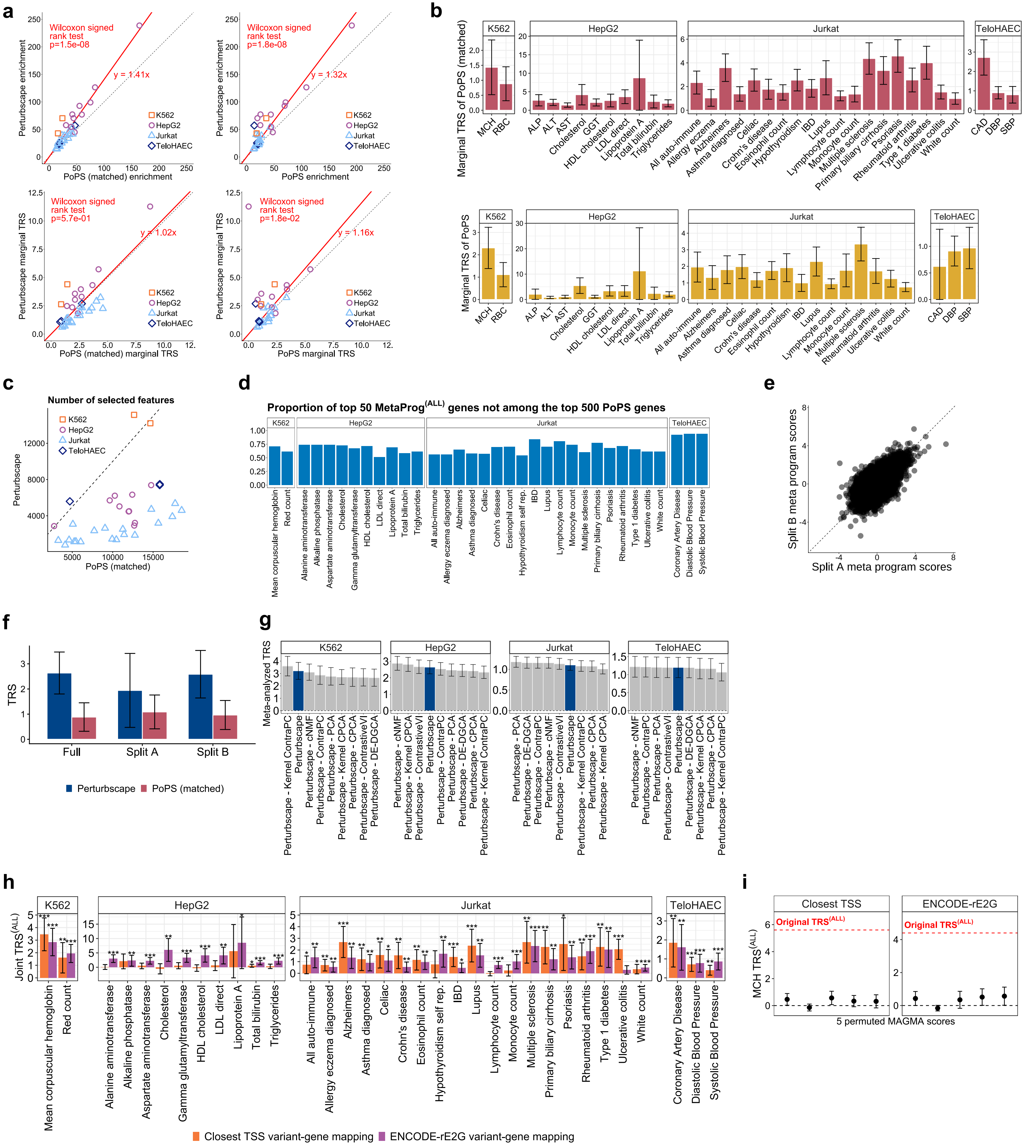

a
Wilcoxon signed rank test p=1.5e-08
y = 1.41x
Perturbscape enrichment
PoPS (matched) enrichment
Wilcoxon signed rank test p=1.8e-08
y = 1.32x
PoPS enrichment
Wilcoxon signed rank test p=5.7e-01
y = 1.02x
Perturbscape marginal TRS
PoPS (matched) marginal TRS
Wilcoxon signed rank test p=1.8e-02
y = 1.16x
PoPS marginal TRS
K562
HepG2
Jurkat
TeloHAEC
b
Marginal TRS of PoPS (matched)
Marginal TRS of PoPS
MCH
RBC
ALP
ALT
AST
Cholesterol
GGT
HDL cholesterol
LDL direct
Lipoprotein A
Total bilirubin
Triglycerides
All auto-immune
Allergy eczema
Alzheimers
Asthma diagnosed
Celiac
Crohn's disease
Eosinophil count
Hypothyroidism
IBD
Lupus
Lymphocyte count
Monocyte count
Multiple sclerosis
Primary biliary cirrhosis
Psoriasis
Rheumatoid arthritis
Type 1 diabetes
Ulcerative colitis
White count
CAD
DBP
SBP
c
Number of selected features
Perturbscape
PoPS (matched)
d
Proportion of top 50 MetaProg(ALL) genes not among the top 500 PoPS genes
Mean corpuscular hemoglobin
Red count
Alanine aminotransferase
Alkaline phosphatase
Aspartate aminotransferase
Gamma glutamyltransferase
Hypothyroidism self rep.
Coronary Artery Disease
Diastolic Blood Pressure
Systolic Blood Pressure
e
Split B meta program scores
Split A meta program scores
f
TRS
Full
Split A
Split B
Perturbscape
PoPS (matched)
g
Meta-analyzed TRS
Perturbscape - Kernel ContraPC
Perturbscape
Perturbscape - cNMF
Perturbscape - ContraPC
Perturbscape - PCA
Perturbscape - Kernel CPCA
Perturbscape - CPCA
Perturbscape - ContrastiveVI
Perturbscape - DE-DGCA
h
Joint TRS(ALL)
Closest TSS variant-gene mapping
ENCODE-rE2G variant-gene mapping
i
MCH TRS(ALL)
Closest TSS
ENCODE-rE2G
Original TRS(ALL)
5 permuted MAGMA scores
