## Supplementary Figure 8 for "Perturbation-induced gene programs map the hidden regulatory architecture of complex diseases"

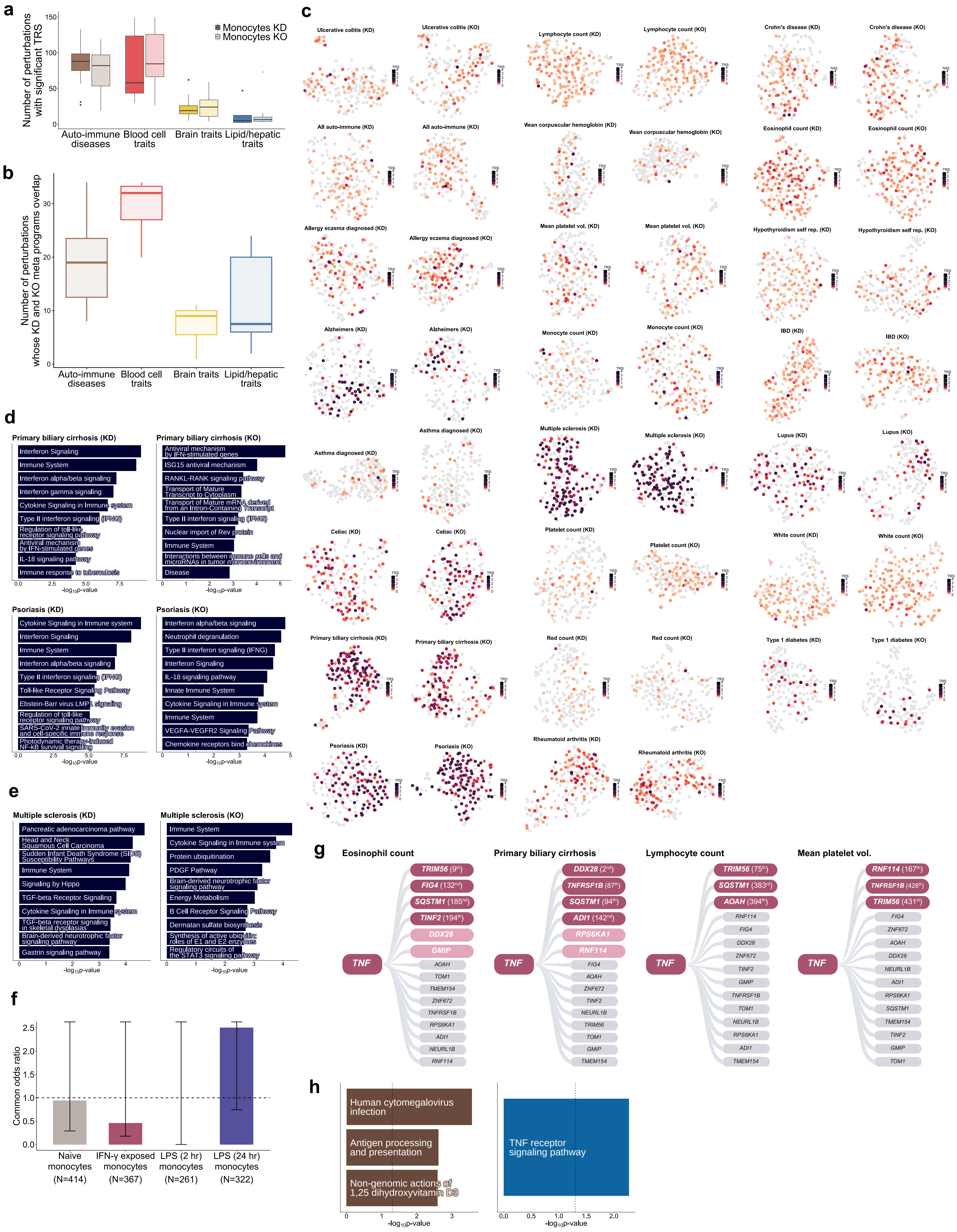

a
Number of perturbations with significant TRS
Monocytes KD
Monocytes KO
Auto-immune diseases
Blood cell traits
Brain traits
Lipid/hepatic traits
b
Number of perturbations whose KD and KO meta programs overlap
Auto-immune diseases
Blood cell traits
Brain traits
Lipid/hepatic traits
c
Ulcerative colitis (KD)
Ulcerative colitis (KO)
Lymphocyte count (KD)
Lymphocyte count (KO)
Crohn's disease (KD)
Crohn's disease (KO)
All auto-immune (KD)
All auto-immune (KO)
Mean corpuscular hemoglobin (KD)
Mean corpuscular hemoglobin (KO)
Eosinophil count (KD)
Eosinophil count (KO)
Allergy eczema diagnosed (KD)
Allergy eczema diagnosed (KO)
Mean platelet vol. (KD)
Mean platelet vol. (KO)
Hypothyroidism self rep. (KD)
Hypothyroidism self rep. (KO)
Alzheimers (KD)
Alzheimers (KO)
Monocyte count (KD)
Monocyte count (KO)
IBD (KD)
IBD (KO)
Asthma diagnosed (KD)
Asthma diagnosed (KO)
Multiple sclerosis (KD)
Multiple sclerosis (KO)
Lupus (KD)
Lupus (KO)
Celiac (KD)
Celiac (KO)
Platelet count (KD)
Platelet count (KO)
White count (KD)
White count (KO)
Primary biliary cirrhosis (KD)
Primary biliary cirrhosis (KO)
Red count (KD)
Red count (KO)
Type 1 diabetes (KD)
Type 1 diabetes (KO)
Psoriasis (KD)
Psoriasis (KO)
Rheumatoid arthritis (KD)
Rheumatoid arthritis (KO)
TRS
d
Primary biliary cirrhosis (KD)
Interferon Signaling
Immune System
Interferon alpha/beta signaling
Interferon gamma signaling
Cytokine Signaling in Immune system
Type II interferon signaling (IFNG)
Regulation of toll-like receptor signaling pathway
Antiviral mechanism by IFN-stimulated genes
IL-18 signaling pathway
Immune response to tuberculosis
$-\log_{10}$p-value
Primary biliary cirrhosis (KO)
Antiviral mechanism by IFN-stimulated genes
ISG15 antiviral mechanism
RANKL-RANK signaling pathway
Transport of Mature Transcript to Cytoplasm
Transport of Mature mRNA derived from an Intron-Containing Transcript
Type II interferon signaling (IFNG)
Nuclear import of Rev protein
Immune System
Interactions between immune cells and microRNAs in tumor microenvironment
Disease
Psoriasis (KD)
Cytokine Signaling in Immune system
Interferon Signaling
Immune System
Interferon alpha/beta signaling
Type II interferon signaling (IFNG)
Toll-like Receptor Signaling Pathway
Ebstein-Barr virus LMP1 signaling
Regulation of toll-like receptor signaling pathway
SARS-CoV-2 innate immunity evasion and cell-specific immune response
Photodynamic therapy-induced NF-kB survival signaling
Psoriasis (KO)
Interferon alpha/beta signaling
Neutrophil degranulation
Type II interferon signaling (IFNG)
Interferon Signaling
IL-18 signaling pathway
Innate Immune System
Cytokine Signaling in Immune system
Immune System
VEGFA-VEGFR2 Signaling Pathway
Chemokine receptors bind chemokines
e
Multiple sclerosis (KD)
Pancreatic adenocarcinoma pathway
Head and Neck Squamous Cell Carcinoma
Sudden Infant Death Syndrome (SIDS) Susceptibility Pathways
Immune System
Signaling by Hippo
TGF-beta Receptor Signaling
Cytokine Signaling in Immune system
TGF-beta receptor signaling in skeletal dysplasias
Brain-derived neurotrophic factor signaling pathway
Gastrin signaling pathway
Multiple sclerosis (KO)
Immune System
Cytokine Signaling in Immune system
Protein ubiquitination
PDGF Pathway
Brain-derived neurotrophic factor signaling pathway
Energy Metabolism
B Cell Receptor Signaling Pathway
Dermatan sulfate biosynthesis
Synthesis of active ubiquitin: roles of E1 and E2 enzymes
Regulatory circuits of the STAT3 signaling pathway
f
Common odds ratio
Naive monocytes (N=414)
IFN-γ exposed monocytes (N=367)
LPS (2 hr) monocytes (N=261)
LPS (24 hr) monocytes (N=322)
g
Eosinophil count
TNF
TRIM56 (9th)
FIG4 (132nd)
SQSTM1 (185nd)
TINF2 (194th)
DDX28
GMIP
AOAH
TOM1
TMEM154
ZNF672
TNFRSF1B
RPS6KA1
ADI1
NEURL1B
RNF114
Primary biliary cirrhosis
TNF
DDX28 (2nd)
TNFRSF1B (87th)
SQSTM1 (94th)
ADI1 (142nd)
RPS6KA1
RNF114
FIG4
AOAH
ZNF672
TINF2
NEURL1B
TRIM56
TOM1
GMIP
TMEM154
Lymphocyte count
TNF
TRIM56 (75th)
SQSTM1 (383rd)
AOAH (394th)
RNF114
FIG4
DDX28
ZNF672
TINF2
GMIP
TNFRSF1B
TOM1
NEURL1B
RPS6KA1
ADI1
TMEM154
Mean platelet vol.
TNF
RNF114 (167th)
TNFRSF1B (428th)
TRIM56 (431st)
FIG4
ZNF672
AOAH
DDX28
NEURL1B
ADI1
RPS6KA1
SQSTM1
TMEM154
TINF2
GMIP
TOM1
h
Human cytomegalovirus infection
Antigen processing and presentation
Non-genomic actions of 1,25 dihydroxyvitamin D3
TNF receptor signaling pathway
$-\log_{10}$p-value
