## Supplementary Figure 9 for "Perturbation-induced gene programs map the hidden regulatory architecture of complex diseases"

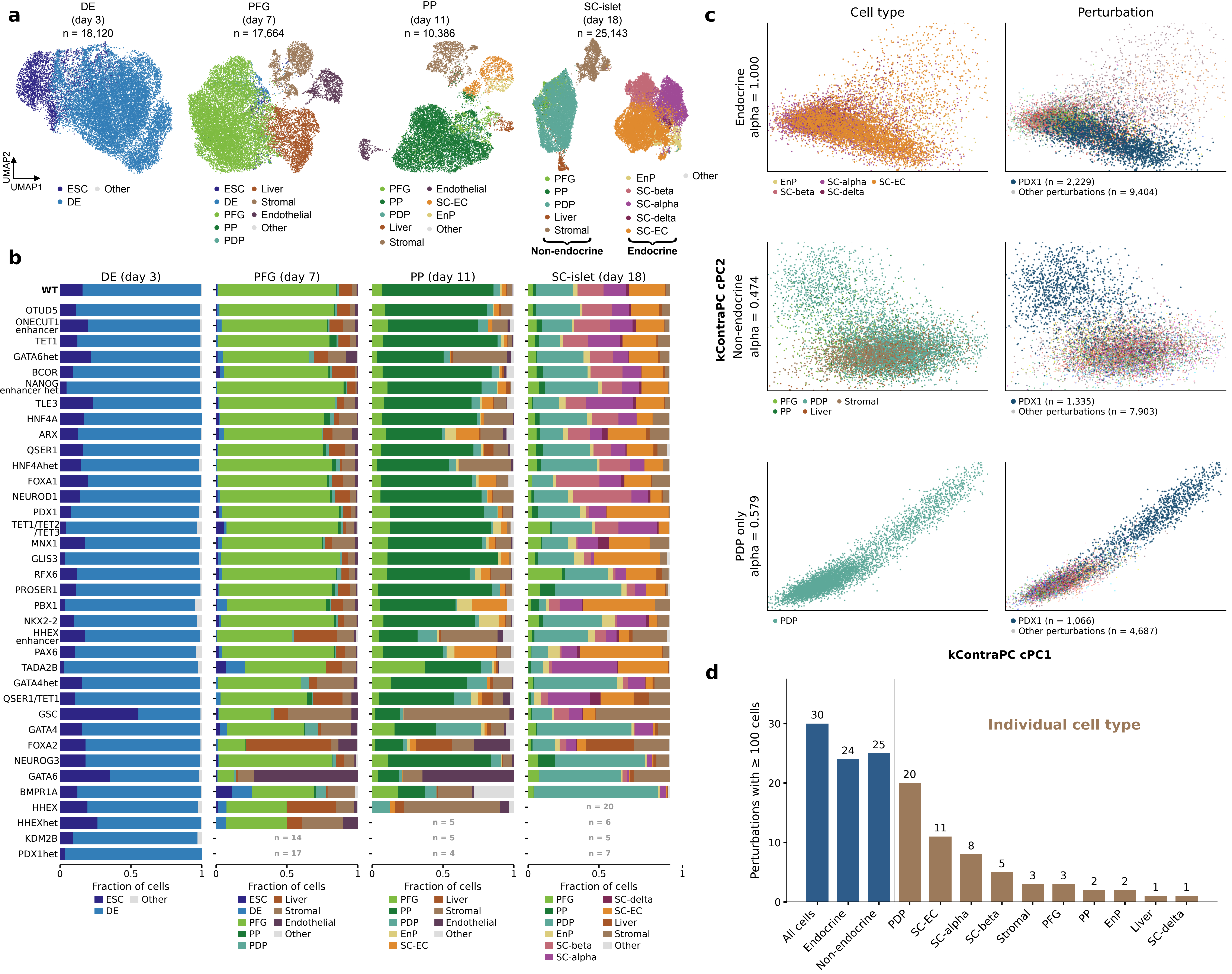

a
DE (day 3) n = 18,120
PFG (day 7) n = 17,664
PP (day 11) n = 10,386
SC-islet (day 18) n = 25,143
UMAP1
UMAP2
ESC
DE
Other
PFG
PP
PDP
Liver
Stromal
Endothelial
SC-EC
EnP
SC-beta
SC-alpha
SC-delta
Non-endocrine
Endocrine
b
DE (day 3)
PFG (day 7)
PP (day 11)
SC-islet (day 18)
WT
OTUD5
ONECUT1 enhancer
TET1
GATA6het
BCOR
NANOG enhancer het
TLE3
HNF4A
ARX
QSER1
HNF4Ahet
FOXA1
NEUROD1
PDX1
TET1/TET2 /TET3
MNX1
GLIS3
RFX6
PROSER1
PBX1
NKX2-2
HHEX enhancer
PAX6
TADA2B
GATA4het
QSER1/TET1
GSC
GATA4
FOXA2
NEUROG3
GATA6
BMPR1A
HHEX
HHEXhet
KDM2B
PDX1het
n = 20
n = 5
n = 6
n = 14
n = 5
n = 5
n = 17
n = 4
n = 7
0
0.5
1
Fraction of cells
c
Cell type
Perturbation
Endocrine alpha = 1.000
PDX1 (n = 2,229)
Other perturbations (n = 9,404)
kContraPC cPC2
Non-endocrine alpha = 0.474
PDX1 (n = 1,335)
Other perturbations (n = 7,903)
PDP only alpha = 0.579
PDX1 (n = 1,066)
Other perturbations (n = 4,687)
kContraPC cPC1
d
Perturbations with ≥ 100 cells
Individual cell type
30
24
25
20
11
8
5
3
3
2
2
1
1
0
10
20
30
All cells
Endocrine
Non-endocrine
PDP
SC-EC
SC-alpha
SC-beta
Stromal
PFG
PP
EnP
Liver
SC-delta
