## Supplementary Figure 10 for "Perturbation-induced gene programs map the hidden regulatory architecture of complex diseases"

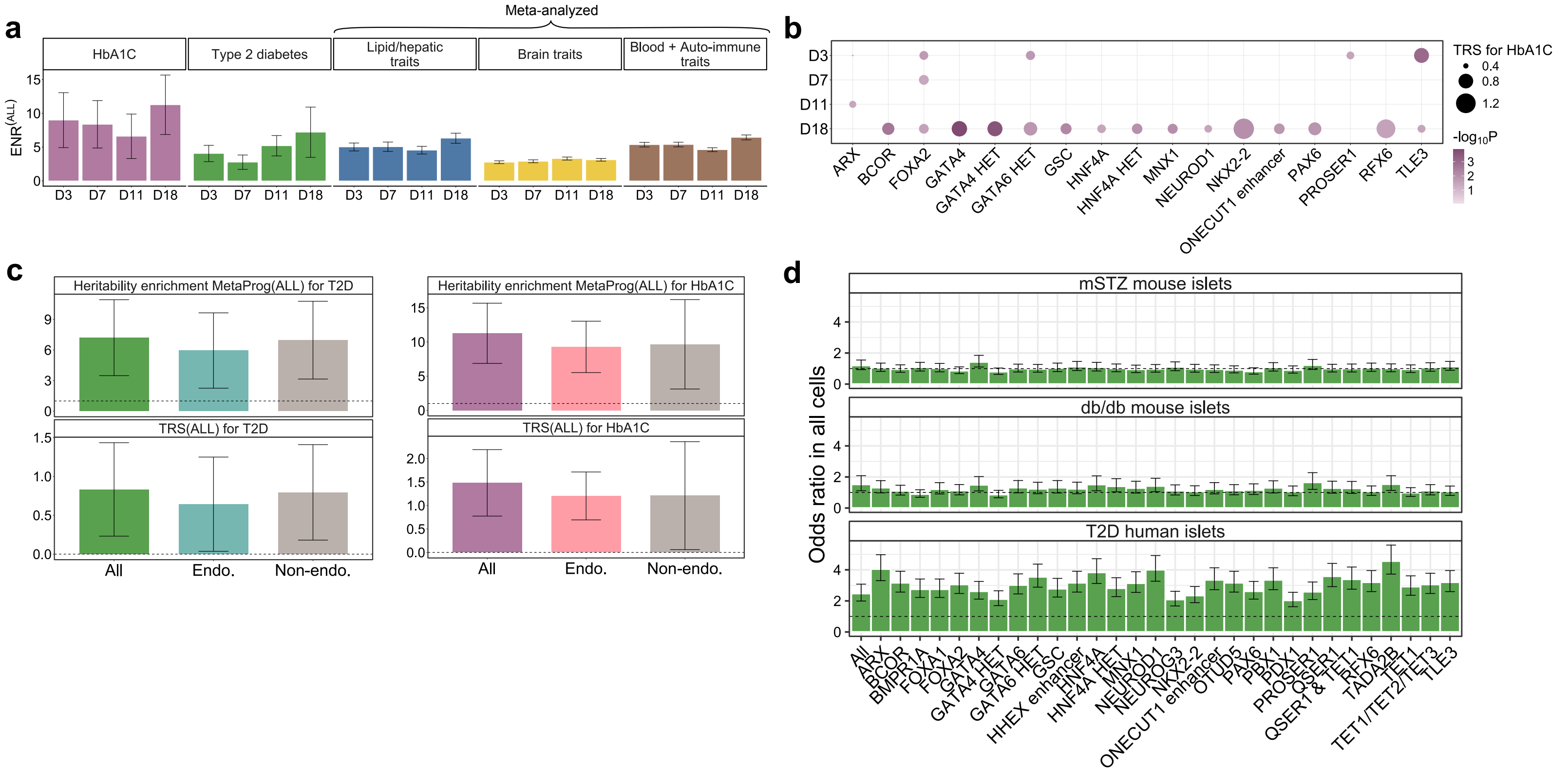
a
Meta-analyzed
HbA1C
Type 2 diabetes
Lipid/hepatic traits
Brain traits
Blood + Auto-immune traits
ENR(ALL)
D3
D7
D11
D18
b
TRS for HbA1C
-log10P
ARX
BCOR
FOXA2
GATA4
GATA4 HET
GATA6 HET
GSC
HNF4A
HNF4A HET
MNX1
NEUROD1
NKX2-2
ONECUT1 enhancer
PAX6
PROSER1
RFX6
TLE3
c
Heritability enrichment MetaProg(ALL) for T2D
Heritability enrichment MetaProg(ALL) for HbA1C
TRS(ALL) for T2D
TRS(ALL) for HbA1C
All
Endo.
Non-endo.
d
mSTZ mouse islets
db/db mouse islets
T2D human islets
Odds ratio in all cells
All
ARX
BCOR
BMPR1A
FOXA1
FOXA2
GATA4
GATA4 HET
GATA6
GATA6 HET
GSC
HHEX enhancer
HNF4A
HNF4A HET
MNX1
NEUROD1
NEUROG3
NKX2-2
ONECUT1 enhancer
OTUD5
PAX6
PBX1
PDX1
PROSER1
QSER1
QSER1 & TET1
RFX6
TADA2B
TET1
TET1/TET2/TET3
TLE3
