## Supplementary Figure 11 for "Perturbation-induced gene programs map the hidden regulatory architecture of complex diseases"

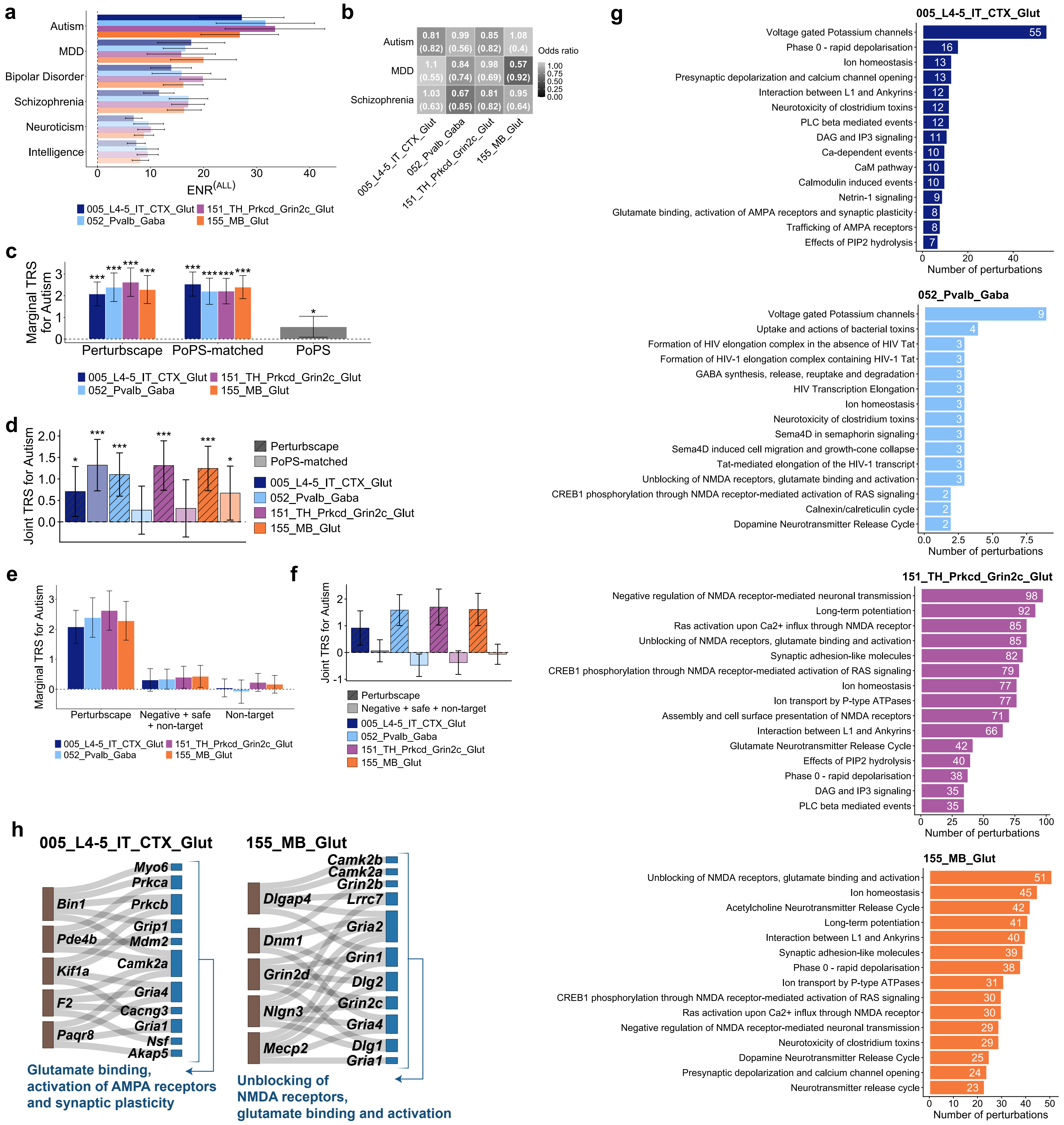
a
Autism
MDD
Bipolar Disorder
Schizophrenia
Neuroticism
Intelligence
0
10
20
30
40
ENR(ALL)
005_L4-5_IT_CTX_Glut
151_TH_Prkcd_Grin2c_Glut
052_Pvalb_Gaba
155_MB_Glut
b
Autism
0.81 (0.82)
0.99 (0.56)
0.85 (0.82)
1.08 (0.4)
MDD
1.1 (0.55)
0.84 (0.74)
0.98 (0.69)
0.57 (0.92)
Schizophrenia
1.03 (0.63)
0.67 (0.85)
0.81 (0.82)
0.95 (0.64)
Odds ratio
1.00
0.75
0.50
0.25
0.00
005_L4-5_IT_CTX_Glut
052_Pvalb_Gaba
151_TH_Prkcd_Grin2c_Glut
155_MB_Glut
c
Marginal TRS for Autism
Perturbscape
PoPS-matched
PoPS
005_L4-5_IT_CTX_Glut
151_TH_Prkcd_Grin2c_Glut
052_Pvalb_Gaba
155_MB_Glut
d
Joint TRS for Autism
2.0
1.5
1.0
0.5
0.0
Perturbscape
PoPS-matched
005_L4-5_IT_CTX_Glut
052_Pvalb_Gaba
151_TH_Prkcd_Grin2c_Glut
155_MB_Glut
e
Marginal TRS for Autism
Perturbscape
Negative + safe + non-target
Non-target
005_L4-5_IT_CTX_Glut
151_TH_Prkcd_Grin2c_Glut
052_Pvalb_Gaba
155_MB_Glut
f
Joint TRS for Autism
Perturbscape
Negative + safe + non-target
005_L4-5_IT_CTX_Glut
052_Pvalb_Gaba
151_TH_Prkcd_Grin2c_Glut
155_MB_Glut
g
005_L4-5_IT_CTX_Glut
Voltage gated Potassium channels 55
Phase 0 - rapid depolarisation 16
Ion homeostasis 13
Presynaptic depolarization and calcium channel opening 13
Interaction between L1 and Ankyrins 12
Neurotoxicity of clostridium toxins 12
PLC beta mediated events 12
DAG and IP3 signaling 11
Ca-dependent events 10
CaM pathway 10
Calmodulin induced events 10
Netrin-1 signaling 9
Glutamate binding, activation of AMPA receptors and synaptic plasticity 8
Trafficking of AMPA receptors 8
Effects of PIP2 hydrolysis 7
Number of perturbations
052_Pvalb_Gaba
Voltage gated Potassium channels 9
Uptake and actions of bacterial toxins 4
Formation of HIV elongation complex in the absence of HIV Tat 3
Formation of HIV-1 elongation complex containing HIV-1 Tat 3
GABA synthesis, release, reuptake and degradation 3
HIV Transcription Elongation 3
Ion homeostasis 3
Neurotoxicity of clostridium toxins 3
Sema4D in semaphorin signaling 3
Sema4D induced cell migration and growth-cone collapse 3
Tat-mediated elongation of the HIV-1 transcript 3
Unblocking of NMDA receptors, glutamate binding and activation 3
CREB1 phosphorylation through NMDA receptor-mediated activation of RAS signaling 2
Calnexin/calreticulin cycle 2
Dopamine Neurotransmitter Release Cycle 2
Number of perturbations
151_TH_Prkcd_Grin2c_Glut
Negative regulation of NMDA receptor-mediated neuronal transmission 98
Long-term potentiation 92
Ras activation upon Ca2+ influx through NMDA receptor 85
Unblocking of NMDA receptors, glutamate binding and activation 85
Synaptic adhesion-like molecules 82
CREB1 phosphorylation through NMDA receptor-mediated activation of RAS signaling 79
Ion homeostasis 77
Ion transport by P-type ATPases 77
Assembly and cell surface presentation of NMDA receptors 71
Interaction between L1 and Ankyrins 66
Glutamate Neurotransmitter Release Cycle 42
Effects of PIP2 hydrolysis 40
Phase 0 - rapid depolarisation 38
DAG and IP3 signaling 35
PLC beta mediated events 35
Number of perturbations
155_MB_Glut
Unblocking of NMDA receptors, glutamate binding and activation 51
Ion homeostasis 45
Acetylcholine Neurotransmitter Release Cycle 42
Long-term potentiation 41
Interaction between L1 and Ankyrins 40
Synaptic adhesion-like molecules 39
Phase 0 - rapid depolarisation 38
Ion transport by P-type ATPases 31
CREB1 phosphorylation through NMDA receptor-mediated activation of RAS signaling 30
Ras activation upon Ca2+ influx through NMDA receptor 30
Negative regulation of NMDA receptor-mediated neuronal transmission 29
Neurotoxicity of clostridium toxins 29
Dopamine Neurotransmitter Release Cycle 25
Presynaptic depolarization and calcium channel opening 24
Neurotransmitter release cycle 23
Number of perturbations
h
005_L4-5_IT_CTX_Glut
Bin1
Pde4b
Kif1a
F2
Paqr8
Myo6
Prkca
Prkcb
Grip1
Mdm2
Camk2a
Gria4
Cacng3
Gria1
Nsf
Akap5
Glutamate binding, activation of AMPA receptors and synaptic plasticity
155_MB_Glut
Dlgap4
Dnm1
Grin2d
Nlgn3
Mecp2
Camk2b
Camk2a
Grin2b
Lrrc7
Gria2
Grin1
Dlg2
Grin2c
Gria4
Dlg1
Gria1
Unblocking of NMDA receptors, glutamate binding and activation
