## Supplementary Figure 12 for "Perturbation-induced gene programs map the hidden regulatory architecture of complex diseases"

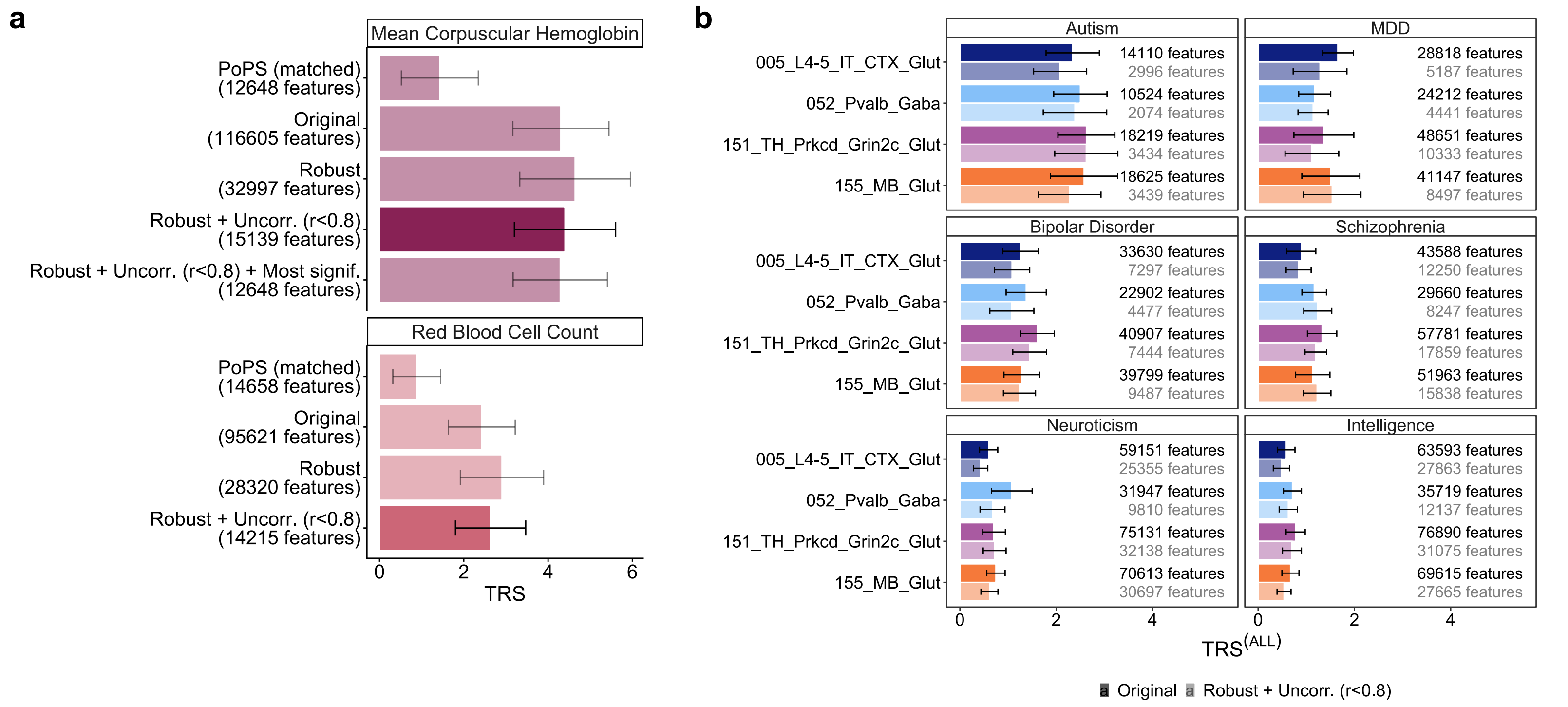
a
Mean Corpuscular Hemoglobin
PoPS (matched)
(12648 features)
Original
(116605 features)
Robust
(32997 features)
Robust + Uncorr. (r<0.8)
(15139 features)
Robust + Uncorr. (r<0.8) + Most signif.
(12648 features)
Red Blood Cell Count
PoPS (matched)
(14658 features)
Original
(95621 features)
Robust
(28320 features)
Robust + Uncorr. (r<0.8)
(14215 features)
0
2
4
6
TRS
b
Autism
005_L4-5_IT_CTX_Glut
052_Pvalb_Gaba
151_TH_Prkcd_Grin2c_Glut
155_MB_Glut
14110 features
2996 features
10524 features
2074 features
18219 features
3434 features
18625 features
3439 features
MDD
28818 features
5187 features
24212 features
4441 features
48651 features
10333 features
41147 features
8497 features
Bipolar Disorder
33630 features
7297 features
22902 features
4477 features
40907 features
7444 features
39799 features
9487 features
Schizophrenia
43588 features
12250 features
29660 features
8247 features
57781 features
17859 features
51963 features
15838 features
Neuroticism
59151 features
25355 features
31947 features
9810 features
75131 features
32138 features
70613 features
30697 features
Intelligence
63593 features
27863 features
35719 features
12137 features
76890 features
31075 features
69615 features
27665 features
0
2
4
TRS(ALL)
Original
Robust + Uncorr. (r<0.8)
