## Supplementary Note for "Perturbation-induced gene programs map the hidden regulatory architecture of complex diseases"

### Supplementary Notes

#### 1 Genome build and gene annotations

All coordinates are reported in GRCh38 unless otherwise specified. We used a curated set of 20,678 gene promoters (one per gene symbol), defined as 500-bp regions centered on the RefSeq TSS with the largest number of coding isoforms. Gene symbols were matched to ENSEMBL IDs using the HUGO database [1] and GENCODE v.29, retaining genes annotated as protein_coding, processed_transcript or lincRNA by GENCODE.

#### 2 Selection of kernel parameters

The inputs $\tilde{\boldsymbol{X}}_t$ and $\tilde{\boldsymbol{X}}_b$ were min–max scaled to the unit interval [0,1], satisfying the bounded-domain assumption required by random-feature approximations [2, 3] described in the next section. Under this scaling, the squared distance $\|\tilde{\boldsymbol{X}}_c(i,\cdot) - \tilde{\boldsymbol{X}}_c(j,\cdot)\|_2^2$ lies in $[0, N_c]$ (for $c \in \{t, b\}$) and the expected pairwise distance is of order $\sqrt{N_c}$; we therefore set $\sigma = \sqrt{N_c}$, keeping the kernel non-degenerate independently of dimension. The contrast parameter $\alpha$ was selected following the same procedure used for cPCA (for kCPCA) and for ContraPC (for kContraPC).

#### 3 Fastfood approximation for scalable kernel contrastive analysis

##### 3.1 Fastfood kernel approximation

The Gaussian RBF kernel, $k(x, x') = \exp(-\|x - x'\|^2/2\sigma^2)$, becomes expensive to use as the number of samples grows, since forming the kernel matrix over $G$ points with $N_c$ dimensions costs $O(G^2 N_c)$ time and $O(G^2)$ memory. A widely used remedy is applying Random Fourier Features [3], which draws a random Gaussian matrix $W \in \mathbb{R}^{m_c \times N_c}$ and map each point to a random-feature vector $\phi(x)$ whose inner products approximate the kernel, $\phi(x)^\top \phi(x') \approx k(x, x')$. This replaces the $G \times G$ kernel matrix with a $G \times m_c$ feature matrix; however, the dense multiplication $Wx$ still costs $O(m_c N_c)$ time and $O(m_c N_c)$ storage, which becomes the bottleneck in high dimensions. Fastfood [2] accelerates this step by replacing the dense Gaussian matrix with a product of structured matrices that behaves like a Gaussian matrix while supporting fast multiplication, reducing the cost to $O(m_c \log N_c)$ time and $O(m_c)$ memory. The resulting features provide an unbiased estimate of the kernel with concentration guarantees, making it practical to approximate the RBF kernel matrix when the input dimension $N_c$ is large. In our analysis, we use the `kernel_approximation.Fastfood` function from the Python package `scikit-learn-extra` (v0.2.0). To ensure convergence, we set $m_c$ to be $2 \cdot N_c$.

##### 3.2 Setup

The Fastfood approximation maps each (scaled) gene-expression vector to a random-feature vector, yielding two feature matrices

$$M_t \in \mathbb{R}^{m_t \times G}, \qquad M_b \in \mathbb{R}^{m_b \times G},$$

where $G$ is the number of genes and $m_t, m_b$ are the numbers of random features for the target and background groups. The Gaussian RBF kernel matrices $K_t$ and $K_b$ are approximated by $\tilde{K}_t$ and $\tilde{K}_b$ which are defined as follows:

$$\tilde{K}_t \;:=\; M_t^\top M_t \in \mathbb{R}^{G \times G}, \qquad \tilde{K}_b \;:=\; M_b^\top M_b \in \mathbb{R}^{G \times G}.$$

Both are symmetric positive semidefinite and are low rank when $m_t, m_b < G$, since $\operatorname{rank}(\tilde{K}_t) \le m_t$ and $\operatorname{rank}(\tilde{K}_b) \le m_b$. The key advantage is that when the feature count is smaller than the number of genes, i.e., $m := m_t + m_b < G$, we avoid eigendecomposing the dense $G \times G$ matrices: we instead work in the $m$-dimensional feature space and map the eigenvectors back. When $m \ge G$ the dense $G \times G$ computation is already optimal, and the algorithm uses it instead.

The two contrastive methods differ in the matrix whose top eigenvectors we want:

$$\text{Kernel cPCA:} \quad \Sigma(\alpha) = \tilde{K}_t - \alpha\, \tilde{K}_b,$$
$$\text{Kernel ContraPC:} \quad \Sigma(\alpha) = (\tilde{K}_b + \gamma I)^{-\alpha}\, \tilde{K}_t.$$

##### 3.3 Kernel cPCA

Kernel cPCA seeks the top eigenvectors of the $G \times G$ symmetric matrix

$$\Sigma(\alpha) \;=\; \tilde{K}_t - \alpha\, \tilde{K}_b \;=\; M_t^\top M_t - \alpha\, M_b^\top M_b.$$

Consider any vector $v \in \mathbb{R}^G$ orthogonal to every feature row, i.e. $M_t v = 0$ and $M_b v = 0$. Then

$$\Sigma(\alpha)v \;=\; M_t^\top (M_t v) - \alpha\, M_b^\top (M_b v) \;=\; 0.$$

So every such $v$ is an eigenvector with eigenvalue $0$; the entire orthogonal complement of the combined row space collapses to zero. Consequently, every eigenvector with a nonzero eigenvalue must lie in

$$\mathcal{R} \;=\; \operatorname{range}\left( \begin{bmatrix} M_t \\ M_b \end{bmatrix}^\top \right), \qquad r := \dim \mathcal{R} \le m_t + m_b = m.$$

When $m < G$, $\mathcal{R}$ is a much smaller $r$-dimensional subspace of $\mathbb{R}^G$. The eigenvectors we use downstream are always the ones corresponding to the largest non-zero eigenvalues.

Let $Q \in \mathbb{R}^{G \times r}$ be an orthonormal basis of $\mathcal{R}$ ($Q^\top Q = I_r$), and write an eigenvector of interest as $v = Qc$. The subspace $\mathcal{R}$ is invariant under $\Sigma(\alpha)$: for any $v \in \mathcal{R}$, $\Sigma(\alpha)v \in \mathcal{R}$. Therefore, $\Sigma(\alpha)v$ remains the same after projection to $\mathcal{R}$, that is, $QQ^\top \Sigma(\alpha)v = \Sigma(\alpha)v$. Let

$$\hat{\Sigma}(\alpha) = Q^\top \Sigma(\alpha) Q \in \mathbb{R}^{r \times r}.$$

Left-multiplying $\Sigma(\alpha)v = \lambda v$ by $Q^\top$ gives $Q^\top \Sigma(\alpha) Q\, c = \lambda\, Q^\top Q\, c = \lambda c$, so each eigenpair $(\lambda, c)$ of $\hat{\Sigma}(\alpha)$ corresponds one-to-one with a nonzero eigenpair $(\lambda,\, v = Qc)$ of $\Sigma(\alpha)$.

Expanding $\hat{\Sigma}(\alpha)$,

$$\hat{\Sigma}(\alpha) = (M_tQ)^\top(M_tQ) - \alpha\,(M_bQ)^\top(M_bQ) = \underbrace{P_t^\top P_t}_{G_t} - \alpha\,\underbrace{P_b^\top P_b}_{G_b}, \qquad P_t = M_tQ,\ P_b = M_bQ.$$

Crucially, $G_t, G_b \in \mathbb{R}^{r\times r}$ do not depend on $\alpha$. They are computed once in the algorithm, so the entire per-$\alpha$ cost is

$$\hat{\Sigma}(\alpha) = G_t - \alpha\, G_b\ (O(r^2)), \quad \text{Eigen-decomposition of } \hat{\Sigma}(\alpha)\ (O(r^3)), \quad v = Qc \text{ (lift back to } \mathbb{R}^G).$$

$Q$ is obtained from a thin SVD of $A = \begin{bmatrix} M_t \\ M_b \end{bmatrix} \in \mathbb{R}^{m\times G}$, $A = UDV^\top$: the right singular vectors with nonzero singular values form an orthonormal basis of $\mathrm{range}(A^\top) = \mathcal{R}$. This costs $O(m^2G)$ and is done once.

##### 3.4 Kernel ContraPC

For $\alpha \neq 0$, the kernel ContraPC builds

$$\Sigma(\alpha) = (\tilde{K}_b + \gamma I)^{-\alpha} \cdot \tilde{K}_t.$$

A thin SVD of $M_b$ yields the background kernel matrix,

$$\tilde{K}_b \;=\; P\ \mathrm{diag}(\mu_i)\, P^\top, \qquad P \in \mathbb{R}^{G\times r_b},\ \ \mu_i = \sigma_i^2.$$

Using the eigendecomposition of $\tilde{K}_b$, we obtain

$$W(\alpha) := (\tilde{K}_b + \gamma I)^{-\alpha} \;=\; \gamma^{-\alpha} I \;+\; P\ \mathrm{diag}\big((\mu_i+\gamma)^{-\alpha} - \gamma^{-\alpha}\big)\, P^\top.$$

$P$ and $\mu$ come from an $O(m_b^2G)$ SVD of $M_b$, replacing the original $O(G^3)$ eigen-decomposition of a dense $G \times G$ matrix, $\tilde{K}_b + \gamma I$.

We note that every eigenvector with nonzero eigenvalue lies in the range. So we restrict $\Sigma(\alpha)$ to its own range. Since $W(\alpha)$ is invertible,

$$\mathrm{range}(\Sigma(\alpha)) = W(\alpha)\cdot \mathrm{range}(\tilde{K}_t) = W(\alpha)\cdot \mathrm{range}({M_t}^\top).$$

Let $Q_t \in \mathbb{R}^{G\times r_t}$ be an orthonormal basis of $\mathrm{range}(M_t^\top)$. By QR factorization of $W(\alpha)Q_t$, we obtain an orthonormal basis $Z \in \mathbb{R}^{G\times r_t}$ of $\mathrm{range}(\Sigma(\alpha))$. Consider

$$\hat{\Sigma}(\alpha) \;=\; Z^\top\Sigma(\alpha)Z \;=\; Z^\top\, W(\alpha)\,(M_t^\top M_t)\, Z \in \mathbb{R}^{r_t\times r_t}.$$

The nonzero eigenpairs of this small (non-symmetric) $\hat{\Sigma}(\alpha)$ coincide exactly with those of $\Sigma(\alpha)$, and we lift $v = Zc$. Every product is thin: $\tilde{K}_tZ = M_t^\top(M_tZ)$ and each application of $W(\alpha)$.

When $\alpha = 0$, $\Sigma(\alpha) = \tilde{K}_t$ is itself symmetric positive semi-definite, so we reduce on $Q_t$ and conduct eigen-decomposition.

$$\hat{\Sigma}(\alpha) \;=\; (M_tQ_t)^\top(M_tQ_t) \;=\; Q_t^\top \tilde{K}_t\, Q_t, \qquad v = Q_tc, \qquad \text{eigen-decomposition of } \hat{\Sigma}(\alpha).$$

#### 4 Comparison of atlas-wide meta-programs with restricted program libraries

We compared four feature-selection strategies of increasing stringency for constructing the atlas-wide meta-program from the perturbation-program library: (i) selecting all significant features from the full library; (ii) selecting a reproducibility-filtered subset, retaining only features that were significant in all 22 leave-one-chromosome-out (LOCO) sets; (iii) further restricting the reproducibility-filtered subset to weakly correlated features ($|r| < 0.8$); and (iv) further restricting the subset from (iii) to the most significant features, ranked by p-value, with the number of retained features matched to that selected by PoPS trained on the matched gene universe.

Performance was comparable across all four strategies in both the *in vitro* and *in vivo* Perturb-seq datasets (**Figure S12a**-**b**). These results indicate that the atlas-wide results are robust to the choice of feature-selection strategy and are not driven by redundancy among highly correlated programs.

#### 5 Tissue-level ENCODE-rE2G maps for variant-to-gene mapping

To link variants to representative genes in the perturbation meta-programs, we used tissue-level ENCODE-rE2G maps, defined as the union of regulatory element–gene links across biosamples from the corresponding broad tissue context (**Table S4**). Exact cell-line-specific rE2G maps frequently produced annotations too sparse for stable genome-wide S-LDSC estimation: when using K562-, Jurkat- and HepG2-specific links, all perturbation-derived annotations contained no more than 0.1% of common variants. We therefore used unions of links across matched hematopoietic, immune, hepatic or vascular biosamples, preserving broad tissue context while providing sufficient regulatory coverage. This strategy is consistent with previous approaches that combine cell-type-resolved gene programs with broader tissue-level regulatory maps for heritability analysis [4], and may be particularly appropriate for transformed cell lines whose regulatory landscapes can differ from those of primary healthy cells.
